# Characterization of a thaumarchaeal symbiont that drives incomplete nitrification in the tropical sponge *Ianthella basta*

**DOI:** 10.1101/527234

**Authors:** Florian U. Moeller, Nicole S. Webster, Craig W. Herbold, Faris Behnam, Daryl Domman, Mads Albertsen, Maria Mooshammer, Stephanie Markert, Dmitrij Turaev, Dörte Becher, Thomas Rattei, Thomas Schweder, Andreas Richter, Margarete Watzka, Per Halkjaer Nielsen, Michael Wagner

**Affiliations:** Division of Microbial Ecology, Department of Microbiology and Ecosystem Science, University of Vienna, Austria.; Australian Institute of Marine Science, Townsville, Queensland, Australia.; Australian Centre for Ecogenomics, School of Chemistry and Molecular Biosciences, University of Queensland, St Lucia, QLD, Australia; Center for Microbial Communities, Department of Chemistry and Bioscience, Aalborg University, 9220 Aalborg, Denmark.; Institute of Marine Biotechnology e.V., Greifswald, Germany; Division of Computational Systems Biology, Department of Microbiology and Ecosystem Science, University of Vienna, Austria.; Institute of Microbiology, Microbial Proteomics, University of Greifswald, Greifswald, Germany; Institute of Pharmacy, Pharmaceutical Biotechnology, University of Greifswald, Greifswald, Germany; Division of Terrestrial Ecosystem Research, Department of Microbiology and Ecosystem Science, University of Vienna, Austria.

## Abstract

Marine sponges represent one of the few eukaryotic groups that frequently harbor symbiotic members of the *Thaumarchaeota*, which are important chemoautotrophic ammonia-oxidizers in many environments. However, in most studies, direct demonstration of ammonia-oxidation by these archaea within sponges is lacking, and little is known about sponge-specific adaptations of ammonia-oxidizing archaea (AOA). Here, we characterized the thaumarchaeal symbiont of the marine sponge *Ianthella basta* using metaproteogenomics, fluorescence *in situ* hybridization, qPCR and isotope-based functional assays. “*Candidatus* Nitrosospongia bastadiensis” is only distantly related to cultured AOA. It is an abundant symbiont that is solely responsible for nitrite formation from ammonia in *I. basta* that surprisingly does not harbor nitrite-oxidizing microbes. Furthermore, this AOA is equipped with an expanded set of extracellular subtilisin-like proteases, a metalloprotease unique among archaea, as well as a putative branched-chain amino acid ABC transporter. This repertoire is strongly indicative of a mixotrophic lifestyle and is (with slight variations) also found in other sponge-associated, but not in free-living AOA. We predict that this feature as well as an expanded and unique set of secreted serpins (protease inhibitors), a unique array of eukaryotic-like proteins, and a DNA-phosporothioation system, represent important adaptations of AOA to life within these ancient filter-feeding animals.

**Originality-Significance Statement:** Many marine sponges harbor symbiotic members of the *Thaumarchaeota*, but there is generally only indirect evidence available about their functional role within these filter-feeding animals. Furthermore, the specific adaptations of thaumarchaeal symbionts to their sponge hosts are incompletely understood. In this study, we thoroughly characterized a thaumarchaeal symbiont residing in the reef sponge *Ianthella basta* and demonstrate by using a combination of molecular tools and isotope techniques, that it is the only ammonia-oxidizer in its host. In contrast to other sponges, *I. basta* does not contain nitrite-oxidizing microbes and thus excretes considerable amounts of nitrite. Furthermore, using metagenomics and metaproteomics we reveal important adaptations of this symbiont, that represents a new genus within the *Thaumarchaeota*, and conclude that it most likely lives as a mixotroph in its sponge host.

## Introduction

Marine sponges (phylum *Porifera*) are among the most basal metazoan lineages (Simion *et al*., 2017), with fossil records suggesting their evolutionary emergence more than 600 million years ago (Love *et al*., 2009). Sponges form a major part of the marine benthic fauna across the world’s oceans (Bell, 2008) where they mediate critical biogeochemical processes through their filtration of immense volumes of seawater (Maldonado *et al*., 2012; de Goeij *et al*., 2013). Many sponges harbor dense, diverse and species-specific communities of microbes (Hentschel *et al*., 2012; Thomas *et al*., 2016), and these associations are often temporally and geographically stable (Luter *et al.*, 2010; Schmitt *et al*., 2012; Astudillo-García *et al.*, 2017). Functional roles that have been assigned to specific sponge-associated microorganisms include the provision of photosynthates (Wilkinson, 1983) and fixed N_2_ (Wilkinson, 1979) by cyanobacterial symbionts and the production of bioactive secondary metabolites (Wilson *et al*., 2014; Freeman *et al*., 2016; Agarwal *et al*., 2017). Furthermore, sponge symbionts can serve as an endogenous food source as exemplified by biomass transfer from a sponge-associated sulfate-reducing bacterial community to host cells (Hoffmann *et al*., 2005), and the ingestion of symbiotic methanotrophs in a deep-sea carnivorous sponge (Vacelet *et al*., 1995). While a vast array of additional putative symbiotic functions have been hypothesized from taxonomic or metagenomic data (Webster and Thomas, 2016), unequivocal evidence for specific sponge symbiont physiologies is comparably rare.

Nitrogen cycling in sponge holobionts has received considerable attention and sponge-symbiont-driven nitrification, denitrification, and anaerobic ammonium oxidation has been described (Bayer *et al*., 2008; Southwell *et al*., 2008; Hoffmann *et al*., 2009; Schläppy *et al*., 2010; Radax *et al*.,2012a). Many marine sponges harbor symbionts phylogenetically related to nitrifying microbes (Steger *et al*., 2008; Hoffman *et al*., 2009; Off *et al*., 2010), indicating that ammonia oxidation via nitrite to nitrate is a widely distributed process in these animals. More specifically, molecular signatures of proteobacterial as well as thaumarchaeal ammonia oxidizers are frequently detected in sponges, although these microbes rarely co-occur in the same host (Bayer *et al*., 2008; Radax *et al*., 2012b). Most sponges that contain symbionts related to ammonia-oxidizers also host bacteria affiliated with known nitrite-oxidizers, particularly members of the genus *Nitrospira* (Hoffmann *et al*., 2009; Off *et al*., 2010; Reveillaud *et al*., 2014; Moitinho-Silva *et al*., 2017a). While many studies simply equate the molecular detection of microbes related to recognized nitrifiers with the occurrence of canonical nitrification in sponges (Fan *et al*., 2012; Mohamed *et al*., 2010), this assumption could be misleading due to the functional versatility of both *Thaumarchaeota* and *Nitrospira* (Mussmann *et al*., 2011; Koch *et al*., 2014; Daims *et al*., 2015; Palatinszky *et al*., 2015).

Thaumarchaeotes are important ammonia oxidizers in many environments (Pester *et al*., 2011). Marine sponges host a particularly high diversity of thaumarchaeotes, with many of the 16S rRNA gene sequences falling into phylogenetic clusters that exclusively contain sponge-derived sequences (Simister *et al*., 2012). The detection of thaumarchaeotes in sponge larvae also suggests vertical transmission or early environmental acquisition of these symbionts (Sharp *et al*., 2007; Steger *et al*., 2008; Schmitt *et al*., 2012). The first genomic information of a member of this phylum was derived from *Ca.* Cenarchaeum symbiosum in the marine sponge *Axinella mexicana* (Hallam *et al*., 2006), and subsequently thaumarchaeal genomes were recovered from the glass sponge *Lophophysema eversa* (Tian *et al*., 2016) and the temperate sponge *Cymbastella concentrica*, with the latter also shown to transcribe genes for ammonia oxidation (referred to as CCThau in Moitinho Silva *et al.*, 2017a and in this manuscript). However, little information is available on the genomic plasticity and mechanisms of host adaptation in sponge thaumarchaeal symbionts. Furthermore, with the exception of a thaumarchaeal symbiont that inhabits the cold-water sponge *Geodia barretti* (Radax *et al*., 2012a, b), direct evidence for the catalysis of ammonia oxidation by sponge-associated thaumarchaeotes is lacking. While transcription and translation of key functional genes like the *amoA* gene encoding a subunit of the ammonia monooxygenase of thaumarchaeotes has been detected across multiple sponge species (Liu *et al*., 2012; Fiore *et al*., 2015; Moitinho-Silva *et al*., 2017a), experimental validation of their involvement in nitrification is required to confirm this activity *in situ* (Mussmann *et al*., 2011).

To better understand the physiological capability of Thaumarchaeota in sponges, we used a metaproteogenomic approach to characterize the Thaumarchaeota symbiont of the marine sponge *Ianthella basta*. *I. basta* is an abundant and ecologically important reef sponge found throughout the Indo-Pacific (Berquist and Kelly, 1995). In contrast to many microbially diverse sponge species, 16S rRNA gene surveys have shown that *I. basta* harbors only three dominant microbial phylotypes which cluster within the α-and γ-proteobacteria as well as the *Thaumarchaeota* (Luter *et al*., 2010). This community structure is stable among different host color morphotypes (Freckelton *et al*., 2012), between individuals sampled from different geographic regions (Luter *et al*., 2010), across different host health states (Luter *et al*., 2010) and in sponges exposed to different environmental stressors (Luter *et al*., 2012). Here we (i) quantify the abundance of the thaumarchaeal symbionts in *I. basta*, (ii) confirm their role as ammonia oxidizers (iii) document the expression of almost 100 thaumarchaeal genes *in situ*, and (iv) reveal sponge-specific adaptations of these archaea including a putative mixotrophic lifestyle. Furthermore, we demonstrate that nitrite oxidation surprisingly does not occur in *I. basta*.

## Results and Discussion

### Quantification of the thaumarchaeal *I. basta* symbiont using FISH and qPCR

FISH was performed on fixed cryosections from one sponge individual using the general archaeal probe Arch915 that is fully complementary to the 16S rRNA of the single archaeal phylotype known to inhabit *I. basta* (Fig. 1). Quantitative FISH across 10 images revealed that the thaumarchaeal symbiont comprised 24 ± 1.6% (standard error, SE) of the total bacterial and archaeal cells detected with probes Arch915 and the probe set EUB338-I-III targeting most bacteria. Absolute quantification of the thaumarchaeal symbiont in five sponge individuals (including those used for metagenome sequencing) using specifically designed qPCR primers, revealed an average absolute abundance of 2.41 ± 0.7 (SE) × 10^10^ 16S rRNA gene copies per g wet weight of sponge tissue. Consistent with the relative abundances derived from FISH, the ratio of thaumarchaeal 16S rRNA sequences to the total 16S rRNA genes derived from qPCR assays targeting the additional α-and γ-proteobacterial symbionts of *I. basta* (data not shown) was 22 ± 2.3 (SE)%. Collectively, these data demonstrate that the *I. basta* thaumarchaeote is a dominant member of the *I. basta* microbiome, occurring at densities that exceed the total microbial biomass of many tropical sponge species (Taylor *et al*., 2007).

**Figure 1.**
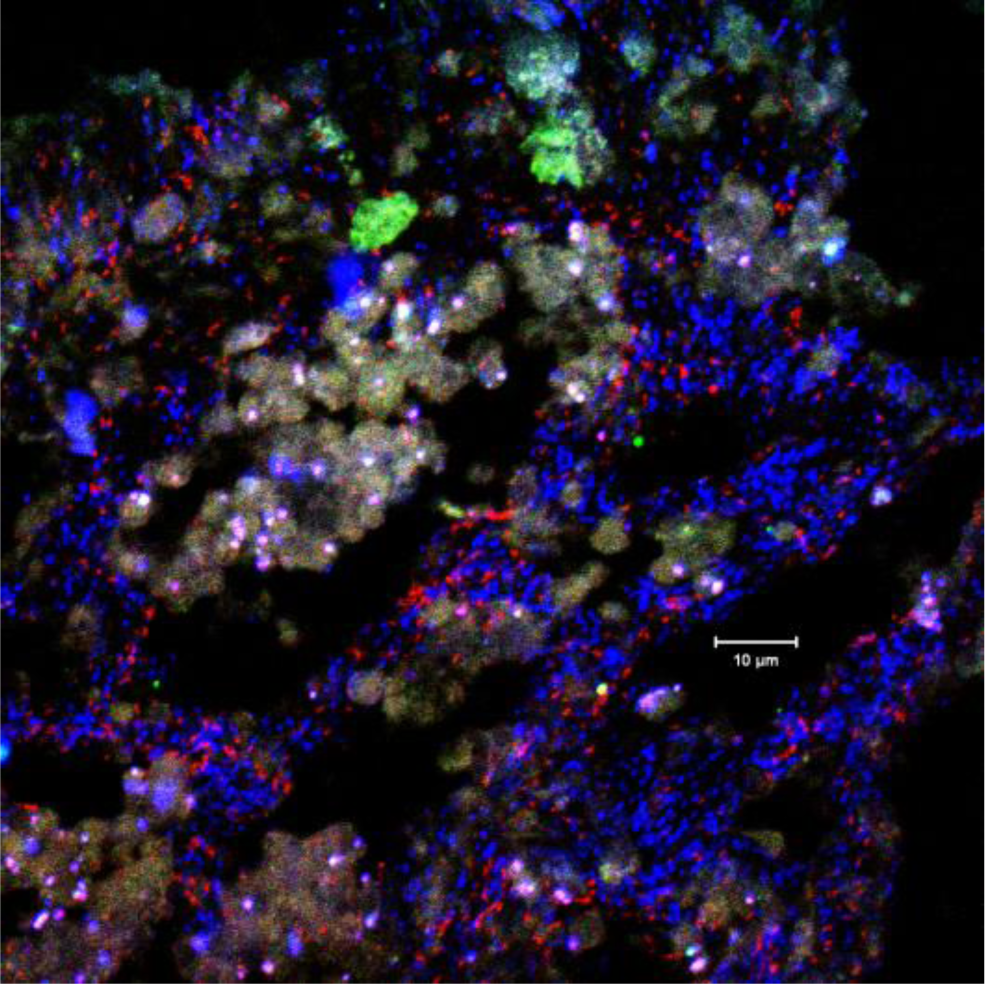
Fluorescence *in situ* hybridization of a 5 μm cryosection of *Ianthella basta* using double-labeled (Stoecker *et al.*, 2010) probe Arch915 in red and the double-labeled probe EUB338-I-III set in blue. Green and white/lila structures represent autofluorescence. As *I. basta* harbors “*Ca.* Nitrosospongia bastadiensis” as the only archaeon, all red signals represent this AOA.

### Metaproteogenomic analyses of the thaumarchaeal symbiont of *I. basta*

An extensive metagenomic data set consisting of 25.8 Gbp of sequence information was obtained for an *I. basta* individual by Illumina sequencing. A 1.99 Mbp Metagenome-Assembled Genome (MAG) consisting of 113 contigs was recovered, representing a nearly complete thaumarchaeal genome (99%) with very low contamination (Supporting Information Table S1). An additional unpublished metagenomic data set (250 Mbp generated from pyrosequencing) derived from a different *I. basta* individual was also screened to confirm the presence of genes of interest (Supporting Information Table S2) and support the phylogenetic inferences displayed in Fig. 2. While the shallow metagenome was excluded from detailed analyses, it confirmed that closely related thaumarchaeal symbionts inhabited both sponge individuals. The average nucleotide identity (ANI) between the two thaumarchaeote MAGs was 98.2%, with 82% coverage of the smaller (Illumina data set) MAG, confirming that members of the same thaumarchaeal species (Konstantinidis *et al*., 2017) reside in both sponge individuals. Both thaumarchaeal symbiont MAGs from *I. basta* possess the highest GC content (64.8%) of any genome-sequenced thaumarchaeote (Fig. 2C) (see Supporting Information for a more detailed discussion of the high GC content).

**Figure 2.**
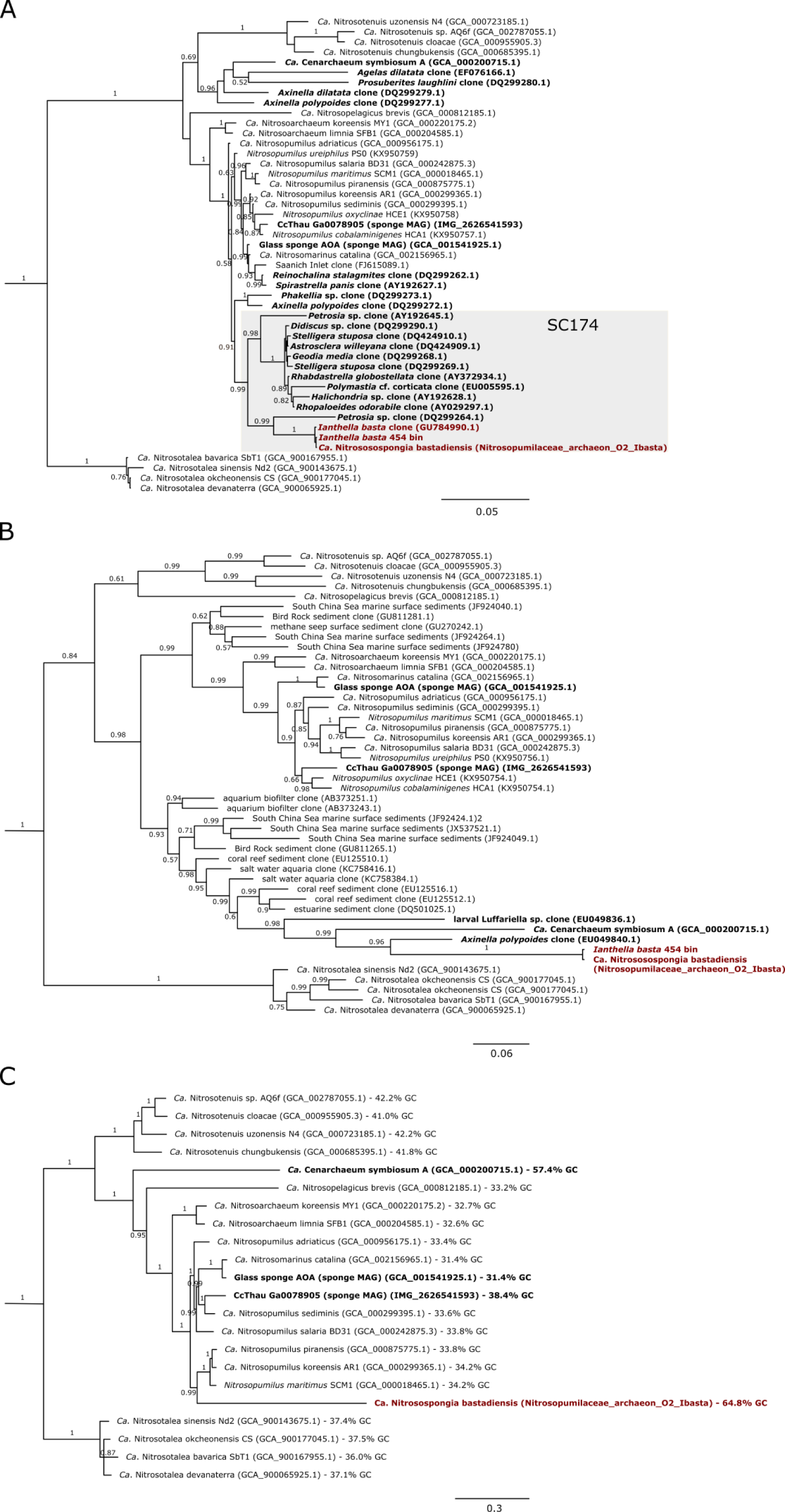
Phylogeny of *Ca.* Nitrosospongia bastadiensis. (A) Bayesian 16S rRNA gene tree. (B) Bayesian *amoA* gene tree. (C) Bayesian phylogenomic tree. For this analysis, *Ca*. N. bastadiensis and *Ca.* C. symbiosum *amoA* sequences were placed into a reference tree, representing all OTU representatives of the curated database provided in Alves *et al.* (2018), using the Evolutionary Placement Algorithm (EPA; Berger *et al*., 2011) implemented in RAxML-HPC 8.2.11 (Stamatakis, 2014). Representative sequences clustering with the sponge symbiont *amoA* sequences were then analyzed along with *amoA* sequences from the other genome-sequenced AOA. Note that *Ca.* Nitrosotalea okcheonensis possesses two *amoA* gene copies (Herbold *et al*., 2017). (C) Bayesian phylogenomic tree based on 34 concatenated universal, single-copy marker genes identified with CheckM (Parks *et al*., 2015). Bayesian posterior support values >0.5 are indicated for each branch. Outgroups for all trees consisted of all three genome-sequenced members of the *Nitrososphaera* cluster, both members of the *Nitrosocosmicus* clade, and *Ca.* Nitrosocaldus icelandicus. In all trees, sequences obtained from sponges are depicted in bold.

Metaproteomic analysis performed on the same *I. basta* individual used for Illumina metagenomic sequencing detected a total of 513 proteins, 96 of which were specifically assigned to the *I. basta* thaumarchaeote (representing 5.4% of the genes in the MAG). When combining the normalized spectral abundance factor (NSAF; Florens *et al*., 2006) values for all samples and analyses, the 96 thaumarchaeal proteins comprised 10.5% of the 513 identified proteins (Supporting Information Table S3). Of the thaumarchaeal proteins, 60.2% (67.4% NSAF) were encoded by gene families shared by all thaumarchaeotes, and 7.1% (3.1% NSAF) were encoded by genes unique to the *I. basta* thaumarchaeote. Of the 96 expressed proteins, 88 were also encoded at a predicted average amino acid identity of 99.4% ± 1.4 (SD) in the second MAG recovered by pyrosequencing. The other 8 protein homologues were also found in the second MAG but at much lower predicted amino acid identities (from 31.4 to 89%).

### *I. basta* contains a symbiont from a new thaumarchaeal genus

Of all genome-sequenced thaumarchaeotes, the *I. basta* symbiont has the highest average genomic amino acid identity (AAI) of 58.1% with *Ca.* Nitrosopumilus piranensis. As this represents a new species within a new genus (60-80% AAI is typical for organisms grouped at the genus level; Luo *et al*., 2014; Konstantinidis *et al*., 2017), we propose the name *Ca.* Nitrosospongia bastadiensis. 16S rRNA gene, *amoA*, and concatenated marker gene phylogenies suggest that *Ca*. N. bastadiensis is a member of the family *Ca. Nitrosopumilaceae* (Qin *et al*., 2016) (Fig. 2). Based on the 16S rRNA gene sequence, *Ca.* N. bastadiensis is a member of the sponge-specific sequence cluster 174 (Simister *et al*., 2012) which does not include the three other sponge thaumarchaeal symbionts (*C. symbiosum,* CCThau, and a putative deep-sea glass sponge thaumarchaeal symbiont) for which genome sequences are available (Hallam *et al*., 2006; Tian *et al*., 2016; Moitinho-Silva *et al*., 2017a). The lack of a close relationship between *Ca.* N. bastadiensis and the other sponge-associated symbionts is consistent with the topology of the concatenated single copy conserved marker gene tree (Fig. 2C), while unexpectedly *amoA* phylogeny supported a clustering of *Ca.* N. bastadiensis with *C. symbiosum* (Fig. 2B). While the *amoA* genes of *C. symbiosum* and *Ca.* N. bastadiensis showed compositional bias, it was not possible to determine whether this caused their monophyletic grouping in the *amoA* gene tree. Upon individual addition of *C. symbiosum* and *Ca*. N. bastadiensis *amoA* gene sequences to an *amoA* data set, their phylogenetic position in the tree varied dependent on taxa selection (data not shown). Querying the *Ca*. N. bastadiensis 16S rRNA gene against the Sponge Microbiome Project (SMP) database (Moitinho-Silva *et al*., 2017b) and against most other publicly available 16S rRNA gene amplicon data sets demonstrated that the habitat of *Ca*. N. bastadiensis is restricted to a few sponge species (see Supporting Information for more details).

### Core metabolism of *Ca.* N. bastadiensis

No major difference in core metabolism was detected between *Ca.* N. bastadiensis and other AOA. Consistent with marine and non-marine thaumarchaeota (Hallam *et al*., 2006; Tourna *et al*., 2011; Spang *et al*., 2012; Park *et al*., 2014; Bayer *et al.*, 2016), *Ca.* N. bastadiensis possesses a complete urease gene cluster in addition to a gene encoding a urea active transporter (*DUR3*; 86% homologous to *Ca.* Nitrosopumilus piranensis) and is thus capable of taking up (or using internally produced) urea and converting it to ammonia and CO_2_. The sponge symbionts *C. symbiosum* and CCThau also encode urea transporters and urease gene clusters. Furthermore, urease subunit gene transcripts from CCThau were found in the metatranscriptomes from *C. concentrica* (Moitinho-Silva *et al*., 2017a). Thus, it seems likely that urea is not only used by marine AOA in Arctic waters (Alonso-Sáez *et al*., 2012), but is also a common substrate for group I.1a *Thaumarchaeota* residing in marine sponges and likely represents an important adaptation to life within their hosts that might be excreting urea (Morley *et al.*, 2016). In contrast to *Nitrososphaera gargensis* (Palatinszky *et al.*, 2015), *Ca.* N. bastadiensis does not contain a cyanase for ammonia generation from cyanate, but like all other AOA, it encodes a protein with modest homology to a creatinine-amidohydrolase indicating that it could utilize sponge-derived creatinine and convert it to creatine. While all genome-sequenced AOA symbionts lack a canonical creatinase that would form urea from creatine, AOA including *Ca.* N. bastadiensis do possess a Xaa-Pro aminopeptidase that has been hypothesized as a functional analog (Moitinho-Silva *et al*., 2017a). All genes encoding the putative subunits of the ammonia monooxygenase (*amo*) enzyme were also found in the *I. basta* thaumarchaeote (*amoA*, *amoB*, and *amoC*), including the hypothetical gene *amoX. AmoB* and *C* were also detected as proteins (Supporting Information Table S3). The *amo* gene arrangement [*amoA*-*amoX*-*amoC*-*amoB*] is syntenic to most analyzed members of *Ca*. Nitrosopumilaceae (Lehtovirta-Morley *et al*., 2011; Park *et al*., 2014). Consistent with other AOAs, no canonical hydroxylamine dehydrogenase was found but *Ca*. N. bastadiensis did encode lineage 1 multicopper oxidases (MCO), which have been identified as candidates for archaeal hydroxylamine dehydrogenases (Kerou *et al*., 2016). Like most other AOA (with the exception of *Ca.* C. symbiosum), *Ca*. N. bastadiensis encodes the putative NO-forming nitrite reductase (*nirK*; found to be highly expressed as protein; Supporting Information Table S3). Interestingly however, the purple cupredoxin Nmar_1307 from *Nitrosopumilus maritimus* that is capable of oxidizing NO to NO_2_-(Hosseinzadeh *et al*., 2016) is absent in *Ca*. N. bastadiensis and other sponge-associated AOA (and also many other AOA), but there are several mononuclear cupredoxins encoded by *Ca*. N. bastadiensis that could have the same function. NO has been postulated as a key intermediate of AOA (Kozlowski *et al*., 2016; Carini *et al*., 2018), a hypothesis that nicely explains the inhibitory effect of the NO scavenger PTIO used in our incubation experiments described below. Consistent with all other AOA, no canonical NO-or N_2_O-reductases were found, although - like in many other thaumarchaeotes (Liu *et al*., 2012, Zhalnina *et al*., 2014; Santoro *et al*., 2015) - the putative nitric oxide reductase accessory proteins, NorQ and NorD are encoded in the genome (with the NorD subunit being expressed).

Intriguingly, *Ca*. N. bastadiensis encodes many putative surface-layer (S-layer) proteins (Supporting Information Fig. S1), with six of them being expressed at combined NSAF values of 6.78% (Supporting Information Table S3). The expanded group of S-layer proteins in *Ca*. N. bastadiensis (and *Ca.* C. symbiosum) may be involved in molecular sieving, adhesion, surface recognition or other types of interactions within the extracellular matrix of the sponge environment. The negatively charged surface of the Thaumarchaeota S-layer was recently proposed to help concentrate the charged solute ammonium into the pseudo-periplasmic space (Li *et al*., 2018). The theoretical isoelectric points of all *Ca.* N bastadiensis S-layer proteins (3.62 – 4.92) were in a similar range to those calculated for *N. maritimus* and *Nitrosoarchaeum limnia* SFB1 S-layer proteins (3.4 – 4.08), consistent with the proposed mechanism for charged solute acquisition.

As expected, *Ca.* N. bastadiensis encodes all key enzymes of the thaumarchaeal 3-hydroxypropionate/4-hydroxybutyrate pathway (Könneke *et al*., 2014; Otte *et al*., 2015) and three proteins catalyzing five steps of this thaumarchaeal CO_2_ fixation pathway were also detected within the metaproteome. Regarding nitrogen assimilation, *Ca.* N. bastadiensis encodes and expresses an *amt* transporter of the *amt-2* lineage which has been hypothesized to be a high affinity ammonia transporter (Nakagawa and Stahl, 2013; Offre *et al*., 2014). The *amt-2* gene of *Ca.* N. bastadiensis is nested within the complex V ATP synthase operon (of which four of the encoded subunits were found to be expressed), indicating that energy generated from ammonia oxidation directly fuels the transport of ammonium for assimilation. Interestingly, in contrast to most other AOA, no putative low affinity *amt* transporter gene was found in *Ca.* N. bastadiensis, a feature shared with the sponge symbionts *Ca.* C. symbiosum (Hallam *et al*., 2006) and CCThau (Fig. 3; Moitinho-Silva *et al*., 2017a). This pattern indicates that sponge thaumarchaeotes are either not exposed to high concentrations of ammonia or that they can assimilate nitrogen under most conditions by uptake of organic nitrogen sources and only use the high affinity ammonia transporter if organic nitrogen sources are scarce. Regarding nitrogen assimilation, it is also noteworthy, that the almost complete bin of *Ca.* N. bastadiensis contains only a single gene encoding a member of the nitrogen regulatory protein PII superfamily, whereas, with the exception of *Ca.* C. symbiosum, higher copy numbers of these genes are generally found in other AOA (Fig. 3; Kerou *et al*., 2016). Furthermore, in contrast to most other AOA, the two key genes coding for subunits of the polyhydroxyalkanoate (PHA) synthase *phaC* and *phaE* are apparently lacking in *Ca.* N. bastadiensis. We also found no indications for use of alternative carbon storage compounds such as starch or glycogen in *Ca.* N. bastadiensis.

**Figure 3.**
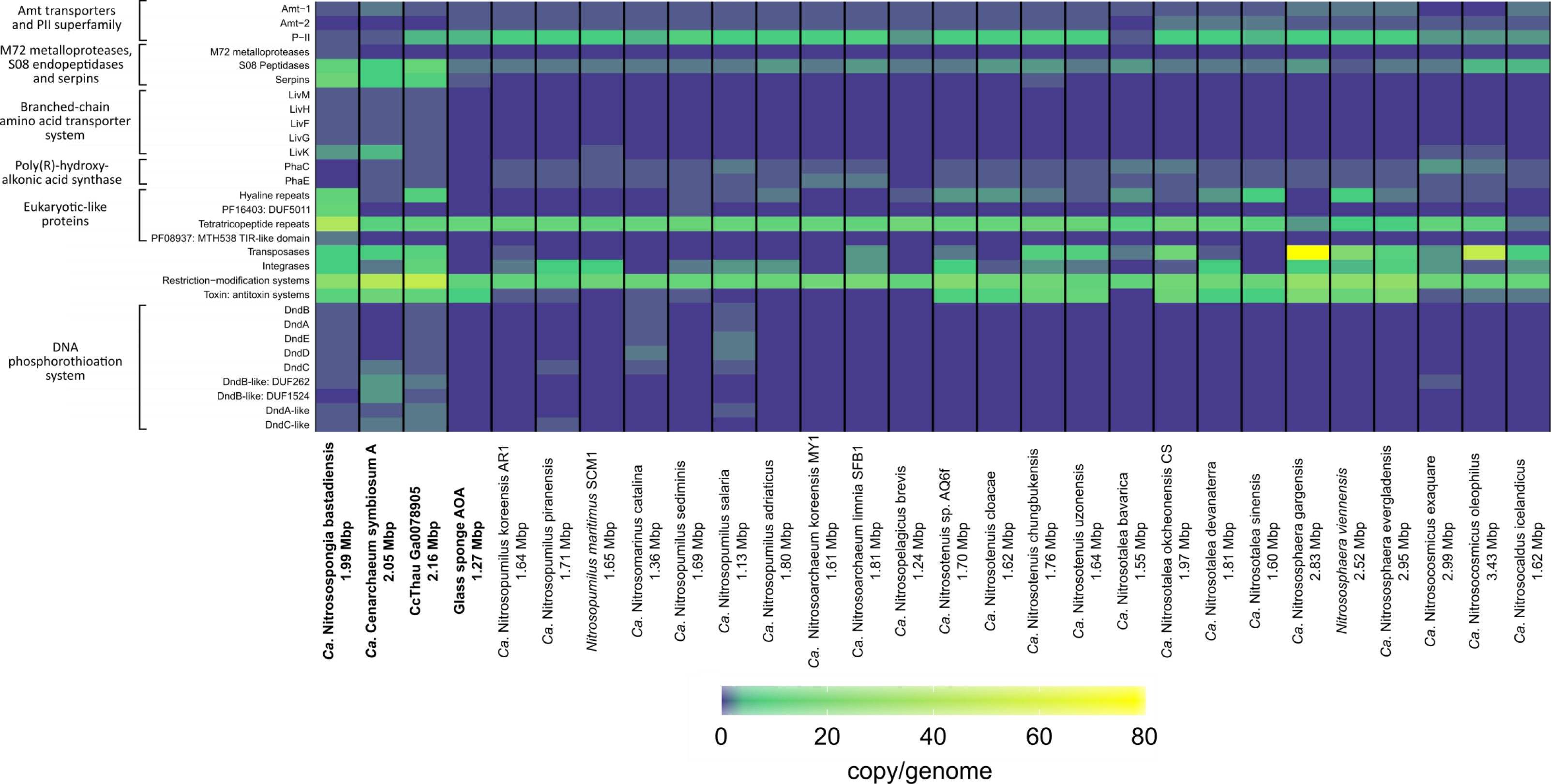
Heat map showing the distribution and gene copy number per genome of selected genes and gene classes among genome-sequenced AOA. The color scale indicates copies per genome and MAG, respectively. Sponge-derived MAGs start on the left and are depicted in bold, followed by members of *Ca*. Nitrosopumilaceae, *Ca*. Nitrosotenuaceae, *Ca*. Nitrosotaleale, the *Nitrososphaerales*, and *Ca*. Nitrosocaldales. Genome sizes are listed next to each species name. An extended version of this Figure is available as Supplemental Information Fig. 5.

In contrast to many free-living AOA (Blainey *et al*., 2011; Jung *et al.*, 2014; Mosier *et al*., 2012; Spang *et al*., 2012; Bayer *et al*., 2016) and the glass-sponge AOA, *Ca.* N. bastadiensis, CCThau, and *Ca* C. symbiosum do not encode an archaellum. Lack of chemotaxis and motility traits has been described for other microbial symbionts (Karimi *et al*., 2018) and likely represents an adaptation to the sponge-associated lifestyle. The absence of an archaellum in *Ca.* N. bastadiensis would also be consistent with vertical transmission of this symbiont, although this remains to be demonstrated.

### Shared gene families of sponge AOA

During genome annotation of *Ca.* N. bastadiensis, we particularly focused on gene families unique to this organism or to *Ca.* N. bastadiensis and other thaumarchaeal sponge symbionts (among all genome-sequenced thaumarchaeotes) as these gene families likely represent adaptations to a sponge-associated lifestyle. While 40% (represented by 681 gene families) of the *Ca.* N. bastadiensis genes were members of core gene families represented in every sequenced thaumarchaeote, 35% (represented by 616 gene families) had no close orthologues among other thaumarchaeotes. Additional pairwise comparisons between *Ca.* N. bastadiensis and all other thaumarchaeote genomes were performed in order to identify shared gene families of species pairs that are absent in all other thaumarchaeotes. Interestingly, the highest number occurred with CCThau (n = 18) and *Ca.* C. symbiosum (n=11) (Supporting Information Fig. S2). Furthermore, 14 gene families were found to be exclusively shared among all three thaumarchaeal sponge symbionts, while only one gene family was found to be shared with the putative deep-sea glass sponge thaumarchaeal symbiont and *C. symbiosum*. In total, 44 gene families were exclusively shared with at least one other thaumarchaeal sponge symbiont representing 2.4% of the *Ca.* N. bastadiensis genes.

### *Ca.* N. bastadiensis and other sponge AOA secrete proteases and their inhibitors

Among the genes that were unique to *Ca.* N. bastadiensis or exclusively shared with other sponge AOA were several genes involved in degradation of extracellular protein and inactivation of extracellular (host-derived) proteases. *Ca.* N. bastadiensis possesses a putatively exported metalloprotease of the M72 family (Drapeau, 1980; Passmore *et al.*, 2015) that was also detected in the metaproteome. This large protease (2027 AA) contains hyaline repeat domains suggestive of an additional adhesive property. The M72 family of metalloendopeptidases have so far not been found in any *Archaea* (Trame *et al*., 2014) and most characterized members of this enzyme family are peptidyl-Asp metalloendopeptidases that hydrolyze bonds on the NH_2_-terminal side of aspartic acid and cysteic acid residues (Drapeau, 1980). Furthermore, the *I. basta* thaumarchaeal symbiont contained 11 genes affiliated with four gene families that encode subtilisin-like serine protease domains (S08A family endopeptidases) (Supporting Information Fig. 3A). Although serine endopeptidases of the S08A family are also found in other thaumarchaeotes, with the exception of *C. symbiosum* (4 copies), and CCThau (13 copies), most contain only two copies. Furthermore, it is noteworthy that two of the four S08A family endopeptidase gene families were either exclusively found in *Ca*. N. bastadiensis or shared with CCThau (Fig. 4, Supporting Information Fig. 3A). Interestingly, several of the *Ca.* N. bastadiensis S08A endopeptidases are predicted to be exported without a membrane anchor, while most other thaumarchaeal S08A serine endopeptidases seem to be membrane anchored.

**Figure 4.**
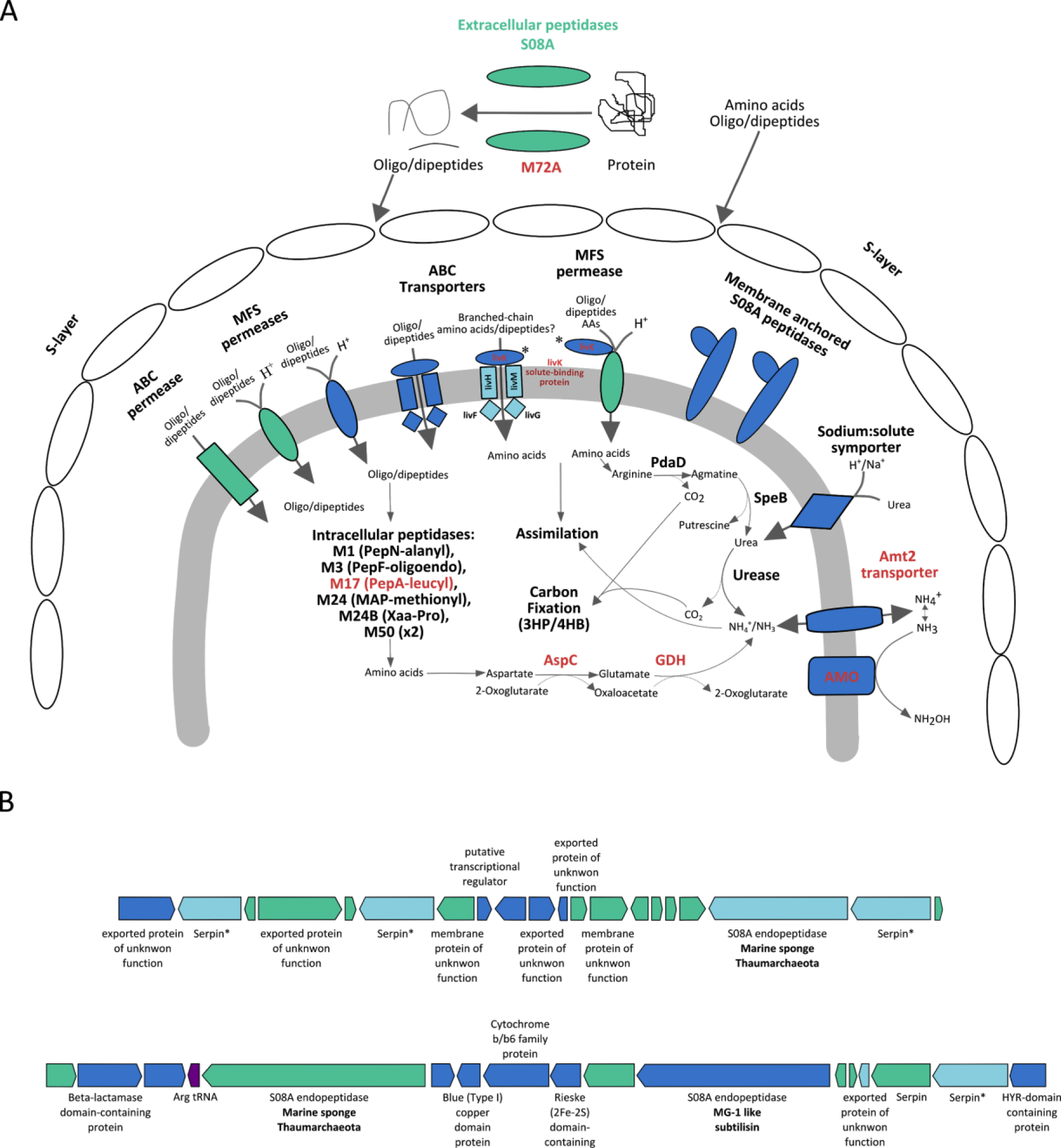
(A) Reconstructed metabolic pathways of genes and their expression (red) detected in *Ca.* N. bastadiensis proposed to be involved in extracellular (and intracellular) protein degradation as well as amino acid transport and assimilation. Predicted proteins (and their respective subunits when relevant) are color coded to denote the degree of homology among all sequenced *Thaumarchaeota*: green - *Ca*. N. bastadiensis unique gene families; light blue - shared exclusively among thaumarchaeal sponge symbionts; dark blue - ubiquitously found in *Thaumarchaeota*. Extracellular proteins derived from the marine environment as well as the sponge mesohyl may be degraded extracellularly or in the thaumarchaeal pseudo-periplasmic space. These resultant oligo/dipeptides and amino acids which can also be derived from the environment can then be transported by a suite of ABC transporters and MFS permeases into the cytoplasm to be further degraded by intracellular peptidases or assimilated. Amino acids such as arginine and aspartate can be further degraded to form NH_3_/NH_4_^+^ for assimilation or export to the pseudo-periplasmic space for ammonia oxidation. Arginine can be decarboxylated by arginine decarboxylase (PdaD) to agmatine which can then be degraded to urea by agmatinase (SpeB). Both proteins are ubiquitously distributed among the *Thaumarchaeota*. All but one of the branched-chain amino acid transporter subunits (LivFGHMK operon) are exclusively found among thaumarchaeal sponge symbionts, while the periplasmic solute binding subunit (LivK – found to be expressed and denoted by an asterisk) can be found not only among sponge symbionts but also in *N. maritimus* and members of the genus *Nitrosocosmicus* (see also Fig. 3). (B) Genetic context map depicting examples of co-localized S08A endopeptidases and serine protease inhibitors (serpins). Color coding as in panel (A), except purple denotes the presence of a tRNA, sites at which gene insertions are common. The asterisk next to serpins color coded as sponge-specific (light-blue) denotes that *Ca*. N. chungbukensis is the sole non-sponge symbiont encoding a serpin and belonging to this orthologous group (see Supplemental Information Fig. S3B). AspC, aspartate aminotransferase; GDH, glutamate dehydrogenase.

*Ca.* N. bastadiensis also encodes 15 genes for the I4 family of serpins which represent serine protease inhibitors. Some of these cluster with serpins from *C. symbiosum* (Hallam *et al*., 2006) and CCThau, although amongst the non-host associated AOA, only *Nitrosotenuis chungkubensis* encodes a serpin (Supporting Information Fig. 3B). Nine of the *Ca.* N. bastadiensis serpins are predicted to be extracellular and are frequently found adjacent to S08A family endopeptidases in the genome (Fig. 4B). Serpins belong to a large family of irreversible inhibitory substrates of proteases, often but not exclusively of the serine class, that has been widely characterized in mammals, insects, plants as well as some viruses, and which can mediate host-microbe interactions (Ventura *et al*., 2012). These findings suggest that AOA living as sponge symbionts use serpins to regulate their own secreted endopeptidases and/or sponge serine proteases found to be highly expressed in other sponges (Riesgo *et al*., 2014), including within the host of *C. symbiosum* (Zaikova, E., Ph.D. thesis, 2007).

### Peptide and amino acid uptake by *Ca.* N. bastadiensis and other sponge AOA

*Ca.* N. bastadiensis, like many other AOA, is well equipped for uptake of oligo-and dipeptides as well as amino acids by encoding a set of transporters widely distributed in this clade. After uptake, C*a.* N. bastadiensis, like other AOA, has the genomic repertoire to degrade these peptides and amino acids and release ammonia (Fig. 4). In addition to features common to many AOA, *Ca.* N. bastadiensis also contains the *livFGHMK* operon which in proteobacteria and cyanobacteria encodes a high-affinity branched-chain amino acid transport system (which can also transport other amino acids) belonging to the ATP binding cassette (ABC) superfamily of transporters (Hoshino, 1979; Adams *et al*., 1990; Hosie *et al*., 2002; Picossi *et al*., 2005) (Fig. 4). Among all other thaumarchaeote genomes, only *C. symbiosum*, CCThau, and a thaumarchaeote MAG from the Caspian Sea (Mehrshad *et al*., 2016) encode this transporter (Supporting Information Fig. S4). The periplasmic subunit LivK from *Ca.* N. bastadiensis was expressed as protein (Supporting Information Table S3) and this subunit was also reported to be expressed in CCThau (Moitinho-Silva *et al*., 2017a). Adjacent to the expressed *livK* in *Ca.* N. bastadiensis is a gene encoding three consecutive LivK domains (along with the requisite ligand binding sites) which is congruent with *C. symbiosum* containing 4 copies of the LivK solute binding component (Supporting Information Fig. S4A). In *C. symbiosum*, the transporter encoding the *livFGHMK* operon is located next to several extracellular trypsin-like serine protease encoding genes, indicating that the transporter is involved in amino acid uptake in this organism. However, given the substrate promiscuity of this transporter family (Adams *et al*., 1990; Valladares *et al*., 2002; Beckers *et al*., 2004; Picossi *et al*., 2005) (Supporting Information Fig. S4), experimental validation will be required before more specific predictions on its function in sponge AOA can be made. Such investigations would be particularly important, as multiple *LivFGHMK* operons are also present in two γ-proteobacterial sponge symbionts (Gauthier *et al*., 2016), suggesting that amino-acid transport and utilization mediated by this transporter-type could be an important feature of sponge symbionts and may even contribute to sponge-mediated dissolved organic matter transfer to higher trophic levels (de Goeij *et al*., 2013). Proteases are also important for sponge metabolism and a body of research has focused on sponge proteases and their inhibitors from a biodiscovery context (Arreguín *et al*., 1993; Wilkesman and Schroeder, 2002; Wilkesman and Schroeder, 2007; Tabares *et al*., 2011). In these analyses generally, sponge holobiont samples are used for enzyme or inhibitor purification and characterization. Thus, our observation that abundant archaeal sponge symbionts secrete proteases and serpins, demonstrate that in such assays, it will remain unclear whether it is the microbial symbionts, or the host animals that are producing the analyzed biomolecules.

A mixotrophic lifestyle for *Thaumarchaeota* has been inferred since the earliest reports on planktonic thaumarchaeotes described incorporation of organic carbon into signature lipids (Ingalls *et al*., 2006) and incorporation of labelled amino acids leucine (Ouverney and Fuhrman, 2000; Herndl *et al*., 2005) and aspartate (Teira *et al*., 2006) into single planktonic cells. While initial findings - that pure cultures of *Thaumarchaeota* in group I.1a (Qin *et al*., 2014) and group I.1b (Tourna *et al*., 2011) could have increased growth rates in the presence of α-keto acids and ammonia - were subsequently proven to be a result of the H_2_O_2_ scavenging ability of α-keto acids (Kim *et al*., 2016; Qin *et al*., 2017), abundant genes and transcripts of S08A family serine proteases and other metallopeptidases in MG-1 deep-sea thaumarchaea have also been attributed to mixotrophic activity (Li *et al*., 2015). A deep sea *Thaumarchaeota* has also been found to encode the *livFGHMK* operon (Mershad *et al*., 2016), which clusters with the two sponge thaumarchaeotes (Supporting Information Fig. S4), highlighting both environments as potential ecological reservoirs of mixotrophic thaumarchaea. Furthermore, growth of thaumarchaeotes in an industrial wastewater treatment plant that was uncoupled to ammonia-oxidation has been described (Mussmann *et al*., 2011). Interestingly, many members of the Aigarchaeota, a sister group to the Thaumarchaeota (Guy and Ettema, 2011), also encode the full *livFGHMK* operon, and a single cell aigarchaeal-like genome from cold marine sediments encodes additional extracellular proteases, di-/tripeptide transporters and aminotransferases (Lloyd *et al*., 2013). It is therefore tempting to speculate that the thaumarchaeal ancestor was a mixotroph or even strict heterotroph (as also indicated by genomic and experimental data that deep branching clade I1c and d members in the Thaumarchaeota lack genes required for ammonia oxidation; Beam *et al*., 2014; Lin *et al*., 2015; Weber *et al*., 2015), and that sponge Thaumarchaeota along with a few other members of this clade retained the capability to use amino acids due to specialized environmental conditions. Future experiments would need to demonstrate uptake of amino acids by *Ca.* N. bastadiensis to reveal whether it fuels assimilation or heterotrophic growth and/or is used for sequential intracellular generation and oxidation of ammonia (de Boer and Laanbroek, 1989; Burton and Prosser, 2001).

### Eukaryotic-like proteins (ELPs) in *Ca*. N. bastadiensis

*Ca*. N. bastadiensis encodes a number of genes containing domains postulated to have an evolutionary origin within the eukaryotes and which are thought to be important for modulating interactions between bacteria and eukaryotic hosts (Callebaut *et al*., 2000; Lurie-Weinberger *et al*., 2010; Patterson *et al*., 2014). Recent metagenomic analyses revealed an abundance of genes encoding such ELPs in the bacterial symbionts of sponges (Fan *et al*., 2012; Reynolds and Thomas, 2016; Díez-Vives *et al*., 2017), with many of these being expressed (Díez-Vives *et al*., 2017). Four types of ELPs are found in *Ca*. N. bastadiensis: Proteins with tetratricopeptide repeats (TPR), the Toll-interleukin-1 receptor (TIR) - like domain PF08937 (DUF1863; Cort *et al*., 2000), immunoglobulin-like (Ig-like) domains (DUF5011; Shigeno-Nakazawa *et al*., 2016), and hyaline repeats (HYR; Callebaut *et al*., 2000) and an extensive discussion of these ELPs is provided in the Supplementary Information. While the *Ca.* N. bastadiensis genome is enriched in proteins containing TPR, TIR, Ig-like, and HYR domains, it lacks the ankyrin repeat proteins, leucine-rich repeat proteins, protein tyrosine kinases, and armadillo-repeat proteins frequently detected in other symbiotic and pathogenic microbes (Fan *et al*., 2012; Jernigan and Bordenstein, 2015).

### Mobile and selfish genetic elements in *Ca.* N. bastadiensis

In contrast to other members of *Ca*. Nitrosopumilaceae, *Ca*. N. bastadiensis along with the other two sponge AOA, *C. symbiosum* and CcThau (but not the deep-sea glass sponge AOA) are enriched in transposases, restriction-modification (RM) systems (including a Type II restriction endonuclease, PF13156, which is exclusively found in sponge AOA - Supporting Information Fig. S5), toxin-antitoxin (T-A) systems, as well as genes putatively involved in DNA phosphorothioation (Fig. 3, Supporting Information Fig. S5). However, no differential enrichment of integrases was detected within sponge-associated thaumarchaotes. The abundance of transposases and other mobile/selfish genetic elements (MGEs/SGEs) in sponge AOA is consistent with what has been reported for other sponge-associated microbes (Fan *et al*., 2012, Horn *et al*., 2016), suggesting that evolution of AOA sponge symbionts is also affected by horizontal gene transfer in the concentrated milieu of environmental bacteria and viruses resulting from sponge feeding and pumping activity.

Among the above-mentioned genetic elements, the complete genetic repertoire for DNA phosporothioation (PT) (Wang *et al*., 2007; You *et al*., 2007) in *Ca*. N. bastadiensis and CcThau (*dndA, B, C, D, E*) is particularly noteworthy as among cultured AOA, only *Ca*. Nitrosomarinus catalina (Ahlgren *et al*., 2017) and *Ca*. Nitrosopumilus salaria also encode this DNA modification system, which acts as a primitive immune system by enabling discrimination between self and non-self DNA. In addition, *Ca*. N. bastadiensis, CcThau and *C. symbiosum* have genes with low similarity to *dndA, B* and *C,* respectively. The *dndB*-like genes contain domains (DUF262, DUF1524) previously identified as components of R-M systems (Miller et al., 2005; Machnika et al., 2015) which are known to be enriched in sponge microbiomes (Fan *et al*., 2012, Horn *et al*., 2016). *Ca*. N. bastadiensis also encodes most enzymes necessary for the production of archaeosine, a highly modified tRNA nucleoside (Phillips *et al*., 2012). Interestingly, amongst all analyzed *Thaumarchaeota*, the critical enzyme (aTGT) of this pathway is missing the RNA binding site (PUA domain) while maintaining the conserved substrate binding pocket found in *Crenarchaeota* (Phillips *et al*., 2012). Although the PUA domain is dispensable for archaeosine formation (Sabina and Söll, 2006) it is as of yet unknown whether *Ca*. N. bastadiensis can use this for DNA modification, as recently described for another system variant in a *Salmonella* species (Thiaville *et al*., 2016).

### Ammonia-oxidation is exclusively mediated by *Ca*. N. bastadiens in *I. basta* and is coupled to carbon fixation

The presence of *amoA*-encoding and expressing thaumarchaea does not prove that these microbes actually perform ammonia-oxidation in a system (Mussmann *et al.*, 2011). Consequently, nitrification rates of *I. basta* harboring the thaumarchaeal symbiont were experimentally determined to verify ammonia-oxidizing activity and to obtain insights into the mean nitrifying activity per symbiont cell. Interpretation of these data was facilitated by the fact that *I. basta* according to previous 16S rRNA gene based surveys (Webster *et al*., 2010; Luter *et al*., 2010; Freckelton *et al*., 2012; Luter *et al*., 2012) and our metagenomic and metaproteomic data contains a single AOA symbiont species and does not harbor bacterial ammonia-oxidizers or comammox organisms (Daims *et al*., 2015). Incubation experiments with freshly collected sponge clones were performed in the presence of different ammonium concentrations ranging from ambient seawater (0.29 ± 0.1 µM) via 25 μM to 100 μM. Gross and net nitrification rates as well as net fluxes of ammonium, nitrite, and nitrate were determined in 24 h laboratory incubation experiments that were repeated several times over 7 days (Supporting Information Fig. S6). Across all treatments, gross and net nitrification rates were similar in magnitude and highly correlated (gross rates = 1.13 × net rates + 0.6; R² = 0.94; *p* < 0.01), suggesting that nitrate removal does not occur at significant rates, with the exception of sponges in the 100 μM NH_4_^+^ treatment (Fig. 5A). Gross nitrification rates of sponges incubated with 25 or 100 μM NH_4_^+^ were significantly greater than in ambient seawater (*p* < 0.05; ANOVA followed by Tukey HSD Test) (Fig. 5A). The stimulation of nitrification by increased ammonium availability, which has also been observed for other sponge species (Corredor *et al*., 1988; Bayer *et al*., 2007; Bayer *et al*., 2008; Schläppy *et al*., 2010), indicates ammonium limitation of the symbiotic nitrifiers under ambient conditions. Almost no net ammonium release was observed from the *I. basta* holobiont in unamended seawater, indicating balanced ammonium production and consumption (Fig. 5B). Net nitrification rates of the *I. basta* holobiont under ambient experimental conditions were slightly higher but generally comparable to those reported for other sponge species (Supplemental Information Table S4). These data likely reflect nitrification rates under natural field conditions, as NH_4_^+^ concentrations in the unamended treatment were consistent with those reported for Orpheus Island, where *I. basta* was collected (Jompa and McCook, 2002).

**Figure 5.**
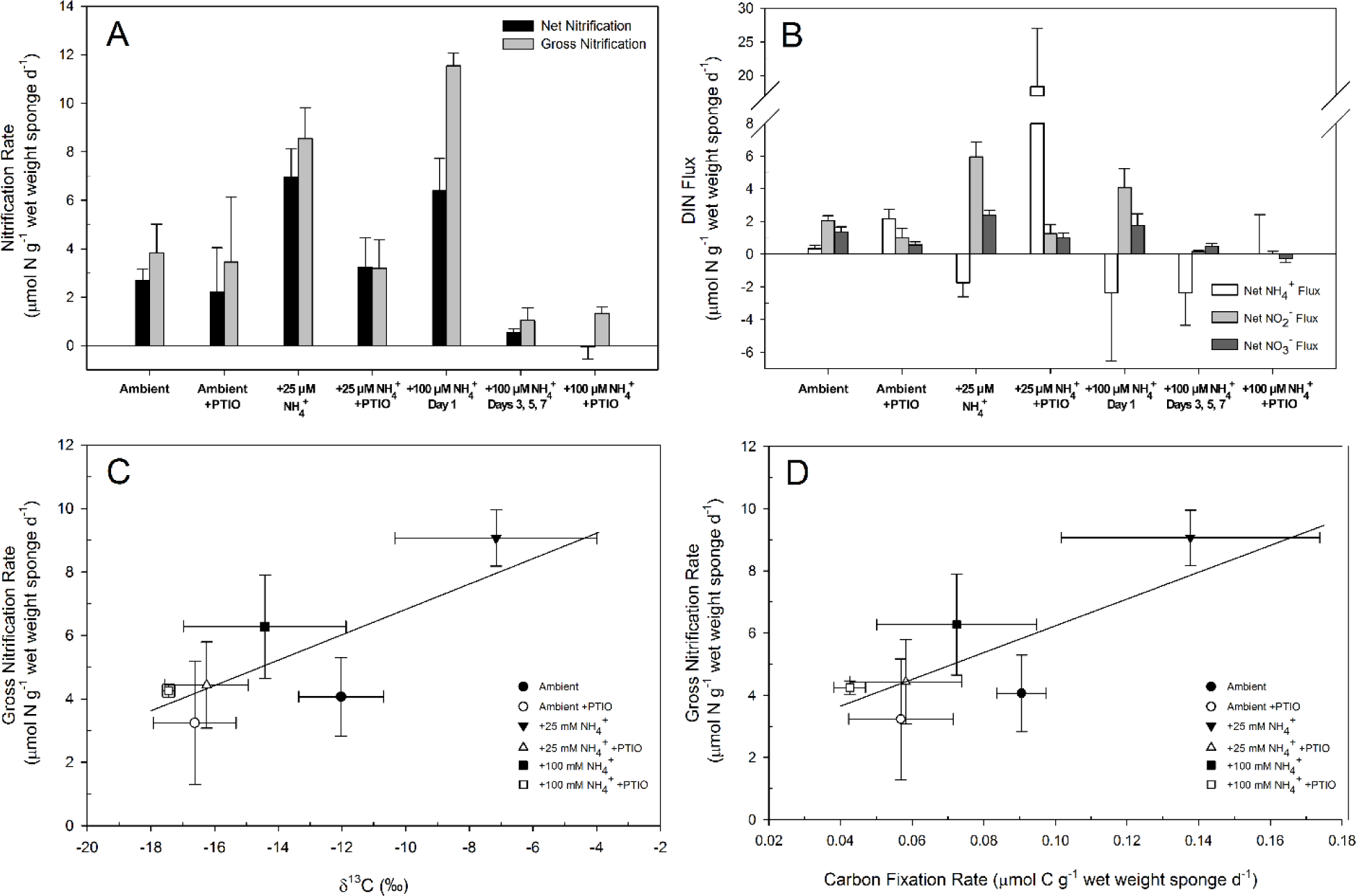
Nitrification activity of the *I. basta* holobiont during 7-day incubation experiments at ambient conditions and with added ammonium (25 or 100 µM) and with/without the AOA inhibitor PTIO. A schematic overview of the corresponding experimental setup is given in Supplementary Information Fig. S6. (A) depicts net nitrification (as calculated from the addition of net NO_2_-and NO_3_-flux) and gross nitrification rates (estimated using the ^15^NO_3_-isotope pool dilution method). (B) shows net fluxes of all DIN species for the ambient, +25 μM NH_4_^+^ and +100 μM NH_4_^+^ treatments. Rates of the different days in the ambient and +25 µM NH4^+^ incubations were averaged as they were not statistically different. (C) shows the relationship between δ^13^C values of sponge clones (labeled from added ^13^C-bicarbonate) sampled on day 7 and gross nitrification rates of the *I. basta* holobiont. To yield average nitrification activity over the course of the incubation, we used the weighted mean of the gross nitrification rates (i.e., days are weights). For comparison, the δ^13^C value reflecting the natural abundance was determined from the sponge individual used for Illumina metagenome sequencing and found to be −21.7 ‰ (not shown). The δ^13^C values derived from the *I. basta* holobiont nitrification experiments displayed a positive and significant correlation with gross nitrification rates (R = 0.656, *p* < 0.005). (D) shows the relationship between gross nitrification rates of the *I. basta* holobiont and the carbon fixation rate of sponge clones sampled on day 7. The carbon fixation rates derived from the *I. basta* holobiont nitrification experiments displayed a positive and significant correlation with gross nitrification rates (R = 0.665, *p* < 0.005). Error bars in panels C and D reflect the standard error of the sample mean, where n = 3 for the carbon fixation rates, for the δ^13^C values, and for all weighted gross nitrification rates in all treatment conditions. PTIO additions started two days after the experiment was commenced, hence fixation of ^13^C-labeled bicarbonate during the first two days was not influenced by inhibition.

Net nitrification rates in sponges subjected to 100 μM NH_4_^+^ for 24 h were significantly lower than gross nitrification rates (*p* < 0.05; two-tailed t-test, Fig. 5A), indicating increased nitrate consumption. Gross and net nitrification rates also became significantly depressed after prolonged exposure (> 3 days) to 100 μM NH_4_^+^, being significantly lower than in the 24 h incubations at 100 μM NH_4_^+^ as well as in the ambient seawater controls (*p* < 0.01, Mann-Whitney U-test) (Fig. 5A). From AOA pure culture studies, there are no indications that 100 μM NH_4_^+^ can be inhibitory for members of this clade, with concentrations of 2 to 20 mM NH_4_^+^ needed for inhibition (Hatzenpichler *et al*., 2008; Martens-Habbena *et al*., 2009; Tourna *et al*., 2011; Li *et al*., 2016; Sauder *et al*., 2017; Sauder *et al*., 2018). While we cannot exclude that *Ca*. N. bastadiensis is particularly sensitive to ammonium, it is also possible that the inhibition was caused by indirect effects. For example, if the sponge host was adversely affected at 100 μM NH_4_^+^, decay of sponge cells could cause stimulation of heterotrophic microbes and thus oxygen limitation. Typical acute toxicities in freshwater fish and invertebrates (96-hour LC_50_) are reached at unionized ammonia (NH_3_) concentrations between 4.7 – 38 μM (with salinity typically having a slight ameliorating effect and tests on marine invertebrates very rare; Boardman *et al*., 2004) and susceptibility to ammonia toxicity can be increased by low O_2_ concentrations (Camargo and Alonso, 2006). At ambient incubation conditions about 4.5% of total ammonia was present as NH_3_, hence *I. basta* may have experienced physiological stress and intermittent hypoxic conditions in the 100 μM NH_4_^+^ treatments.

PTIO, a scavenger of free radical nitric oxide (NO) (Amano and Noda, 1995; Ellis *et al*., 2001), has been described as a specific inhibitor of AOA (Yan *et al*., 2012; Martens-Habbena *et al*., 2015) and was therefore used to demonstrate that *Ca*. N. bastadiensis is responsible for ammonia-oxidation in *I. basta*. As expected, the addition of PTIO to sponge incubations in the ambient and 25 μM NH_4_^+^ amended treatments, resulted in significantly depressed gross and net nitrification rates and increased ammonium accumulation when compared to the non-PTIO treated sponge incubations (*p* < 0.05; except for gross nitrification rates under ambient conditions, p = 0.885; Mann-Whitney U-test; Fig. 5A and Fig 5B). In the case of the 100 μM NH_4_^+^ amendment, gross and net nitrification was already strongly reduced in the absence of PTIO after 3 days of incubation, so further inhibition by PTIO could not be demonstrated under these conditions (Fig. 5A). Residual ammonia oxidizing activity in some of the experiments in the presence of 75 μM PTIO was unexpected as pure AOA cultures are known to be fully or almost completely inhibited by this concentration of PTIO (Shen *et al*., 2013; Martens-Habbena *et al*., 2015). NO production by *I. basta* (many sponges express NO synthases, Riesgo *et al*., 2014) may have contributed to PTIO inactivation thereby lowering its inhibitory effect on the AOA. To our knowledge, PTIO has not previously been used in sponge microbiome research (but is commonly applied to eukaryotic tissues including sponges; Müller *et al*, 2006; Ueda *et al*., 2016) and further optimization of the concentration is recommended for future applications in these animals.

In 29 of the 33 sponge incubations (incubations with added PTIO or inhibited by addition of 100 μM NH_4_^+^ were excluded) (Fig. 5B), accumulation rates of NO_2_-exceeded those of NO_3_-by an average factor of 2.8 and this difference was found to be significant in the ambient and 25 μM NH_4_^+^ treatments (both *p* < 0.01, Mann-Whitney U-test). In contrast, no significant difference between NO_2_-and NO_3_-production was observed in the PTIO-amended incubations nor in those experiments where prolonged exposure to 100 μM NH_4_^+^ negatively affected net nitrification rates (all: *p* > 0.05, Mann-Whitney U-test).

All nitrification experiments were performed in intermittently closed aquaria (Supporting Information Fig. S6) to prevent loss of labeled CO_2_ but to provide sufficient oxygen exchange to maintain sponge health. Additional control experiments were performed at ambient and 100 μM NH_4_^+^, to compare DIN fluxes and inferred net nitrification rates using this setup, to incubations performed in constantly open containers. Interestingly, slightly but significantly higher NO_2_-fluxes and net nitrification rates were observed with ambient seawater in the intermittently closed aquaria (both *p* < 0.01, two-tailed t-test; Supporting Information Fig. S7), indicating that ammonia-oxidizers in the closed system either benefited from the minimized loss of the added bicarbonate (via CO_2_ off-gassing) or from potentially increased ammonia production from stressed sponge clones.

To determine the contribution of the seawater microbial community to net nitrification, rates were also determined from aquaria seawater without sponges, revealing minimal nitrification with net rates of only 0.06 ± 0.3 μM N d^-1^ (n = 12).

To calculate cell specific ammonia oxidation rates for *Ca.* N. bastadiensis, its 16S rRNA genes were quantified at the end of the 7 day sponge incubation under ambient and +25 μM NH_4_^+^ treatments. Sponges subjected to ambient and +25 μM NH_4_^+^ treatments contained thaumarchaeal symbiont gene copy numbers of 8.58 ± 4.9 × 10^10^ and 1.64 ± 0.12 × 10^10^ per g wet weight (SE, n = 3, for both), respectively. By dividing the net nitrification rate on day 7 by the 16S rRNA gene copy number of *Ca*. N. bastadiensis on this day, average cell specific rates were estimated to be 0.11 ± 0.08 and 0.66 ± 0.09 fmol NH_4_^+^ oxidized per cell per day (SE, n = 3, for both) for the ambient and +25 μM NH_4_^+^ incubations, respectively. Inferred cell specific ammonia oxidation rates for *Ca*. N. bastadiensis were ~20-120 times lower than what has been reported for *N. maritimus* (Martens-Habbena *et al*., 2009) and ~6-65 times lower than coastal marine seawater (Wuchter *et al*., 2006), while the highest ammonia oxidation rate of *Ca*. N. bastadiensis was comparable to the lowest rates detected in the sponge *Phakellia ventilabrum* (Radax *et al*., 2012a).

Furthermore, as ^13^C-bicarbonate was added during all experiments (Supporting Information Fig. S6), we were able to measure the δ^13^C values of sponge tissues and calculate inorganic carbon fixation rates to compare with the gross nitrification rates. PTIO was found to reduce the δ^13^C value in both the ambient and +25 μM NH_4_^+^ treatments (Fig. 5C; both: *p* < 0.05, one-tailed t-test), while no significant differences were observed with the +100 μM NH_4_^+^ treatments. In addition, gross nitrification rates and carbon fixation were significantly positively correlated (Fig. 5D). These data suggest that *Ca*. N. bastadiensis contributes significantly to carbon fixation by the sponge holobiont. This finding is consistent with (i) the detection of key genes for CO_2_ fixation in the *I. basta* thaumarchaeote metagenome bin, (ii) the detection of some of the respective proteins in the metaproteome, (iii) and the absence of autotrophic CO_2_ fixation pathways in the metagenomic bins of the alpha-and gamma-symbionts (data not shown). In addition, the slope of the positive linear relationship between the gross nitrification rate and the carbon fixation rate (Fig. 5D; 43.0), reflects the gross nitrification: carbon fixation ratio (N:C ratio), which is most likely over-estimated since the experimental setup of the intermittently closed aquaria still allowed for the loss of added ^13^C-bicarbonate. In addition, CO_2_ production derived from respiratory activities within the *I. basta* holobiont were not accounted for, which may also lead to an underestimation of carbon fixation rates. Despite these possible biases the N:C ratio determined for *I. basta* is within the range typically found for aquatic environmental samples (2-60)(Andersson *et al*, 2006). The relatively high N:C ratio along with the low cell-specific ammonia-oxidation rates within the *I. basta* holobiont are consistent with the proteogenomic-derived hypothesis that *Ca.* N. bastadiensis is not sustaining its population using ammonia as its only substrate, but exploits peptides and/or amino acids as additional sources of energy, nitrogen, and carbon by growing as a mixotroph.

Surprisingly, nitrite-oxidizing bacteria (NOB) could not be detected in *I. basta* by amplicon sequencing, metagenomic sequencing, or with FISH. Consistent with the absence of NOB, NO_2_^-^ accumulated to high concentrations ranging from 8 – 21 μM in ambient treatments containing *I. basta* (Fig. 5B, Supporting Information Fig. S7), making this sponge one of the few natural systems in which greater NO_2_-than NO_3_-concentrations occur (Brezonik and Lee, 1968; Lam *et al*., 2011; Schaefer and Hollibaugh, 2017 and refs. therein). It will be interesting to explore in future studies whether nitrite production contributes (via nitrite toxicity; Camargo and Alonso, 2006 and references therein) to protection from predators or to the unusually low microbial diversity in *I. basta*.

Given the absence of NOB in *I. basta*, the observed production of nitrate is difficult to explain. One possibility is that some NO released by either the host or a host-associated microorganism is detoxified to nitrate by another member of the sponge holobiont. However, we did not detect genes with homology to those encoding the NO detoxifying enzyme, flavohemoglobin-NO-dioxygenase (Hmp), which converts NO together with O_2_ to NO_3_-(Gardner, 2012 and references therein), in our microbial metagenomic datasets. Still, NO dioxygenases (NODs), could be encoded in the sponge genome since neuroglobin-like sequences are encoded in other sponges (Lechauve *et al*., 2013) and members of this enzyme family have been shown to have NOD activity *in vitro* (Brunori *et al*., 2005). Furthermore, a NADH-cytochrome *b5* reductase, which belongs to the same family as Hmp, was found to be highly expressed at all life stages of the sponge *Amphimedon queenslandica* (Conaco *et al*., 2012). Alternatively, partial nitrite oxidation may be catalyzed by free-living NOB in the seawater, although nitrification rates in the control were negligible.

### Conclusions and outlook

In this study we combined proteogenomic and experimental analyses to show that *Ca.* N. bastadiensis, the first characterized but yet uncultured representative of a new genus within the thaumarchaeotes, is responsible for ammonia oxidation in the widespread marine sponge *I. basta*. While *Ca.* N. bastadiensis is equipped with the typical genetic repertoire of free-living AOA, it also exhibits a number of putative adaptations to a host-associated lifestyle. Several of these adaptive features were unique to *Ca.* N. bastadiensis, whereas others were shared exclusively with the previously described sponge symbionts *Ca.* C. symbiosum (Hallam *et al*., 2006) and the AOA symbiont of the sponge *C. concentrica* (Moitinho-Silva *et al*., 2017a). Many of these putative adaptations to a sponge-associated lifestyle were not encoded in the recently sequenced genome of an AOA from a glass sponge (Tian *et al*., 2016), indicating that this archaeon is not an obligate symbiont or that thriving in a glass sponge requires very different traits. Our results confirm an emerging view that marine sponge microbiomes tend to converge on shared functional traits, a process shaped by the environmental niche provided by the sponge host and governed by specific ecological factors such as high dissolved nutrient loads and frequent contact with resident and transient microorganisms (Liu *et al*., 2012). Whereas other studies have focused on this phenomenon by comparing functional convergence across entire sponge microbiomes (Fan *et al*., 2012; Liu *et al*., 2012; Horn *et al*., 2016), we demonstrate that in symbiotic marine sponge *Thaumarchaeota*, a similar evolutionary convergence is achieved that stands in contrast to their strictly chemoautotrophic non-symbiotic free-living relatives and that this process is likely not solely achieved by gene acquisition but rather by selective gene retention and gene family expansion.

With *I. basta* emerging as a model species for sponge symbiosis research, additional work should be undertaken to address the conspicuous absence of nitrite-oxidizing bacteria and ascertain why, in contrast to most tropical sponge species, *I. basta* hosts such a low diversity of microbial symbionts. Furthermore, hypotheses about the interaction of *Ca.* N. bastadiensis with other members of the sponge holobiont should be experimentally confirmed. For example, assimilation of amino acids and peptides should be tested using stable isotope probing and the function of unique serine proteases, serpins and ELPs should be analyzed via heterologous gene expression to reveal mechanistic insights into the interaction of this archaeon with its host. Finally, the mechanism for symbiont acquisition should be assessed via screening for *Ca*. N. bastadiensis in gametes and larvae.

## Experimental Procedures

### Sponge collection

Large adult specimens (n=4) of the sponge *Ianthella basta* were collected from Orpheus Island (18°36.878’S, 146°29.990’E), Queensland, Australia, cut into 10 cm × 10 cm explants and transferred to racks on the reef. After a 12 week healing period in the field, sponge clones were collected in two separate sampling trips by scuba diving between September and October 2011 and transported to the indoor temperature-controlled aquarium at the Australian Institute of Marine Science (AIMS), Townsville, where they were acclimated at ambient temperature (25 ºC) for 48 h and then randomly assigned to experimental treatments. The two adult *I. basta* specimens used for metaproteogenomic analyses were collected from Orpheus Island in October 2010 and 2011 at Orpheus Island. Upon sample collection, specimens were cut into small strips, immediately snap-frozen in liquid N_2_ and subsequently stored at −80 ºC until DNA extraction.

### Sponge incubations and nitrification rate measurements

To infer nitrification activity of the *I. basta* holobiont, a 7-day incubation experiment was performed using triplicate sponge clones under different ammonium concentrations: ambient (0.29 ± 0.1 µM), 25 and 100 µM NH_4_^+^. A schematic representation of the experimental design is given in Supporting Information Fig. S6. Specifically, sponge clones (1.5 – 13.0 g post-experimental wet weight; mean = 5.1; standard deviation = 2.2) were incubated at 25 ºC in the dark in 1.5 L acid-washed glass containers completely filled with 5 μm filtered, seawater or with seawater amended with 25 μM or 100 μM NH_4_Cl. Additionally, abiotic control experiments were performed with ambient seawater without sponge clones that were incubated for 24 h under identical conditions. Sponge clones were transferred to a new container with fresh seawater every 24 h during the 7-day incubation, in order to reduce the effects of O_2_ depletion and NH_4_^+^ accumulation. To assess carbon fixation in concert with nitrification, NaH^13^CO_3_-(100 μM; 99% ^13^C) was added every 12 h to all experimental treatments. To prevent loss of ^13^C-labeled CO_2_, containers were closed for six hours after the NaH^13^CO_3_-addition and subsequently opened until the next NaH^13^CO_3_-addition to avoid oxygen depletion. The effect of such intermittently closed containers on nitrification activity was assessed in a control experiment by comparing ambient seawater and 100 μM NH_4_Cl amended seawater in constantly open containers (Supporting Information Fig. S7). Nitrification activity was determined as gross nitrification rates using ^15^N isotope pool dilution technique (Inselsbacher *et al*., 2007), and net fluxes of individual dissolved inorganic N species (DIN), namely NH_4_^+^, NO_2_-, NO_3_-. Gross nitrification rates were also measured on day 1, 3 and 5, where at the beginning of each day K^15^NO_3_ was added to a final ^15^NO_3_-concentration of 1 – 10% of the nitrate pool (Murphy *et al*., 2003) in the filtered ambient seawater. Gross nitrification rates were calculated from seawater samples collected after a short equilibration following ^15^N-label addition (0 h) and after 20 h. Net fluxes of NH_4_^+^, NO_2_-, and NO_3_-were measured on day 1, 3, 5 and 7 and were calculated as change in concentration over time (i.e., between 6 and 18 h after ^15^N-label addition to avoid biased net NO_3_-fluxes through a stimulation of consumptive processes from the added ^15^NO_3_-). Net nitrification refers to the change in concentration of NO_2_-+NO_3_-. Furthermore, we used the AOA-specific inhibitor PTIO (2-phenyl-4,4,5,5-tetramethylimidazoline-1-oxyl 3-oxide; Tokyo Chemical Industry) (Martens-Habbena *et al*., 2015) in additional parallel incubations, which was added at a concentration of 75 µM at the beginning of day 3 and 7.

Seawater samples (10 mL) were taken from each aquarium and filtered using 0.45 µm Sartorius Minisart cellulose acetate filters (Göttingen, Germany). Duplicate samples for dissolved inorganic nitrogen (NH_4_^+^, NO_2_-, NO_3_-) were measured on a Seal AA3 segmented flow analyzer and referenced against OSIL standards and in-house reference samples. For ^15^N-analysis of NO_2_-+NO_3_-, sample water was filtered through pre-combusted GF/Fs (Whatman International; treated for 4 h at 450°C), and subsequently through 0.2 µm filters (Sartorius). All samples were immediately frozen at −20 ºC for later analysis. Prior to shipment to the University of Vienna, samples were thawed at room temperature and the microbial inhibitor pheynylmercuric acetate was added (to a final concentration of 10 µM). Upon arrival in Vienna the samples were promptly stored at −80ºC. Nitrite and nitrate were isolated together from seawater by sequential microdiffusion (Sørensen and Jensen, 1991). To remove ammonium from sample water, 100 mg MgO, and an acid trap (acidified cellulose filter disc enclosed in a semi-permeable Teflon membrane) was added to 9 mL of sample and 1.5 mL of 3 M KCl. After 5 days shaking at 35ºC, the acid traps were removed, and 50 mg of Devarda’s alloy was added along with a new acid trap, and shaken at 35°C for 7 days. Devarda’s alloy is a reducing catalyst converting both NO_2_-and NO_3_-to NH_4_^+^ and the subsequently formed NH_3_ was collected in the acid trap. Acid traps were dried over concentrated sulfuric acid and analyzed for ^15^N by an elemental analyser (EA 1110, CE Instruments, Milan, Italy) coupled to an isotope ratio mass spectrometer (IRMS; Finnigan MAT Delta^Plus^ IRMS with a Finnigan MAT ConFlo III interface). Gross nitrification rates were calculated based on Wanek *et al*. (2010). Gross and net rates are expressed as µmol nitrogen species per g wet weight sponge per day (µmol N g^-1^ d^-1^).

For the determination of ^13^C enrichment in whole sponge tissue at the end of the incubation, sponge tissue was freeze-dried, ground to a fine powder, and stored at dry conditions prior to analysis. The δ ^13^C values of sponge tissue were determined using an EA-IRMS system as described above. Carbon fixation rates were calculated based on the δ ^13^C values treatments amended with 100 µM NaH^13^CO_3_-. For the calculations, we used a background (natural abundance) bicarbonate concentration of 1975 µM.

### DNA extraction from whole sponge tissue and qPCR for symbiont quantification

Between 80-150 mg of *I. basta* tissue was thawed, rinsed successively (3x) in 1X calcium-and magnesium-free artificial seawater (CMF-ASW) and immediately ground into a paste with a mortar and pestle in liquid N_2_. After resuspension in TE buffer (10 mM Tris–HCL pH 8.0, 1 mM EDTA), DNA was extracted from the suspension using an adapted SDS-based isolation method (Zhou *et al*., 1996) and using 1% polyvinylpyrrolidone. DNA was extracted from the two individuals used for metaproteogenomics (see below) as well as three healthy individuals from a previous study (Luter *et al*., 2010). Additionally, DNA was extracted from a subset of sponge clones that were subjected to nitrification incubations. Quantitative PCR (qPCR) was used to estimate the number of specific thaumarchaeal, as well as α-and γ-proteobacterial symbionts by quantifying the 16S rRNA gene using specific primers designed for each symbiont phylotype. The following primer sets were thus used for the dominant *I. basta* thaumarchaeal, and α-, and γ-proteobacterial symbionts respectively; IBthaum16S_523F, 5′-CCG TAG CCT GCC CTG TTA G-3′, IBthaum16S_727R, 5′-GCT TTC ATC CCT CAC CGT-3′; IBalpha16S_1010F, 5′-CGG AGA CGC TTC CTT CG −3′, IBalpha16S_1206R, 5′-GCC CAG CCC ATA AAT GC-3′; IBgamma16S_466F, 5′-TAC CCY TGY GTT TTG ACG-3′, IBgamma16S_655R, 5′-CCR CTT CTC TCT RCC ATA C-3′. An iCycler real-time PCR system (Bio-Rad, Hercules, CA) was used to measure all samples, in duplicate wells per reaction and reactions were performed in a 25 μL volume with 1 μL of DNA template. All symbiont 16S rRNA gene qPCR assays used SYBR®Green reaction mixtures containing 12.5 μL iQ SYBR^®^Green Supermix (Bio-Rad), and optimized concentrations of 400 nM primer as well as 0.25 mg ml^-1^ BSA. Cycling conditions were 95ºC for 5 min followed by 40 cycles of 95ºC for 40 s, 58ºC for 30 s, and 72ºC for 40 s. Fluorescence intensities were measured after each cycle, and a final elongation at 72ºC was followed by a melting curve analysis from 55-95ºC in 10 s increments of 0.5ºC.

Standard curves were generated for each primer set using serial dilutions of a standard containing a known number of the target sequences. Standards used the M13 primer set to amplify 16S rRNA gene clones derived from the sponge symbionts. PCR products were visualized on an agarose gel, purified separately using the QIAquick PCR Purification Kit (Qiagen), followed by fluorometric quantification of DNA concentrations using PicoGreen (Molecular Probes, Eugene, OR) and a NanoDrop ND-3300 Fluorospectrometer (NanoDrop). Gene abundance was calculated based on DNA concentration and product size. Dilution series ranging from 10^6^ to 10^0^ copies μL^-1^ were used to generate standard curves. Final 16S rRNA gene abundances for each microbial symbiont were then normalized by the wet weight of the sponge tissue used for DNA extraction.

### Cryosectioning and FISH for symbiont quantification

The *I. basta* individual collected for metagenomic sequencing in October 2010 was also assessed using FISH. Briefly, after sample collection, the *I. basta* specimen was cut into tissue strips of approximately 2 mm^3^, fixed in 4% PFA for 1 h at room temperature and stored in ethyl alcohol (EtOH)-phosphate-buffered saline (PBS) at −20ºC. For FISH, PFA-fixed samples of *I. basta* were embedded in Neg-50 (Richard-Allan Scientific), and cut to 5-μm sections (Leica CM3050 S). A double-labeled Arch915 probe in Cy3 (Thermo Fisher Scientific, Waltham, MA, USA) was used for the microscopic visualization and quantification of the thaumarchaeal symbiont of *I. basta*. To calculate the relative abundance of the thaumarchaeal symbiont, equimolar amounts of the double-labeled probes EUB338-I, EUB338-II, and EUB338-III (Fluos and Cy5) (Stoecker *et al*., 2009) were used for quantification of most bacteria. Hybridizations were prepared with an equimolar mixture of both probes and using 25% formamide in the hybridization buffer, with the stringency of the washing buffer adjusted accordingly. As a negative control, the nonEUB338-I (reverse complementary probe to EUB338-I) was applied on one section per well per slide hybridized (Wallner *et al*., 1993). All hybridized samples were analyzed with a confocal laser scanning microscope (CLSM) (LSM 510 Meta; Zeiss, Oberkochen, Germany). Archaeal and bacterial cells were counted by eye on 10 randomly selected images derived from multiple tissue sections obtained from a single *I. basta* individual, and the proportion of archaeal cells to total prokaryotic cells was calculated.

### Microbial cell enrichment for metaproteogenomics

To separate symbiont cells from host tissue prior to DNA extraction, approximately 15.7 g wet weight of *I. basta* was rinsed successively (3x) in 1X CMF-ASW. Sponge tissue was cut into small pieces (<1 cm^3^), ground on ice in 1X CMF-ASW with a mortar and pestle, transferred into a glass douncer on ice, and sponge tissue was dissociated through shear force and vortexing. The supernatant was transferred into multiple Eppendorf microcentrifuge tubes, centrifuged at 39 × g for 15 min at 4ºC to pellet larger sponge particles, filtered through a 5 μm Sartorius filter and centrifuged at 10,844 × g for 15 min at 4 ºC to pellet microbial cells. Microscopic examination of the pellet revealed an enrichment of microbial cells and an absence of sponge nuclei. For the sponge individual sampled in October 2010, DNA was extracted from this pellet using the procedure referenced above. For the individual sampled in October 2011, the pellets were resuspended in 1X CMF-ASW and the suspension was layered in 1.8 ml amounts onto 6.5 ml cushions of 30% Gastrografin dissolved in 1X CMF-ASW + 0.2 M EDTA, and centrifuged at 40,008 × g for 1 hour at 4ºC in an Beckman L-100 XP ultracentrifuge with the SW 41 Ti swing rotor (Beckman Coulter). Following centrifugation, a cell-rich layer above the 30% Gastrografin cushion (‘Fraction A’) and the cell-rich pellet (‘Fraction B’) were carefully removed, resuspended in 10 mM Tris, and re-pelleted at 24,400 × g for 15 min at 4ºC in a new solution of 10 mM Tris.

The same *I. basta* individual sampled in October 2011 was used for proteomic analyses. However, in order to minimize protein degradation, biomass preparation methods were shortened and modified. The first cell fraction used for proteomic analysis involved rinsing in 1X CMF-ASW and tissue homogenization using a mortar/pestle and glass douncer as described above. However, the tissue homogenization steps occurred in 1X TE buffer with Roche Complete Protease Inhibitor (Roche). Supernatant from the glass douncer was immediately collected into 2 mL Eppendorf tubes and frozen at −80ºC before shipment on dry ice to Greifswald, Germany, for protein extraction and downstream analyses. A second cell fraction comprised a 5 μm filtrate from the supernatant in the glass douncer. Finally, a crude sponge homogenate sample was obtained by direct grinding of sponge tissue (without rinsing in 1X CMF-ASW) in liquid N_2_ followed by freezing at −80ºC in 1X TE with Roche Complete. These three fractions could thus be characterized as a sponge homogenate without sponge skeleton, a sponge homogenate without sponge nuclei, and a direct sponge homogenate, respectively.

### DNA extraction, library preparation and sequencing

DNA was extracted from the individual sampled in October 2011 using the FastDNA spin kit for soil (MP Biomedicals, Solon, OH, USA) from the cell-rich layers above the 30% Gastrografin cushion and in the ultracentrifuged pellet according to the manufacturer’s instructions. Sequencing libraries were prepared using the Nextera kit (Illumina Inc.) according to the manufacturer’s instructions and concentrations measured using the QuantIT kit (Molecular Probes, Life Technologies, Naerum, Denmark). The libraries were paired-end (2×150 bp) sequenced on an Illumina HiSeq2000 using the TruSeq PE Cluster Kit v3-cBot-HS and TruSeq SBS kit v.3-HS sequencing kit and on an Illumina MiSeq using v3 2×300 bp kits.

For the individual sampled in October 2010, cell pellets were pooled from the 5 μm filtration step and DNA was extracted using the modified protocol of Zhou *et al*. (1996) described above. Metagenomic sequences were then generated at the Ramaciotti Sequencing Centre (Sydney, Australia) using the GS FLX instrument using Titanium chemistry (Roche) on a 454 half-sequencing-plate (454 Life Sciences, Branford, CT, USA).

### Metagenome assembly and genome binning

Metagenome reads in fastq format, obtained from the Illumina sequencing runs, were end-trimmed at a minimum phred score of 15, a minimum length of 50 bp, allowing no ambiguous nucleotides and Illumina sequencing adaptors removed. Trimmed reads from each dataset were assembled using Spades version 3.11.0 (Bankevich *et al*., 2012), using default parameters and genomes were binned using Metabat v 2.12.0 (Kang *et al*., 2015). MAGs from multiple Illumina datasets were dereplicated using dRep (Olm *et al*., 2017) and provisionally classified using CheckM (Parks *et al*., 2015). A single high-quality archaeal MAG was recovered after dereplication and uploaded to MaGe (Vallenet *et al*., 2009) for annotation. This MAG has been submitted to the European Nucleotide Archive with the accession number PRJEB29556.

For the 454-pyrosequencing run, artificially amplified reads were dereplicated using CD-HIT (Li and Godzik, 2006), assembled with MIRA (Chevreux *et al*., 1999) and binned with a genome-specific Phymm model (Brady and Salzberg, 2009), which was trained from the contigs that contained phylogenetic marker genes. The MAG derived from this pyrosequencing run was similarly GC-rich (64.2%), but considerably larger (6.58 Mb), slightly less complete (98.06%), and considerably more fragmented (2508 scaffolds). This MAG is available on the MaGe platform as “*Ianthella basta symbiont* thaum” at http://www.genoscope.cns.fr/agc/microscope/home/index.php.

### Comparative genomics

The annotated archaeal MAG was downloaded from MaGe and compared to published thaumarchaeotal genomes (Supporting Information Table S5) using genomic average nucleotide identity (gANI), average amino acid identity (AAI), and through construction of orthologous gene families. For all analyses, annotated genes were supplemented with additional gene calls predicted by Prodigal (Hyatt *et al*., 2010). gANI was calculated with MiSI (Varghese *et al*., 2015). AAI (Konstantinidis and Tiedje, 2005) was calculated using bidirectional best blastp hits (Camacho *et al*., 2009) that aligned over at least 70% of gene length with average identity values weighted according to gene length. Orthologous gene families were constructed using Orthofinder (Emms and Kelly, 2015). For functional annotation of eukaryotic-like proteins (ELPs) and mobile/selfish genetic elements, predicted genes from all sequenced thaumarchaeal genomes were searched against the Protein Family A (Pfam-A) database (v31.0) (Finn *et al*., 2014) using Hmmer 3 (hmmer.org.) and the gathering threshold option (-cutga). Results were screened for the presence of domains associated with ELPs and mobile/selfish genetic elements.

### Phylogenetic analyses

Bayesian trees were constructed using Phylobayes v 4.1c (Lartillot and Philippe, 2004) using the best model identified for each dataset by ModelFinder (Kalyaanmoorthy *et al*., 2017). Phylogenomic reconstruction was based on a concatenated amino-acid alignment of 34 marker genes constructed with CheckM (Parks *et al*., 2015) with ten independent runs of 11000 generations under the LG4 model. 6000 generations of each independent run were discarded as burn-in and the remaining trees from each run were pooled for calculation of a consensus tree and for determining posterior branch support. For the 16S rRNA gene phylogenies, top representative hits in a blastn query against the Genbank nr database of the full-length sequence from *Ca*. N. bastadiensis and *Ca*. C. symbiosum, along with sequences from sequenced thaumarchaeal genomes were aligned with SINA (Pruesse *et al*., 2012) and analyzed further as described above. For the *amoA* gene tree, the *Ca*. N. bastadiensis and *Ca.* C. symbiosum *amoA* sequences were placed into a reference tree, representing all OTU representatives of the curated database provided in Alves *et al*. (2018), using the Evolutionary Placement Algorithm (EPA; Berger *et al*., 2011) implemented in RAxML-HPC 8.2.11 (Stamatakis, 2014). Representative sequences clustering with the *Ca*. N. bastadiensis *amoA* gene sequence along with *amoA* sequences from the aforementioned genome-sequenced thaumarchaeota were then aligned with MUSCLE (Edgar, 2004) and analyzed. For both 16S rRNA and *amoA* gene phylogenies, ten independent runs of 30000 generations under the GTR model were used. 7500 generations of each independent run were discarded as burn-ins and the remaining trees from each run were pooled for calculation of a consensus tree and for determining posterior branch support.

### Protein identification and proteome analyses

1D PAGE followed by liquid chromatography-based mass spectrometry (1D-PAGE-LC-MS/MS) were used for protein and peptide separation and identification as described previously (Washburn *et al.*, 2001; Otto *et al.*, 2010), with slight modifications. MS spectra and MS/MS spectra were acquired from eluting peptides ionized with electrospray ionization (ESI) and analyzed in a LTQ Orbitrap Velos hybrid mass spectrometer (Thermo Fisher Scientific, Waltham, MA, USA), as described previously (Verberkmoes *et al.*, 2009; Otto *et al.*, 2010), with minor modifications. Samples of the sponge homogenate without sponge skeleton and crude sponge homogenate processed in liquid N_2_ were analyzed in technical duplicates, whereas the sponge homogenate without sponge nuclei was analyzed only once. All MS/MS spectra were searched against predicted protein sequence databases composed of the *I. basta* symbiont-enriched metagenome bins and common laboratory contaminants using the Sorcerer SEQUEST (v.27, rev. 11) algorithm. The CD-HIT software (Li and Godzik, 2006) was used to remove redundancies from the database due to the potential for strain-level redundancies. Protein identifications were filtered with Scaffold version 3.5.1 applying the “sequest” filter criteria described previously (Heinz *et al*. 2012). For protein identification only peptides identified with high mass accuracy (maximum ± 10 ppm difference between calculated and observed mass) were considered and at least two exclusively unique peptides were required to identify a protein. False-discovery rates (FDRs) were estimated with searches against a target-decoy database as described previously (Peng *et al*., 2003; Elias and Gygi, 2007). Peptide FDRs were between 2.5% and 3.1%, and protein FDRs were below 0.4% throughout all samples. For relative quantitation of proteins, normalized spectral abundance factor were calculated for each sample according to the method of Florens *et al*. (2006) and averaged for all replicates and samples. The proteomics dataset for this study have been submitted to the PRIDE archive.

## Acknowledgements

This work was supported by the Marie Curie Initial Training Network - SYMBIOMICS. In addition, F.U.M., C.W.H, M.M., and M.W. were supported by the European Research Council Advanced Grant project NITRICARE 294343. M.A. and P.H.N. were supported by research grants (15510 and 16578) from VILLUM FONDEN. Many thanks to Christian Hentschker for Orbitrap Velos measurements and database searches.

