## Supplementary Materials for "Characterization of a thaumarchaeal symbiont that drives incomplete nitrification in the tropical sponge *Ianthella basta*"

### Table of contents:

1. Supplementary Results and Discussion

2. Supplementary Experimental Procedures

3. Figures S1 – S7

4. Table S1-S5

5. References

### Supplementary Results and Discussion

**High GC content of *Ca.* Nitrosospongia bastadiensis.** Both thaumarchaeal symbiont MAGs from *I. basta* possess the highest GC content (64.8%) of any genome-sequenced thaumarchaeote (Fig. 2C). While the GC content in the genome of *Ca.* Cenarchaeum symbiosum is similarly high (57.4%), the thaumarchaeal MAGs from the sponges *Cymbastela concentrica* (Moitinho-Silva *et al*., 2017a) and a glass sponge (Tian *et al*., 2016) had a much lower GC content (Fig. 2C). The high GC content found in the *I. basta* thaumarchaeal symbiont is consistent with the high GC content evident in Mediterranean sponge metagenomes (GC content 58-63%), particularly when compared to seawater metagenomes (GC content 41%) collected at the same location (Horn *et al*., 2016). Similarly, the GC content of six sponge microbiome metagenomes (Fan *et al*., 2012) that we queried had an average GC content of 57.8% ± 6.7(SD), with three great barrier reef sponge species having an average microbiome metagenomic GC content of 63.3% ± 2.2 (SD). In contrast, the average GC composition of selected marine and coral microbiomes are ~48% and ~45%, respectively (Reichenberger *et al*., 2015). While several environmental factors are thought to affect the genomic GC content of bacteria and archaea (Foerstner *et al*., 2005; Wang *et al*., 2006), increased rates of homologous recombination via GC-biased gene conversion has recently been proposed as a crucial factor universally influencing the nucleotide content of microbial genes and genomes (Lassalle *et al*., 2015). The possibility that sponge microbiomes are hot spots for input of novel genetic material via lateral gene transfer events (Fan *et al*., 2012; Horn *et al*., 2016), and also display increased homologous recombination rates should be assessed in future work.

**Environmental distribution of *Ca.* Nitrosospongia bastadiensis**. The *Ca*. N. bastadiensis 16S rRNA gene was queried against the Sponge Microbiome Project (SMP) database containing amplicon data sets from 268 sponge species including *I. basta* (Moitinho-Silva *et al*., 2017b). The top hits were inserted into our reference 16S rRNA gene tree (Fig. 2A) using the Evolutionary Placement Algorithm (EPA; Berger *et al*., 2011). 76 OTUs placed adjacent to the sponge-specific sequence cluster 174, including one with 100% identity to *Ca.* N. bastadiensis. These 76 OTUs comprised, on average, 17.2% ± 7.1 (SD) of all reads obtained from *I. basta* individuals in the SMP dataset, consistent with abundances determined by FISH and qPCR (see main text). Interestingly, the *Ca*. N. bastadiensis-adjacent OTUs were also found to comprise 0.25 – 7.5% of the total reads obtained from *Ancorina alata*, *Stellata maori*, *Stellata aremaria* sampled in New Zealand and *Xestospongia exigua* sampled on the Great Barrier Reef. In *X. exigua*, one low abundance OTU (0.17% of all reads) had 100% nucleotide identity with the V4 region of the *Ca*. N. bastadiensis 16S rRNA gene. Among all environmental samples covered by the EMP, sequences highly similar or identical to *Ca*. N. bastadiensis, with abundances above 0.1%, were exclusively found in sponges. Consistent with a habitat restricted to a few sponge species, no hits above 97.6% similarity to the 16S rRNA gene of *Ca*. N. bastadiensis were detected by the Integrated Microbial NGS platform that queries most publicly available 16S rRNA gene amplicon data sets (but not the SMP dataset) (Lagkouvardos *et al*., 2016).

**Eukaryotic-like proteins (ELPs) in *Ca.* Nitrosospongia bastadiensis**. Four types of ELPs are found in *Ca*. N. bastadiensis: Proteins with tetratricopeptide repeats (TPR), the Toll-interleukin-1 receptor (TIR) -like domain PF08937 (DUF1863; Cort *et al*., 2000), immunoglobulin-like (Ig-like) domains (DUF5011; Shigeno-Nakazawa *et al*., 2016), and hyaline repeats (HYR; Callebaut *et al*., 2000). The TPR were enriched in *Ca*. N. bastadiensis when compared to other genome sequenced thaumarchaeotes, while the TIR and Ig-like domains were exclusive to *Ca*. N. bastadiensis (Fig. 3, Supporting Information Fig. S5). Of the TPR containing proteins in *Ca*. N. bastadiensis, 41 represent TPR gene families not previously detected in other thaumarchaeotes. TPR-containing proteins, which mediate protein-protein interactions in eukaryotes (Blatch and Lässle, 1999), are thought to be important for the survival of sponge symbionts in their phagocytic hosts. Two TPR-containing proteins from sponge microbiomes cloned into *E. coli* affected amoeba phagocytosis (Reynolds and Thomas, 2016), however, both proteins contained the TPR-Sel1 motif which was absent in the TPR-containing proteins from *Ca*. N. bastadiensis. TIR-like proteins have also been found in other marine sponges (Wiens *et al*., 2006; Gauthier *et al*., 2010), with these proteins known to be key mediators of the metazoan innate immune response as well as playing a role in regulating metabolic and bioenergetic pathways through modulating $NAD^+$ levels (Essuman *et al*., 2018). TIR-like proteins were expressed when sponges were subjected to bacteria-analogue lipoproteins (Wiens *et al*., 2006) and lipopolysaccharides (Wiens *et al*., 2005), which in turn caused the expression of a caspase likely involved in apoptosis and a macrophage-expressed protein, respectively. In this context, it is interesting to note that a protein encoding the DUF1863 domain from a zoonotic *Staphylococcus aureus* can decrease the survivability of mice infected by this strain (Patterson *et al*., 2014). Furthermore, the DUF1863 domain was recently shown to be a critical component of a bacterial defense system against myophages and may be involved in recognizing specific phage patterns (Doron *et al*., 2018).

An evolutionary homology between choanoflagellates and sponge choanocytes has long been speculated (Maldonado, 2005; Mah *et al*., 2014; Laundon et al., 2018). Both have diverse and abundant receptor tyrosine kinases (RTKs) (Srivastava *et al*., 2010; Miller, 2012), which are crucial components of metazoan signal transduction systems. Interestingly, choanoflagellate proteins with HYR-like domains were recently predicted to act as receptor tyrosine kinases (RTKs) (Manning *et al*., 2008). As HYR domains are structurally related to Ig and FN3 domains (Callebaut *et al*., 2000) and choanoflagellates lack the Ig domains found in many metazoan RTKs, the HYR domains may be fulfilling the role of the Ig domains in metazoan RTKs. Furthermore, the Ig-like DUF5011 domain has been found on the extracellular portion of two choanoflagellate homologues of the tyrosine kinase substrate BCAR1 (Shigeno-Nakazawa *et al*., 2016). Consequently, the numerous DUF5011 and HYR domain containing proteins of *Ca.* N. bastadiensis may be interacting with the host signaling network as *I. basta* likely harbors similar extracellular domains as part of its signal transduction and gene regulatory processes. This host-symbiont interaction is further supported by observations that (i) many *Ca.* N. bastadiensis proteins encoding these domains are predicted to be exported and (ii) five extracellular proteins of the *Ca.* N. bastadiensis exclusive gene family containing the DUF5011 domain were detected in the metaproteome (Supporting Information Table S3).

### Supplementary Experimental Procedures

**Phylogenetic analyses.** S-layer proteins were identified in all sequenced thaumarchaeota based on orthologous groups identified with Orthofinder containing members previously identified as SLPs (Li *et al*., 2018), or also classified as arCOG08647 in EggNOG version 4.5 (Huerta-Cepas *et al*., 2016). A phylogenetic reconstruction was performed with a thaumarchaeal-specific dataset with a minimal sequence length of 300 amino acids. Sequences were aligned using mafft (Katoh and Standley, 2013) and automatically trimmed using trimAl version 1.4 (Capella-Gutiérrez *et al*., 2009) and the -gappyout function. After model selection using ModelFinder (Kalyaanmoorthy *et al*., 2017) and a maximum-likelihood amino acid phylogenetic tree was generated using the model LG+F+G4 in IQ-Tree, version 1.6.2 (Nguyen *et al*., 2015) with 1,000 ultrafast bootstraps (UFBoot).

Genes encoding S08A family endopeptidases were identified in all sequenced thaumarchaeota if they contained the PF00082 (Peptidase_S8) domain when searched against the Pfam-A database. The complete sequence set was identified using thaumarchaeal amino acid sequences as individual queries for blastp searches against the Genbank nr database and only top hits containing the PF00082 domain were included. In total 277 representative sequences of the S08A family endopeptidases were used for phylogenetic analyses and sequences were trimmed according to the presence of the PF00082 domain before alignment using mafft (Katoh and Standley, 2013). ModelFinder (Kalyaanmoorthy *et al*., 2017) was used for model selection and maximum-likelihood phylogenetic analyses implemented with IQ-Tree, version 1.6.2 (Nguyen *et al*., 2015), using the LG+I+G4 model and 1,000 ultrafast bootstraps (UFBoot). Phylogenetic analyses for serpins (PF00079) were conducted in a similar fashion and 75 representative sequences were analyzed (after filtering out 46 distant homologues) using the WAG+I+G4 model and 1,000 ultrafast bootstraps.

Phylogenetic analyses were conducted on amino acid sequences of the *LivK* periplasmic branched chain amino acid transporter subunit as well as a concatenated alignment of the *LivFGHMK* operon, using the best model identified for each dataset by ModelFinder (Kalyaanmoorthy *et al*., 2017) and implemented in IQ-Tree, version 1.6.2 (Nguyen *et al*., 2015) with 1,000 ultrafast bootstraps (UFBoot). The complete sequence set was identified using individual *Ca*. N. bastadiensis *LivFGHMK* operon subunits as individual queries for blastp searches against the Genbank nr database. Top hits were included along with additional sequences demonstrated to be functional for active substrate transport within the hydrophobic amino-acid uptake transporter (HAAT) family (TCDB:3.A.14; Saier *et al*., 2009). Sequences were aligned using mafft (Katoh and Standley, 2013) and automatically trimmed using trimAl version 1.4 (Capella-Gutiérrez *et al*., 2009) and the -gappyout function. Models used for maximum-likelihood phylogenetic analyses were LG+F+G4 and LG+F+I+G4 for the LivK subunit and the concatenated *livFGHMK* operon, respectively.

**Database screening for 16S rRNA gene amplicons.** A reference set of 16S rRNA gene sequences (N=65) from sequenced thaumarchaeotal genomes and amplicons associated with sponges, including *Ca.* N. bastadensis were aligned with SINA (Pruesse *et al*., 2012) and used to construct a reference tree in RaXML (Stamatakis, 2014). Sequence tags identified by the Sponge Microbiome Project (Moitinho-Silva *et al*., 2017b) were mapped to the reference set using blastn (Camacho *et al*., 2009), requiring at least 70% alignment and 90% identity. Successfully mapped reads were then aligned to the reference alignment using SINA and placed into the reference tree using RaXML-EPA (Berger *et al*, 2011). In addition, 16S rRNA gene sequences related to *Ca.* N. bastadensis were identified in short read archive (SRA) datasets using IMNGS (www.imngs.org - Lagkouvardos *et al*., 2016) with default parameters.

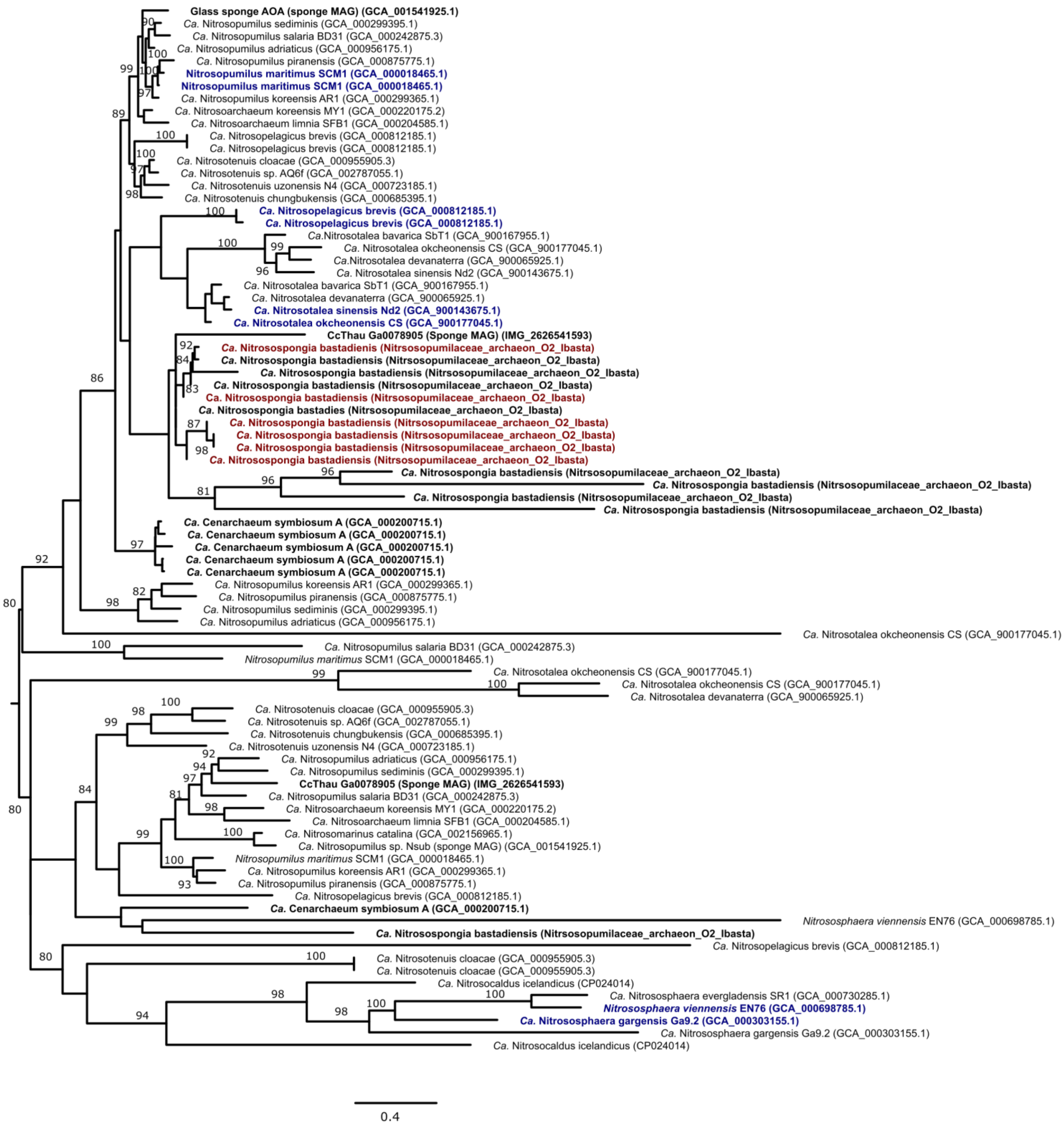
Glass sponge AOA (sponge MAG) (GCA_001541925.1)
Ca. Nitrosopumilus sediminis (GCA_000299395.1)
Ca. Nitrosopumilus salaria BD31 (GCA_000242875.3)
Ca. Nitrosopumilus adriaticus (GCA_000956175.1)
Ca. Nitrosopumilus piranensis (GCA_000875775.1)
Nitrosopumilus maritimus SCM1 (GCA_000018465.1)
Nitrosopumilus maritimus SCM1 (GCA_000018465.1)
Ca. Nitrosopumilus koreensis AR1 (GCA_000299365.1)
Ca. Nitrosoarchaeum koreensis MY1 (GCA_000220175.2)
Ca. Nitrosoarchaeum limnia SFB1 (GCA_000204585.1)
Ca. Nitrosopelagicus brevis (GCA_000812185.1)
Ca. Nitrosopelagicus brevis (GCA_000812185.1)
Ca. Nitrosotenuis cloacae (GCA_000955905.3)
Ca. Nitrosotenuis sp. AQ6f (GCA_002787055.1)
Ca. Nitrosotenuis uzonensis N4 (GCA_000723185.1)
Ca. Nitrosotenuis chungbukensis (GCA_000685395.1)
Ca. Nitrosopelagicus brevis (GCA_000812185.1)
Ca. Nitrosopelagicus brevis (GCA_000812185.1)
Ca.Nitrosotalea bavarica SbT1 (GCA_900167955.1)
Ca. Nitrosotalea okcheonensis CS (GCA_900177045.1)
Ca.Nitrosotalea devanaterra (GCA_900065925.1)
Ca. Nitrosotalea sinensis Nd2 (GCA_900143675.1)
Ca. Nitrosotalea bavarica SbT1 (GCA_900167955.1)
Ca. Nitrosotalea devanaterra (GCA_900065925.1)
Ca. Nitrosotalea sinensis Nd2 (GCA_900143675.1)
Ca. Nitrosotalea okcheonensis CS (GCA_900177045.1)
CcThau Ga0078905 (Sponge MAG) (IMG_2626541593)
Ca. Nitrosospongia bastadiensis (Nitrsosopumilaceae_archaeon_O2_Ibasta)
Ca. Nitrosospongia bastadiensis (Nitrsosopumilaceae_archaeon_O2_Ibasta)
Ca. Nitrosospongia bastadiensis (Nitrsosopumilaceae_archaeon_O2_Ibasta)
Ca. Nitrosospongia bastadiensis (Nitrsosopumilaceae_archaeon_O2_Ibasta)
Ca. Nitrosospongia bastadiensis (Nitrsosopumilaceae_archaeon_O2_Ibasta)
Ca. Nitrosospongia bastadies (Nitrsosopumilaceae_archaeon_O2_Ibasta)
Ca. Nitrosospongia bastadiensis (Nitrsosopumilaceae_archaeon_O2_Ibasta)
Ca. Nitrosospongia bastadiensis (Nitrsosopumilaceae_archaeon_O2_Ibasta)
Ca. Nitrosospongia bastadiensis (Nitrsosopumilaceae_archaeon_O2_Ibasta)
Ca. Nitrosospongia bastadiensis (Nitrsosopumilaceae_archaeon_O2_Ibasta)
Ca. Nitrosospongia bastadiensis (Nitrsosopumilaceae_archaeon_O2_Ibasta)
Ca. Nitrosospongia bastadiensis (Nitrsosopumilaceae_archaeon_O2_Ibasta)
Ca. Nitrosospongia bastadiensis (Nitrsosopumilaceae_archaeon_O2_Ibasta)
Ca. Nitrosospongia bastadiensis (Nitrsosopumilaceae_archaeon_O2_Ibasta)
Ca. Cenarchaeum symbiosum A (GCA_000200715.1)
Ca. Cenarchaeum symbiosum A (GCA_000200715.1)
Ca. Cenarchaeum symbiosum A (GCA_000200715.1)
Ca. Cenarchaeum symbiosum A (GCA_000200715.1)
Ca. Cenarchaeum symbiosum A (GCA_000200715.1)
Ca. Nitrosopumilus koreensis AR1 (GCA_000299365.1)
Ca. Nitrosopumilus piranensis (GCA_000875775.1)
Ca. Nitrosopumilus sediminis (GCA_000299395.1)
Ca. Nitrosopumilus adriaticus (GCA_000956175.1)
Ca. Nitrosotalea okcheonensis CS (GCA_900177045.1)
Ca. Nitrosopumilus salaria BD31 (GCA_000242875.3)
Nitrosopumilus maritimus SCM1 (GCA_000018465.1)
Ca. Nitrosotalea okcheonensis CS (GCA_900177045.1)
Ca. Nitrosotalea okcheonensis CS (GCA_900177045.1)
Ca. Nitrosotalea devanaterra (GCA_900065925.1)
Ca. Nitrosotenuis cloacae (GCA_000955905.3)
Ca. Nitrosotenuis sp. AQ6f (GCA_002787055.1)
Ca. Nitrosotenuis chungbukensis (GCA_000685395.1)
Ca. Nitrosotenuis uzonensis N4 (GCA_000723185.1)
Ca. Nitrosopumilus adriaticus (GCA_000956175.1)
Ca. Nitrosopumilus sediminis (GCA_000299395.1)
CcThau Ga0078905 (Sponge MAG) (IMG_2626541593)
Ca. Nitrosopumilus salaria BD31 (GCA_000242875.3)
Ca. Nitrosoarchaeum koreensis MY1 (GCA_000220175.2)
Ca. Nitrosoarchaeum limnia SFB1 (GCA_000204585.1)
Ca. Nitrosomarinus catalina (GCA_002156965.1)
Ca. Nitrosopumilus sp. Nsub (sponge MAG) (GCA_001541925.1)
Nitrosopumilus maritimus SCM1 (GCA_000018465.1)
Ca. Nitrosopumilus koreensis AR1 (GCA_000299365.1)
Ca. Nitrosopumilus piranensis (GCA_000875775.1)
Ca. Nitrosopelagicus brevis (GCA_000812185.1)
Ca. Cenarchaeum symbiosum A (GCA_000200715.1)
Nitrososphaera viennensis EN76 (GCA_000698785.1)
Ca. Nitrosospongia bastadiensis (Nitrsosopumilaceae_archaeon_O2_Ibasta)
Ca. Nitrosopelagicus brevis (GCA_000812185.1)
Ca. Nitrosotenuis cloacae (GCA_000955905.3)
Ca. Nitrosotenuis cloacae (GCA_000955905.3)
Ca. Nitrosocaldus icelandicus (CP024014)
Ca. Nitrososphaera evergladensis SR1 (GCA_000730285.1)
Nitrososphaera viennensis EN76 (GCA_000698785.1)
Ca. Nitrososphaera gargensis Ga9.2 (GCA_000303155.1)
Ca. Nitrososphaera gargensis Ga9.2 (GCA_000303155.1)
Ca. Nitrosocaldus icelandicus (CP024014)
0.4

**Figure S1**. Maximum-likelihood amino acid phylogenetic tree (after automatic model selection with IQ-Tree, version 1.6.2) of putative S-layer proteins found in all analyzed thaumarchaeal genomes (28 in total). Sponge-derived sequences are depicted in bold black, while highly expressed proteins from *Ca*. N. bastadiensis are depicted in bold red (6 in total). Highlighted in bold blue are putative S-layer proteins found to be highly expressed in previous proteomics studies (Santoro *et al*., 2015; Palatinszky *et al*., 2015; Kerou *et al*., 2016; Qin *et al*., 2017; Herbold *et al*., 2017). Values at nodes represent ultrafast bootstraps (UFBoot) with only values ≥80% shown for each branch.

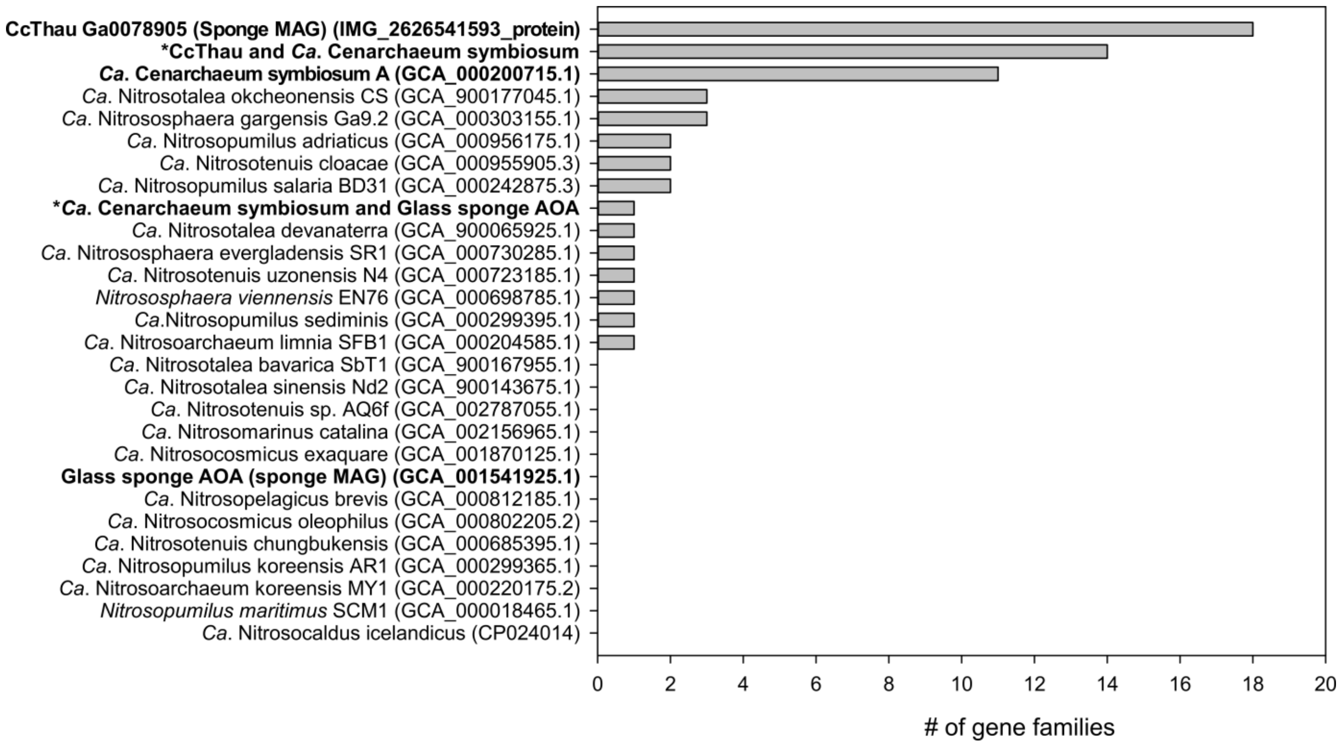

**Figure S2.** Number of gene families shared exclusively between *Ca*. N. bastadiensis and each genome-sequenced member of the *Thaumarchaeota* including the sponge thaumarchaeal symbionts, CCThau, *Ca*. C. symbiosum, and the glass sponge AOA (in bold). (*) indicates gene families shared by *Ca*. N. bastadiensis with *Ca.* C. symbiosum and CcThau exclusively, or with *Ca*. C. symbiosum and the glass sponge AOA exclusively.

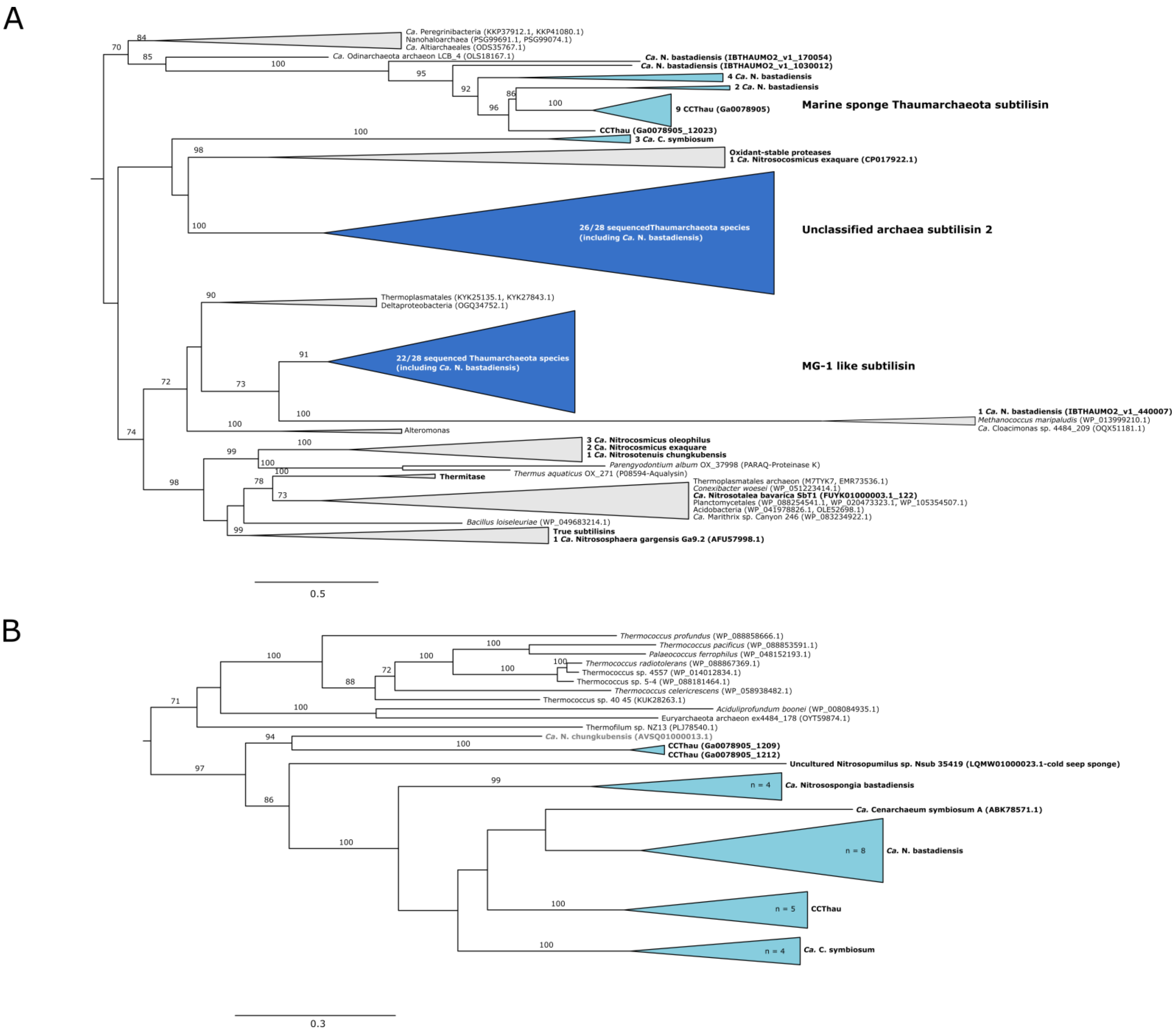

A
Ca. Peregrinibacteria (KKP37912.1, KKP41080.1)
Nanohaloarchaea (PSG99691.1, PSG99074.1)
Ca. Altiarchaeales (ODS35767.1)
Ca. Odinarchaeota archaeon LCB_4 (OLS18167.1)
Ca. N. bastadiensis (IBTHAUMO2_v1_170054)
Ca. N. bastadiensis (IBTHAUMO2_v1_1030012)
4 Ca. N. bastadiensis
2 Ca. N. bastadiensis
9 CCThau (Ga0078905)
Marine sponge Thaumarchaeota subtilisin
CCThau (Ga0078905_12023)
3 Ca. C. symbiosum
Oxidant-stable proteases
1 Ca. Nitrosocosmicus exaquare (CP017922.1)
26/28 sequencedThaumarchaeota species (including Ca. N. bastadiensis)
Unclassified archaea subtilisin 2
Thermoplasmatales (KYK25135.1, KYK27843.1)
Deltaproteobacteria (OGQ34752.1)
22/28 sequenced Thaumarchaeota species (including Ca. N. bastadiensis)
MG-1 like subtilisin
1 Ca. N. bastadiensis (IBTHAUMO2_v1_440007)
Methanococcus maripaludis (WP_013999210.1)
Ca. Cloacimonas sp. 4484_209 (OQX51181.1)
Alteromonas
3 Ca. Nitrocosmicus oleophilus
2 Ca. Nitrocosmicus exaquare
1 Ca. Nitrosotenuis chungkubensis
Parengyodontium album OX_37998 (PARAQ-Proteinase K)
Thermus aquaticus OX_271 (P08594-Aqualysin)
Thermitase
Thermoplasmatales archaeon (M7TYK7, EMR73536.1)
Conexibacter woesei (WP_051223414.1)
Ca. Nitrosotalea bavarica SbT1 (FUYK01000003.1_122)
Planctomycetales (WP_088254541.1, WP_020473323.1, WP_105354507.1)
Acidobacteria (WP_041978826.1, OLE52698.1)
Ca. Marithrix sp. Canyon 246 (WP_083234922.1)
Bacillus loiseleuriae (WP_049683214.1)
True subtilisins
1 Ca. Nitrososphaera gargensis Ga9.2 (AFU57998.1)
0.5
B
Thermococcus profundus (WP_088858666.1)
Thermococcus pacificus (WP_088853591.1)
Palaeococcus ferrophilus (WP_048152193.1)
Thermococcus radiotolerans (WP_088867369.1)
Thermococcus sp. 4557 (WP_014012834.1)
Thermococcus sp. 5-4 (WP_088181464.1)
Thermococcus celericrescens (WP_058938482.1)
Thermococcus sp. 40 45 (KUK28263.1)
Aciduliprofundum boonei (WP_008084935.1)
Euryarchaeota archaeon ex4484_178 (OYT59874.1)
Thermofilum sp. NZ13 (PLJ78540.1)
Ca. N. chungkubensis (AVSQ01000013.1)
CCThau (Ga0078905_1209)
CCThau (Ga0078905_1212)
Uncultured Nitrosopumilus sp. Nsub 35419 (LQMW01000023.1-cold seep sponge)
n = 4
Ca. Nitrosospongia bastadiensis
Ca. Cenarchaeum symbiosum A (ABK78571.1)
n = 8
Ca. N. bastadiensis
n = 5
CCThau
n = 4
Ca. C. symbiosum
0.3

**Figure S3.** Maximum-likelihood amino acid phylogenetic trees (after automatic model selection with IQ-Tree, version 1.6.2) of (A) S08A family endopeptidases and (B) serine protease inhibitors (serpins). Color coding donates the degree of homology among all sequenced *Thaumarchaeota*: light blue – shared exclusively among thaumarchaeal sponge symbionts; dark blue – ubiquitously found in *Thaumarchaeota*. In (A), 6 out of 8 *Ca*. N. bastadiensis S08A family endopeptidases, within the "Marine sponge Thaumarchaeota subtilisin" clade, are predicted to be exported. On the other hand, the majority of the ubiquitous "Unclassified archaea subtilisin 2" (24/27) and all except one of the "MG-1 like subtilisin" (naming convention after Li *et al*., 2015) are predicted to be membrane anchored S08A endopeptidases. In (B), 3 of the 15 serpins found in *Ca*. N. bastadiensis were excluded from phylogenetic analyses due to truncated length. Values at nodes represent ultrafast bootstraps (UFBoot) with only values ≥80% shown for each branch.

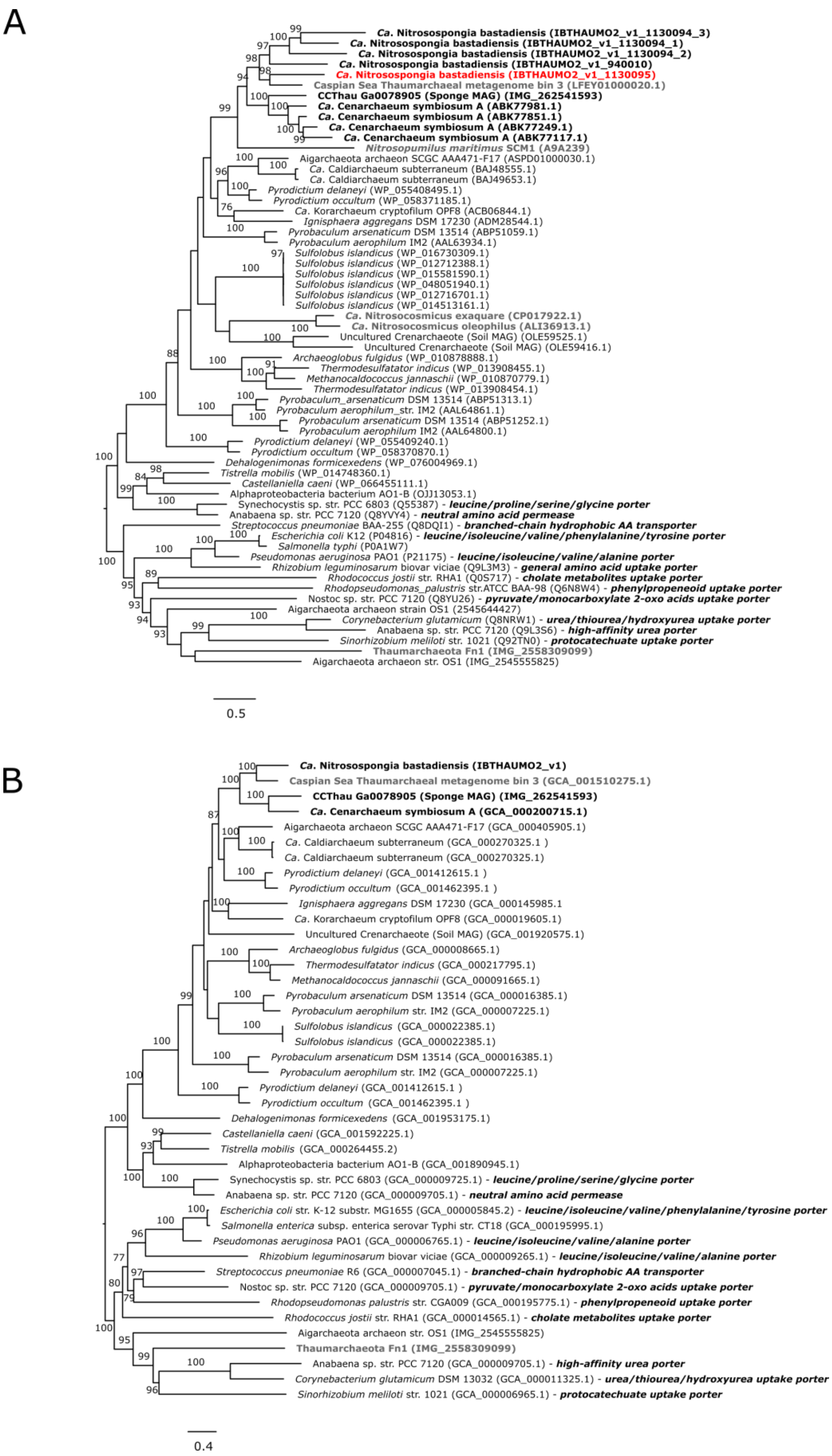
A
Ca. Nitrosospongia bastadiensis (IBTHAUMO2_v1_1130094_3)
Ca. Nitrosospongia bastadiensis (IBTHAUMO2_v1_1130094_1)
Ca. Nitrosospongia bastadiensis (IBTHAUMO2_v1_1130094_2)
Ca. Nitrosospongia bastadiensis (IBTHAUMO2_v1_940010)
Ca. Nitrosospongia bastadiensis (IBTHAUMO2_v1_1130095)
Caspian Sea Thaumarchaeal metagenome bin 3 (LFEY01000020.1)
CCThau Ga0078905 (Sponge MAG) (IMG_262541593)
Ca. Cenarchaeum symbiosum A (ABK77981.1)
Ca. Cenarchaeum symbiosum A (ABK77851.1)
Ca. Cenarchaeum symbiosum A (ABK77249.1)
Ca. Cenarchaeum symbiosum A (ABK77117.1)
Nitrosopumilus maritimus SCM1 (A9A239)
Aigarchaeota archaeon SCGC AAA471-F17 (ASPD01000030.1)
Ca. Caldiarchaeum subterraneum (BAJ48555.1)
Ca. Caldiarchaeum subterraneum (BAJ49653.1)
Pyrodictium delaneyi (WP_055408495.1)
Pyrodictium occultum (WP_058371185.1)
Ca. Korarchaeum cryptofilum OPF8 (ACB06844.1)
Ignisphaera aggregans DSM 17230 (ADM28544.1)
Pyrobaculum arsenaticum DSM 13514 (ABP51059.1)
Pyrobaculum aerophilum IM2 (AAL63934.1)
Sulfolobus islandicus (WP_016730309.1)
Sulfolobus islandicus (WP_012712388.1)
Sulfolobus islandicus (WP_015581590.1)
Sulfolobus islandicus (WP_048051940.1)
Sulfolobus islandicus (WP_012716701.1)
Sulfolobus islandicus (WP_014513161.1)
Ca. Nitrosocosmicus exaquare (CP017922.1)
Ca. Nitrosocosmicus oleophilus (ALI36913.1)
Uncultured Crenarchaeote (Soil MAG) (OLE59525.1)
Uncultured Crenarchaeote (Soil MAG) (OLE59416.1)
Archaeoglobus fulgidus (WP_010878888.1)
Thermodesulfatator indicus (WP_013908455.1)
Methanocaldococcus jannaschii (WP_010870779.1)
Thermodesulfatator indicus (WP_013908454.1)
Pyrobaculum_arsenaticum DSM 13514 (ABP51313.1)
Pyrobaculum aerophilum_str. IM2 (AAL64861.1)
Pyrobaculum arsenaticum DSM 13514 (ABP51252.1)
Pyrobaculum aerophilum IM2 (AAL64800.1)
Pyrodictium delaneyi (WP_055409240.1)
Pyrodictium occultum (WP_058370870.1)
Dehalogenimonas formicexedens (WP_076004969.1)
Tistrella mobilis (WP_014748360.1)
Castellaniella caeni (WP_066455111.1)
Alphaproteobacteria bacterium AO1-B (OJJ13053.1)
Synechocystis sp. str. PCC 6803 (Q55387) - leucine/proline/serine/glycine porter
Anabaena sp. str. PCC 7120 (Q8YVY4) - neutral amino acid permease
Streptococcus pneumoniae BAA-255 (Q8DQI1) - branched-chain hydrophobic AA transporter
Escherichia coli K12 (P04816) - leucine/isoleucine/valine/phenylalanine/tyrosine porter
Salmonella typhi (P0A1W7)
Pseudomonas aeruginosa PAO1 (P21175) - leucine/isoleucine/valine/alanine porter
Rhizobium leguminosarum biovar viciae (Q9L3M3) - general amino acid uptake porter
Rhodococcus jostii str. RHA1 (Q0S717) - cholate metabolites uptake porter
Rhodopseudomonas_palustris str.ATCC BAA-98 (Q6N8W4) - phenylpropeneoid uptake porter
Nostoc sp. str. PCC 7120 (Q8YU26) - pyruvate/monocarboxylate 2-oxo acids uptake porter
Aigarchaeota archaeon strain OS1 (2545644427)
Corynebacterium glutamicum (Q8NRW1) - urea/thiourea/hydroxyurea uptake porter
Anabaena sp. str. PCC 7120 (Q9L3S6) - high-affinity urea porter
Sinorhizobium meliloti str. 1021 (Q92TN0) - protocatechuate uptake porter
Thaumarchaeota Fn1 (IMG_2558309099)
Aigarchaeota archaeon str. OS1 (IMG_2545555825)
0.5
B
Ca. Nitrosospongia bastadiensis (IBTHAUMO2_v1)
Caspian Sea Thaumarchaeal metagenome bin 3 (GCA_001510275.1)
CCThau Ga0078905 (Sponge MAG) (IMG_262541593)
Ca. Cenarchaeum symbiosum A (GCA_000200715.1)
Aigarchaeota archaeon SCGC AAA471-F17 (GCA_000405905.1)
Ca. Caldiarchaeum subterraneum (GCA_000270325.1 )
Ca. Caldiarchaeum subterraneum (GCA_000270325.1)
Pyrodictium delaneyi (GCA_001412615.1 )
Pyrodictium occultum (GCA_001462395.1 )
Ignisphaera aggregans DSM 17230 (GCA_000145985.1
Ca. Korarchaeum cryptofilum OPF8 (GCA_000019605.1)
Uncultured Crenarchaeote (Soil MAG) (GCA_001920575.1)
Archaeoglobus fulgidus (GCA_000008665.1)
Thermodesulfatator indicus (GCA_000217795.1)
Methanocaldococcus jannaschii (GCA_000091665.1)
Pyrobaculum arsenaticum DSM 13514 (GCA_000016385.1)
Pyrobaculum aerophilum str. IM2 (GCA_000007225.1)
Sulfolobus islandicus (GCA_000022385.1)
Sulfolobus islandicus (GCA_000022385.1)
Pyrobaculum arsenaticum DSM 13514 (GCA_000016385.1)
Pyrobaculum aerophilum str. IM2 (GCA_000007225.1)
Pyrodictium delaneyi (GCA_001412615.1 )
Pyrodictium occultum (GCA_001462395.1 )
Dehalogenimonas formicexedens (GCA_001953175.1)
Castellaniella caeni (GCA_001592225.1)
Tistrella mobilis (GCA_000264455.2)
Alphaproteobacteria bacterium AO1-B (GCA_001890945.1)
Synechocystis sp. str. PCC 6803 (GCA_000009725.1) - leucine/proline/serine/glycine porter
Anabaena sp. str. PCC 7120 (GCA_000009705.1) - neutral amino acid permease
Escherichia coli str. K-12 substr. MG1655 (GCA_000005845.2) - leucine/isoleucine/valine/phenylalanine/tyrosine porter
Salmonella enterica subsp. enterica serovar Typhi str. CT18 (GCA_000195995.1)
Pseudomonas aeruginosa PAO1 (GCA_000006765.1) - leucine/isoleucine/valine/alanine porter
Rhizobium leguminosarum biovar viciae (GCA_000009265.1) - leucine/isoleucine/valine/alanine porter
Streptococcus pneumoniae R6 (GCA_000007045.1) - branched-chain hydrophobic AA transporter
Nostoc sp. str. PCC 7120 (GCA_000009705.1) - pyruvate/monocarboxylate 2-oxo acids uptake porter
Rhodopseudomonas palustris str. CGA009 (GCA_000195775.1) - phenylpropeneoid uptake porter
Rhodococcus jostii str. RHA1 (GCA_000014565.1) - cholate metabolites uptake porter
Aigarchaeota archaeon str. OS1 (IMG_2545555825)
Thaumarchaeota Fn1 (IMG_2558309099)
Anabaena sp. str. PCC 7120 (GCA_000009705.1) - high-affinity urea porter
Corynebacterium glutamicum DSM 13032 (GCA_000011325.1) - urea/thiourea/hydroxyurea uptake porter
Sinorhizobium meliloti str. 1021 (GCA_000006965.1) - protocatechuate uptake porter
0.4

**Figure S4**. Maximum-likelihood amino acid phylogenetic trees (after automatic model selection with IQ-Tree, version 1.6.2) for the periplasmic subunit (A) *LivK* and for the concatenated (B) *LivFGHMK* operon. *LivK* was found to be highly expressed (expressed copy in red) in the *Ca*. N. bastadiensis proteome and can occur in multiple copies among *LivFGHMK* encoding microorganisms. *Ca*. N. bastadiensis has two *LivK* genes and a gene containing three fused *LivK* domains and the predicted proteins from these five *LivK* genes/domains were included in the phylogenetic analysis. *LivK* is also found in some other thaumarchaeotes (bold grey), but these lack the *LivFGHM* genes necessary for branched-chain amino acid transport. In a few cases in the concatenated *LivFGHMK* tree, multiple operons were present in a given genome. For both trees, sponge thaumarchaeal sequences are highlighted in bold black while other thaumarchaeal sequences are highlighted in bold grey. Sequences from organisms where the specific transport functions have been identified, have those specific functions annotated. These annotations are taken from information collated in the Transporter Classification database (Saier *et al*., 2009). Values at nodes represent ultrafast bootstraps (UFBoot) with only values ≥80% shown for each branch.

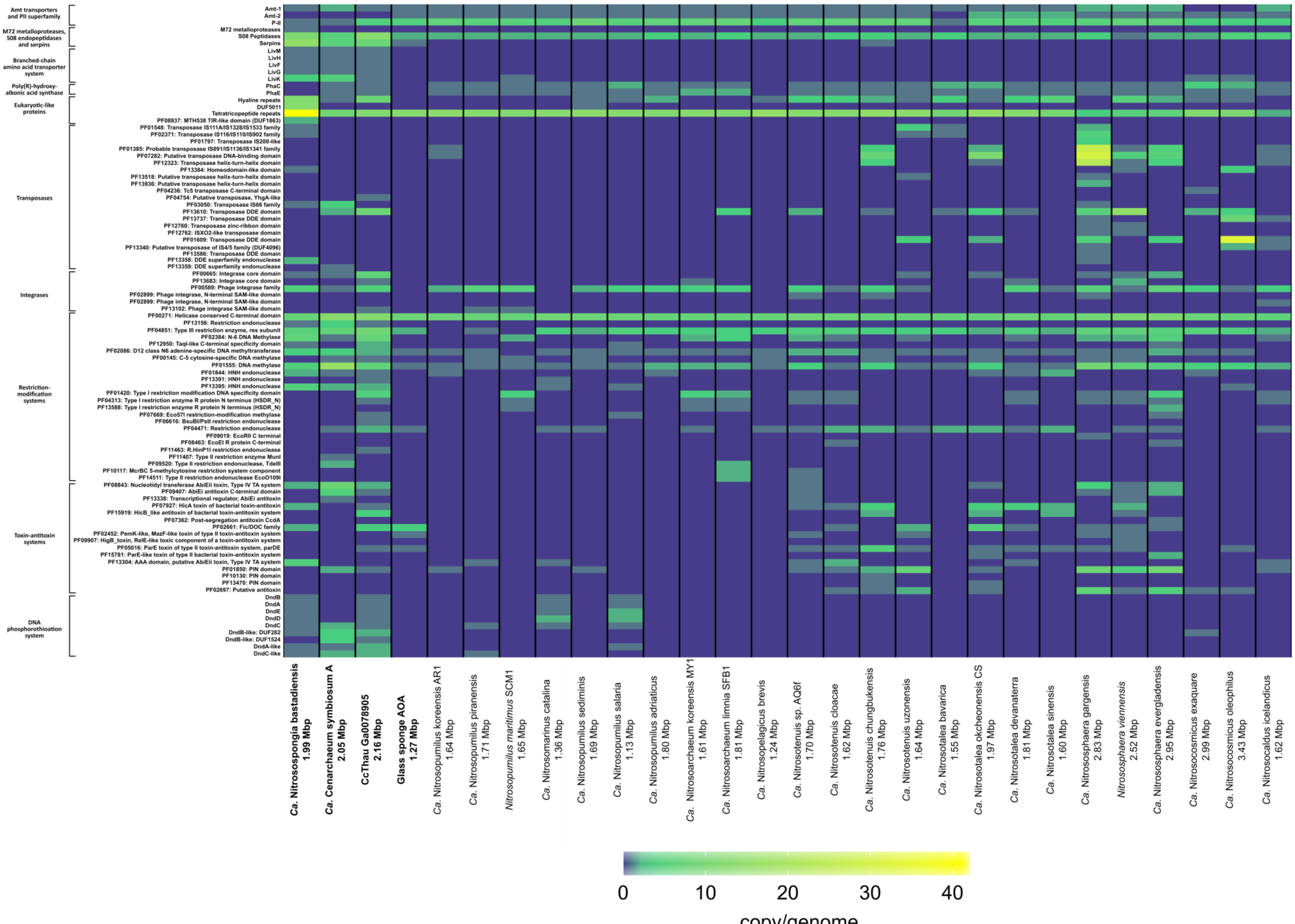
Amt transporters and PII superfamily
M72 metalloproteases, S08 endopeptidases and serpins
Branched-chain amino acid transporter system
Poly(R)-hydroxy-alkonic acid synthase
Eukaryotic-like proteins
Transposases
Integrases
Restriction-modification systems
Toxin-antitoxin systems
DNA phosphorothioation system
Amt-1
Amt-2
P-II
M72 metalloproteases
S08 Peptidases
Serpins
LivM
LivH
LivF
LivG
LivK
PhaC
PhaE
Hyaline repeats
DUF5011
Tetratricopeptide repeats
PF08937: MTH538 TIR-like domain (DUF1863)
PF01548: Transposase IS111A/IS1328/IS1533 family
PF02371: Transposase IS116/IS110/IS902 family
PF01797: Transposase IS200-like
PF01385: Probable transposase IS891/IS1136/IS1341 family
PF07282: Putative transposase DNA-binding domain
PF12323: Transposase helix-turn-helix domain
PF13384: Homeodomain-like domain
PF13518: Putative transposase helix-turn-helix domain
PF13936: Putative transposase helix-turn-helix domain
PF04236: Tc5 transposase C-terminal domain
PF04754: Putative transposase, YhgA-like
PF03050: Transposase IS66 family
PF13610: Transposase DDE domain
PF13737: Transposase DDE domain
PF12760: Transposase zinc-ribbon domain
PF12762: ISXO2-like transposase domain
PF01609: Transposase DDE domain
PF13340: Putative transposase of IS4/5 family (DUF4096)
PF13586: Transposase DDE domain
PF13358: DDE superfamily endonuclease
PF13359: DDE superfamily endonuclease
PF00665: Integrase core domain
PF13683: Integrase core domain
PF00589: Phage integrase family
PF02899: Phage integrase, N-terminal SAM-like domain
PF02899: Phage integrase, N-terminal SAM-like domain
PF13102: Phage integrase SAM-like domain
PF00271: Helicase conserved C-terminal domain
PF13156: Restriction endonuclease
PF04851: Type III restriction enzyme, res subunit
PF02384: N-6 DNA Methylase
PF12950: TaqI-like C-terminal specificity domain
PF02086: D12 class N6 adenine-specific DNA methyltransferase
PF00145: C-5 cytosine-specific DNA methylase
PF01555: DNA methylase
PF01844: HNH endonuclease
PF13391: HNH endonuclease
PF13395: HNH endonuclease
PF01420: Type I restriction modification DNA specificity domain
PF04313: Type I restriction enzyme R protein N terminus (HSDR_N)
PF13588: Type I restriction enzyme R protein N terminus (HSDR_N)
PF07669: Eco57I restriction-modification methylase
PF06616: BsuBI/PstI restriction endonuclease
PF04471: Restriction endonuclease
PF09019: EcoRII C terminal
PF08463: EcoEI R protein C-terminal
PF11463: R.HinP1I restriction endonuclease
PF11407: Type II restriction enzyme MunI
PF09520: Type II restriction endonuclease, TdeIII
PF10117: McrBC 5-methylcytosine restriction system component
PF14511: Type II restriction endonuclease EcoO109I
PF08843: Nucleotidyl transferase AbiEii toxin, Type IV TA system
PF09407: AbiEi antitoxin C-terminal domain
PF13338: Transcriptional regulator, AbiEi antitoxin
PF07927: HicA toxin of bacterial toxin-antitoxin
PF15919: HicB_like antitoxin of bacterial toxin-antitoxin system
PF07362: Post-segregation antitoxin CcdA
PF02661: Fic/DOC family
PF02452: PemK-like, MazF-like toxin of type II toxin-antitoxin system
PF09907: HigB_toxin, RelE-like toxic component of a toxin-antitoxin system
PF05016: ParE toxin of type II toxin-antitoxin system, parDE
PF15781: ParE-like toxin of type II bacterial toxin-antitoxin system
PF13304: AAA domain, putative AbiEii toxin, Type IV TA system
PF01850: PIN domain
PF10130: PIN domain
PF13470: PIN domain
PF02697: Putative antitoxin
DndB
DndA
DndE
DndD
DndC
DndB-like: DUF262
DndB-like: DUF1524
DndA-like
DndC-like
Ca. Nitrosospongia bastadiensis 1.99 Mbp
Ca. Cenarchaeum symbiosum A 2.05 Mbp
CcThau Ga0078905 2.16 Mbp
Glass sponge AOA 1.27 Mbp
Ca. Nitrosopumilus koreensis AR1 1.64 Mbp
Ca. Nitrosopumilus piranensis 1.71 Mbp
Nitrosopumilus maritimus SCM1 1.65 Mbp
Ca. Nitrosomarinus catalina 1.36 Mbp
Ca. Nitrosopumilus sediminis 1.69 Mbp
Ca. Nitrosopumilus salaria 1.13 Mbp
Ca. Nitrosopumilus adriaticus 1.80 Mbp
Ca. Nitrosoarchaeum koreensis MY1 1.61 Mbp
Ca. Nitrosoarchaeum limnia SFB1 1.81 Mbp
Ca. Nitrosopelagicus brevis 1.24 Mbp
Ca. Nitrosotenuis sp. AQ6f 1.70 Mbp
Ca. Nitrosotenuis cloacae 1.62 Mbp
Ca. Nitrosotenuis chungbukensis 1.76 Mbp
Ca. Nitrosotenuis uzonensis 1.64 Mbp
Ca. Nitrosotalea bavarica 1.55 Mbp
Ca. Nitrosotalea okcheonensis CS 1.97 Mbp
Ca. Nitrosotalea devanaterra 1.81 Mbp
Ca. Nitrosotalea sinensis 1.60 Mbp
Ca. Nitrososphaera gargensis 2.83 Mbp
Nitrososphaera viennensis 2.52 Mbp
Ca. Nitrososphaera evergladensis 2.95 Mbp
Ca. Nitrosocosmicus exaquare 2.99 Mbp
Ca. Nitrosocosmicus oleophilus 3.43 Mbp
Ca. Nitrosocaldus icelandicus 1.62 Mbp
0
10
20
30
40
copy/genome

**Figure S5.** Heat map showing the distribution and gene copy number per genome of selected genes, gene classes and PFAM annotations among genome-sequenced AOA. The color scale ranges from 0 (dark blue) to 40 (yellow) and indicates copies per genome. Sponge-derived genomes start on the left and are depicted in bold, followed by members of *Ca*. Nitrosopumilaceae, *Ca*. Nitrosotenuaceae, *Ca*. Nitrosotaleale, the *Nitrososphaerales*, and *Ca*. Nitrosocaldales, respectively. Genome sizes for each member are listed below each name.

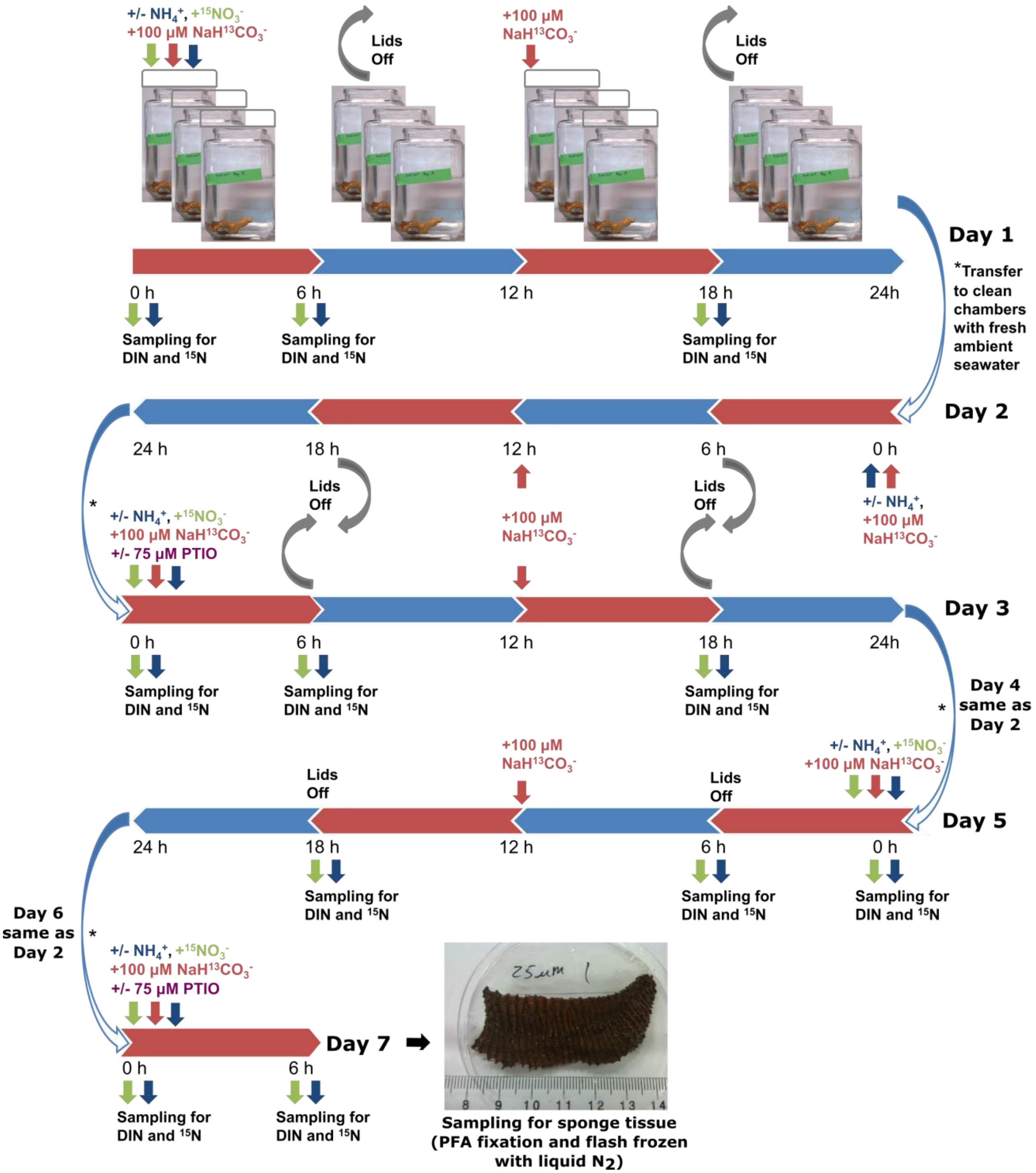

**Figure S6.** Experimental design for the *I. basta* holobiont nitrification incubations. The intermittently closed setup was employed to avoid oxygen depletion while minimizing loss of $^{13}$C-labeled $HCO_3^-$. Green and blue arrows denote samples used for the calculation of net and gross nitrification rates, respectively. Days 4, 5 and 6 were recovery days for those incubations to which PTIO was added.

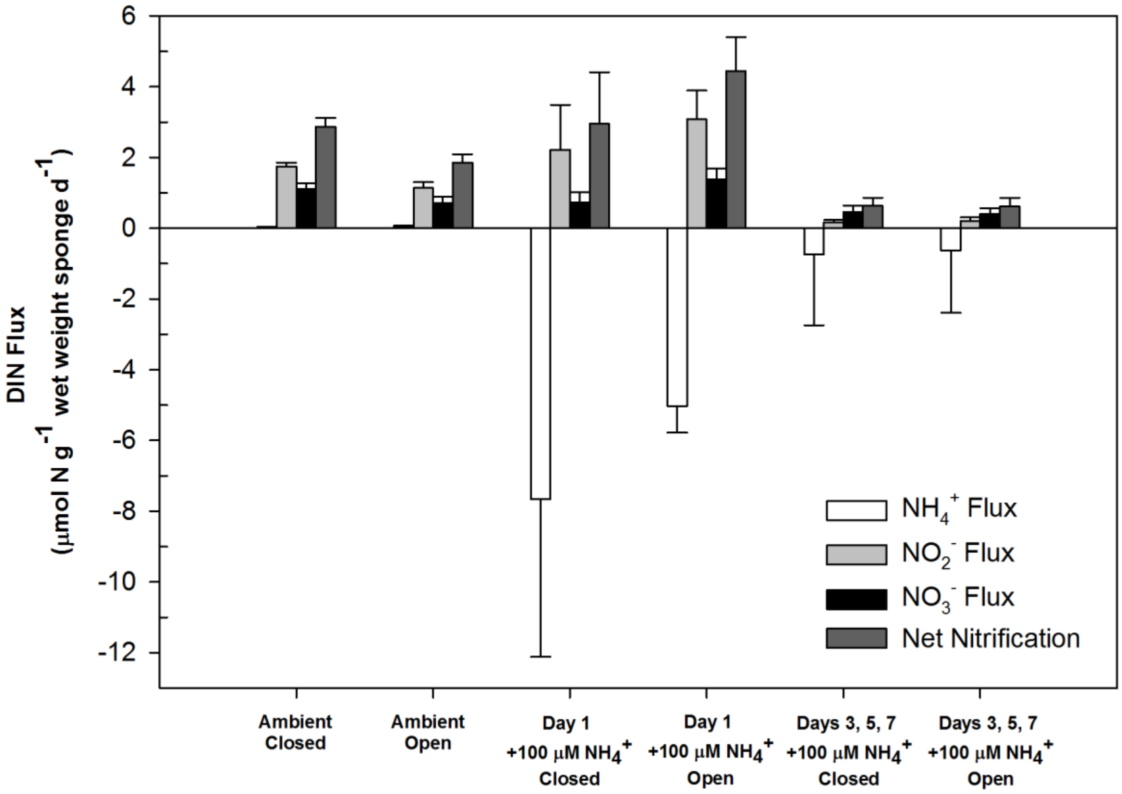

**Figure S7.** Comparison of average DIN ($NH_4^+$, $NO_3^-$, $NO_2^-$) flux and net nitrification rates exhibited by the *I. basta* holobiont in multiple-day incubations between intermittently closed (Supplementary Figure 1) and completely open aquaria at ambient conditions and at 100 µM $NH_4^+$. Pair-wise comparisons of individual DIN species flux and net nitrification between open and intermittently closed aquaria, revealed significant differences in net $NO_2^-$ flux and net nitrification between ambient treatments (both: $p < 0.01$, Mann-Whitney U-test).

**Table S1**. Overview of genome binning parameters, statistics and associated metadata for the *Ca.* N. bastadiensis genome bin.

| Bin ID | IBThaumO2 |
|---|---|
| Analysis project type | metagenome-assembled genome (MAG) |
| Taxa_id | 16S rRNA and multi-marker phylogenetics |
| Assembly software | Spades |
| Annotation | MaGe |
| Genome Quality | High Quality Draft |
| Completeness (%) | 99.03 |
| Contamination (%) | 0.97 |
| Completeness/Contamination Software | CheckM |
| Number of contigs | 113 |
| 16S rRNA gene recovered | yes |
| 16S rRNA gene recovery software | Rnammer / MaGe |
| Number of standard tRNAs extracted | 20 |
| tRNA extraction software | tRNA-scan / MaGe |
| Binning software | metabat2 |
| Binning parameters | kmer |
| Genome size (Mbp) | 1.99 |
| N50 (bp) | 35099 |
| Longest contig (bp) | 135,585 |
| Average contig length (bp) | 17,669.70 |
| GC content (%) | 64.8 |
| Protein coding sequences | 2,342 |
| rRNAs | 3 |
| tRNAs | 42 |

**Table S2.** Presence of genes of interest (see Figure 3) in the two *Ca.* N. bastadiensis MAGs (obtained from two *I. basta* individuals by Illumina and 454 pyrosequencing, respectively) as determined by BLASTp.

| | qseqid | sseqid | pident | length | qlen | slen | evalue | bitscore | query match coverage | subject match coverage |
|---|---|---|---|---|---|---|---|---|---|---|
| M72 metalloproteases | IBTHAUMO2_v1_880008 | IBTHAUMv1_16080007 | 64.242 | 660 | 2028 | 2440 | 0 | 729 | 32.54 | 26.76 |
| S08 Peptidases | IBTHAUMO2_v1_1030012 | IBTHAUMv1_12290002 | 91.578 | 5141 | 5956 | 6201 | 0 | 9250 | 86.32 | 82.26 |
| | IBTHAUMO2_v1_150018 | IBTHAUMv1_17130004 | 99.286 | 560 | 908 | 611 | 0 | 1133 | 61.67 | 91.49 |
| | IBTHAUMO2_v1_160002 | IBTHAUMv1_1950001 | 100 | 659 | 1181 | 1349 | 0 | 1355 | 55.80 | 48.78 |
| | IBTHAUMO2_v1_170054 | IBTHAUMv1_3990003 | 99.646 | 1695 | 1695 | 1695 | 0 | 3440 | 100.00 | 99.94 |
| | IBTHAUMO2_v1_220007 | IBTHAUMv1_13830026 | 99.542 | 655 | 655 | 655 | 0 | 1257 | 100.00 | 99.85 |
| | IBTHAUMO2_v1_250004 | IBTHAUMv1_4210010 | 65.672 | 268 | 303 | 1619 | 1.16E-98 | 315 | 88.45 | 16.24 |
| | IBTHAUMO2_v1_440007 | IBTHAUMv1_14780018 | 99.856 | 696 | 699 | 696 | 0 | 1429 | 99.57 | 99.86 |
| | IBTHAUMO2_v1_470031 | IBTHAUMv1_15480003 | 99.794 | 1459 | 1459 | 1459 | 0 | 2938 | 100.00 | 99.93 |
| | IBTHAUMO2_v1_470037 | IBTHAUMv1_15480010 | 99.446 | 1264 | 1264 | 1266 | 0 | 2486 | 100.00 | 99.76 |
| | IBTHAUMO2_v1_600001 | IBTHAUMv1_1950001 | 89.893 | 1029 | 1563 | 1349 | 0 | 1843 | 65.83 | 76.20 |
| | IBTHAUMO2_v1_810001 | IBTHAUMv1_16080007 | 92.872 | 2413 | 2385 | 2440 | 0 | 4403 | 101.17 | 98.11 |
| Serpins | IBTHAUMO2_v1_1040003 | IBTHAUMv1_16780011 | 78.208 | 413 | 472 | 412 | 0 | 620 | 87.50 | 99.76 |
| | IBTHAUMO2_v1_150002 | IBTHAUMv1_9590002 | 100 | 412 | 412 | 412 | 0 | 835 | 100.00 | 99.76 |
| | IBTHAUMO2_v1_150006 | IBTHAUMv1_2680003 | 99.589 | 487 | 487 | 487 | 0 | 979 | 100.00 | 99.79 |
| | IBTHAUMO2_v1_150019 | IBTHAUMv1_17130003 | 100 | 468 | 501 | 499 | 0 | 943 | 93.41 | 93.59 |
| | IBTHAUMO2_v1_160001 | IBTHAUMv1_15480013 | 98.14 | 645 | 646 | 649 | 0 | 1268 | 99.85 | 99.23 |
| | IBTHAUMO2_v1_250003 | IBTHAUMv1_9590002 | 83.846 | 390 | 413 | 412 | 0 | 665 | 94.43 | 92.48 |
| | IBTHAUMO2_v1_250005 | IBTHAUMv1_3270002 | 50.427 | 117 | 107 | 377 | 4.53E-25 | 94.4 | 109.35 | 30.77 |
| | IBTHAUMO2_v1_470041 | IBTHAUMv1_17030009 | 92.308 | 273 | 389 | 649 | 0 | 521 | 70.18 | 41.91 |
| | IBTHAUMO2_v1_470042 | IBTHAUMv1_16080008 | 74.297 | 498 | 498 | 498 | 0 | 667 | 100.00 | 99.20 |
| | IBTHAUMO2_v1_530010 | IBTHAUMv1_3270002 | 97.706 | 218 | 240 | 377 | 3.43E-150 | 421 | 90.83 | 57.56 |
| | IBTHAUMO2_v1_530011 | IBTHAUMv1_3270002 | 98.052 | 154 | 154 | 377 | 7.63E-104 | 299 | 100.00 | 40.58 |
| | IBTHAUMO2_v1_630004 | IBTHAUMv1_4530014 | 100 | 419 | 419 | 419 | 0 | 853 | 100.00 | 99.76 |
| | IBTHAUMO2_v1_740001 | IBTHAUMv1_2680003 | 95.833 | 72 | 74 | 487 | 2.60E-36 | 124 | 97.30 | 14.58 |
| | IBTHAUMO2_v1_810003 | IBTHAUMv1_16080004 | 100 | 474 | 474 | 474 | 0 | 970 | 100.00 | 99.79 |
| | IBTHAUMO2_v1_880009 | IBTHAUMv1_16080008 | 100 | 498 | 498 | 498 | 0 | 1014 | 100.00 | 99.80 |
| LivM | IBTHAUMO2_v1_1130089 | IBTHAUMv1_13040008 | 99.717 | 353 | 353 | 353 | 0 | 677 | 100.00 | 99.72 |
| LivH | IBTHAUMO2_v1_1130090 | IBTHAUMv1_13040007 | 100 | 306 | 306 | 306 | 0 | 587 | 100.00 | 99.67 |
| LivF | IBTHAUMO2_v1_1130093 | IBTHAUMv1_13040004 | 100 | 238 | 238 | 238 | 1.80E-172 | 471 | 100.00 | 99.58 |
| LivG | IBTHAUMO2_v1_1130092 | IBTHAUMv1_13040005 | 99.209 | 253 | 269 | 253 | 0 | 496 | 94.05 | 99.60 |

| | | | | | | | | | | |
|---|---|---|---|---|---|---|---|---|---|---|
| LivK | IBTHAUMO2_v1_1130094 | IBTHAUMv1_13040003 | 99.688 | 642 | 1149 | 642 | 0 | 1236 | 55.87 | 99.84 |
| LivK | IBTHAUMO2_v1_1130095 | IBTHAUMv1_16960002 | 93.738 | 527 | 527 | 511 | 0 | 993 | 100.00 | 99.80 |
| LivK | IBTHAUMO2_v1_940010 | IBTHAUMv1_330001 | 100 | 314 | 402 | 314 | 0 | 622 | 78.11 | 99.68 |
| DndB | IBTHAUMO2_v1_810011 | IBTHAUMv1_190016 | 44.688 | 320 | 362 | 319 | 2.21E-98 | 292 | 88.40 | 98.43 |
| DndA | IBTHAUMO2_v1_880013 | IBTHAUMv1_140010 | 70.712 | 379 | 391 | 388 | 0 | 566 | 96.93 | 97.42 |
| DndE | IBTHAUMO2_v1_880014 | IBTHAUMv1_140009 | 39.655 | 116 | 123 | 126 | 1.18E-32 | 108 | 94.31 | 91.27 |
| DndD | IBTHAUMO2_v1_880015 | IBTHAUMv1_190014 | 34.074 | 675 | 675 | 660 | 1.80E-107 | 337 | 100.00 | 99.55 |
| DndC | IBTHAUMO2_v1_880016 | IBTHAUMv1_140006 | 56.592 | 493 | 473 | 502 | 0 | 576 | 104.23 | 98.01 |
| DndB-like: DUF262 | IBTHAUMO2_v1_1050033 | IBTHAUMv1_380002 | 27.711 | 166 | 425 | 344 | 2.67E-10 | 58.9 | 39.06 | 44.48 |
| DndA-like | IBTHAUMO2_v1_690025 | IBTHAUMv1_3820004 | 99.482 | 386 | 386 | 386 | 0 | 772 | 100.00 | 99.74 |
| DndC-like | IBTHAUMO2_v1_690027 | IBTHAUMv1_17710003 | 100 | 280 | 280 | 280 | 0 | 582 | 100.00 | 99.64 |
| PF08937: MTH538 TIR-like | IBTHAUMO2_v1_460006 | IBTHAUMv1_14780023 | 100 | 130 | 130 | 170 | 7.13E-94 | 265 | 100.00 | 75.88 |
| | IBTHAUMO2_v1_620008 | IBTHAUMv1_14750015 | 100 | 172 | 172 | 172 | 3.99E-126 | 349 | 100.00 | 99.42 |
| Hyaline repeats | IBTHAUMO2_v1_150018 | IBTHAUMv1_17130004 | 99.286 | 560 | 908 | 611 | 0 | 1133 | 61.67 | 91.49 |
| | IBTHAUMO2_v1_1050019 | IBTHAUMv1_14910004 | 99.774 | 442 | 442 | 442 | 0 | 880 | 100.00 | 99.77 |
| | IBTHAUMO2_v1_1050020 | IBTHAUMv1_14910005 | 75.036 | 701 | 705 | 702 | 0 | 988 | 99.43 | 99.15 |
| | IBTHAUMO2_v1_1110002 | IBTHAUMv1_460002 | 44.798 | 471 | 491 | 487 | 8.48E-124 | 368 | 95.93 | 96.51 |
| | IBTHAUMO2_v1_1110003 | IBTHAUMv1_15140009 | 49.099 | 444 | 443 | 452 | 2.55E-124 | 366 | 100.23 | 97.35 |
| | IBTHAUMO2_v1_220002 | IBTHAUMv1_150003 | 86.853 | 715 | 712 | 715 | 0 | 1217 | 100.42 | 99.86 |
| | IBTHAUMO2_v1_450016 | IBTHAUMv1_3920005 | 97.511 | 442 | 630 | 442 | 0 | 862 | 70.16 | 99.77 |
| | IBTHAUMO2_v1_470043 | IBTHAUMv1_16080007 | 92.531 | 241 | 237 | 2440 | 1.02E-125 | 389 | 101.69 | 9.80 |
| | IBTHAUMO2_v1_720042 | IBTHAUMv1_14780010 | 48.347 | 242 | 268 | 742 | 1.76E-63 | 209 | 90.30 | 31.81 |
| | IBTHAUMO2_v1_810001 | IBTHAUMv1_16080007 | 92.872 | 2413 | 2385 | 2440 | 0 | 4403 | 101.17 | 98.11 |
| | IBTHAUMO2_v1_880008 | IBTHAUMv1_16080007 | 64.242 | 660 | 2028 | 2440 | 0 | 729 | 32.54 | 26.76 |
| PF16403: DUF5011 | IBTHAUMO2_v1_1030012 | IBTHAUMv1_12290002 | 91.578 | 5141 | 5956 | 6201 | 0 | 9250 | 86.32 | 82.26 |
| | IBTHAUMO2_v1_170054 | IBTHAUMv1_3990003 | 99.646 | 1695 | 1695 | 1695 | 0 | 3440 | 100.00 | 99.94 |
| | IBTHAUMO2_v1_20012 | IBTHAUMv1_11360007 | 95.513 | 936 | 1651 | 962 | 0 | 1722 | 56.69 | 97.19 |
| | IBTHAUMO2_v1_240080 | IBTHAUMv1_11360001 | 98.649 | 2073 | 2073 | 2073 | 0 | 4182 | 100.00 | 99.95 |
| | IBTHAUMO2_v1_290009 | IBTHAUMv1_13830036 | 99.644 | 1966 | 1966 | 1966 | 0 | 3962 | 100.00 | 99.95 |
| | IBTHAUMO2_v1_320050 | IBTHAUMv1_2050001 | 97.937 | 1115 | 1328 | 1117 | 0 | 2196 | 83.96 | 99.73 |
| | IBTHAUMO2_v1_460001 | IBTHAUMv1_14780028 | 91.978 | 1633 | 1773 | 4551 | 0 | 2982 | 92.10 | 35.62 |
| | IBTHAUMO2_v1_470031 | IBTHAUMv1_15480003 | 99.794 | 1459 | 1459 | 1459 | 0 | 2938 | 100.00 | 99.93 |
| | IBTHAUMO2_v1_590001 | IBTHAUMv1_20280001 | 98.699 | 615 | 648 | 1427 | 0 | 1256 | 94.91 | 43.03 |
| | IBTHAUMO2_v1_590078 | IBTHAUMv1_13120027 | 99.522 | 837 | 1040 | 848 | 0 | 1659 | 80.48 | 98.58 |

| | | | | | | | | | | |
|---|---|---|---|---|---|---|---|---|---|---|
| | IBTHAUMO2_v1_620022 | IBTHAUMv1_12370012 | 98.862 | 1845 | 1845 | 2170 | 0 | 3714 | 100.00 | 84.98 |
| | IBTHAUMO2_v1_700001 | IBTHAUMv1_18570002 | 76.798 | 2836 | 3095 | 8426 | 0 | 4177 | 91.63 | 33.54 |
| | IBTHAUMO2_v1_990001 | IBTHAUMv1_4410003 | 99.731 | 1116 | 1116 | 11182 | 0 | 2271 | 100.00 | 9.97 |
| Tetratricopeptide repeats | IBTHAUMO2_v1_20005 | IBTHAUMv1_4210007 | 100 | 542 | 542 | 561 | 0 | 1096 | 100.00 | 96.43 |
| | IBTHAUMO2_v1_260017 | IBTHAUMv1_1700002 | 99.743 | 389 | 389 | 394 | 0 | 786 | 100.00 | 98.48 |
| | IBTHAUMO2_v1_450009 | IBTHAUMv1_15960007 | 98.947 | 665 | 665 | 665 | 0 | 1304 | 100.00 | 99.85 |
| | IBTHAUMO2_v1_590071 | IBTHAUMv1_13120019 | 98.704 | 463 | 463 | 463 | 0 | 926 | 100.00 | 99.78 |
| | IBTHAUMO2_v1_690013 | IBTHAUMv1_11050005 | 100 | 236 | 236 | 236 | 4.75E-174 | 475 | 100.00 | 99.58 |
| | IBTHAUMO2_v1_250010 | IBTHAUMv1_1040006 | 97.196 | 214 | 214 | 214 | 2.07E-147 | 406 | 100.00 | 99.53 |
| | IBTHAUMO2_v1_590041 | IBTHAUMv1_4080009 | 98.78 | 738 | 738 | 738 | 0 | 1435 | 100.00 | 99.86 |
| | IBTHAUMO2_v1_730001 | IBTHAUMv1_5300001 | 97.778 | 90 | 123 | 144 | 3.63E-61 | 181 | 73.17 | 61.81 |
| | IBTHAUMO2_v1_1130054 | IBTHAUMv1_13940010 | 100 | 262 | 262 | 262 | 0 | 514 | 100.00 | 99.62 |
| | IBTHAUMO2_v1_10010 | IBTHAUMv1_1840004 | 33.738 | 412 | 416 | 754 | 3.80E-61 | 208 | 99.04 | 54.24 |
| | IBTHAUMO2_v1_1080005 | IBTHAUMv1_16190004 | 99.153 | 118 | 118 | 118 | 1.76E-83 | 237 | 100.00 | 99.15 |
| | IBTHAUMO2_v1_1120015 | IBTHAUMv1_5290001 | 59.541 | 566 | 738 | 565 | 0 | 629 | 76.69 | 99.47 |
| | IBTHAUMO2_v1_170044 | IBTHAUMv1_3990013 | 99.425 | 174 | 174 | 174 | 1.18E-117 | 328 | 100.00 | 99.43 |
| | IBTHAUMO2_v1_200001 | IBTHAUMv1_15480023 | 99.693 | 326 | 417 | 326 | 0 | 643 | 78.18 | 99.69 |
| | IBTHAUMO2_v1_20006 | IBTHAUMv1_4210006 | 100 | 495 | 495 | 495 | 0 | 997 | 100.00 | 99.80 |
| | IBTHAUMO2_v1_260040 | IBTHAUMv1_1140003 | 99.635 | 274 | 274 | 274 | 0 | 539 | 100.00 | 99.64 |
| | IBTHAUMO2_v1_330008 | IBTHAUMv1_17060002 | 94.872 | 429 | 457 | 443 | 0 | 805 | 93.87 | 96.61 |
| | IBTHAUMO2_v1_410008 | IBTHAUMv1_13120019 | 45.02 | 251 | 274 | 463 | 1.19E-66 | 212 | 91.61 | 52.92 |
| | IBTHAUMO2_v1_410009 | IBTHAUMv1_1610001 | 94.574 | 387 | 399 | 394 | 0 | 697 | 96.99 | 97.97 |
| | IBTHAUMO2_v1_440013 | IBTHAUMv1_14780004 | 86.288 | 598 | 598 | 598 | 0 | 1035 | 100.00 | 99.83 |
| | IBTHAUMO2_v1_450076 | IBTHAUMv1_15360002 | 100 | 299 | 299 | 299 | 0 | 563 | 100.00 | 99.67 |
| | IBTHAUMO2_v1_480001 | IBTHAUMv1_3180004 | 100 | 339 | 339 | 339 | 0 | 669 | 100.00 | 99.71 |
| | IBTHAUMO2_v1_480002 | IBTHAUMv1_3180005 | 99.76 | 416 | 434 | 416 | 0 | 793 | 95.85 | 99.76 |
| | IBTHAUMO2_v1_510001 | IBTHAUMv1_2270002 | 100 | 279 | 279 | 534 | 0 | 553 | 100.00 | 52.06 |
| | IBTHAUMO2_v1_690014 | IBTHAUMv1_11050004 | 98.305 | 236 | 236 | 236 | 9.73E-172 | 469 | 100.00 | 99.58 |
| | IBTHAUMO2_v1_650002 | IBTHAUMv1_20210002 | 100 | 262 | 262 | 262 | 0 | 516 | 100.00 | 99.62 |
| | IBTHAUMO2_v1_700005 | IBTHAUMv1_21200002 | 96.032 | 252 | 256 | 256 | 4.84E-171 | 469 | 98.44 | 98.05 |
| | IBTHAUMO2_v1_750012 | IBTHAUMv1_2270002 | 100 | 241 | 243 | 534 | 4.86E-176 | 492 | 99.18 | 44.94 |
| | IBTHAUMO2_v1_790015 | IBTHAUMv1_15480026 | 99.77 | 435 | 435 | 435 | 0 | 852 | 100.00 | 99.77 |
| | IBTHAUMO2_v1_880020 | IBTHAUMv1_2270002 | 99.17 | 241 | 268 | 534 | 1.46E-174 | 489 | 89.93 | 44.94 |
| | IBTHAUMO2_v1_940002 | IBTHAUMv1_4830003 | 96.811 | 439 | 439 | 439 | 0 | 838 | 100.00 | 99.77 |

| | | | | | | | | | | |
|---|---|---|---|---|---|---|---|---|---|---|
| | IBTHAUMO2_v1_990018 | IBTHAUMv1_16760014 | 100 | 140 | 140 | 140 | 2.46E-92 | 261 | 100.00 | 99.29 |
| | IBTHAUMO2_v1_200002 | IBTHAUMv1_15480022 | 93.885 | 278 | 283 | 286 | 4.26E-178 | 489 | 98.23 | 96.85 |
| | IBTHAUMO2_v1_20010 | IBTHAUMv1_4210003 | 98.396 | 187 | 187 | 187 | 1.12E-126 | 352 | 100.00 | 99.47 |
| | IBTHAUMO2_v1_720008 | IBTHAUMv1_4130013 | 100 | 127 | 127 | 127 | 1.21E-88 | 250 | 100.00 | 99.21 |
| | IBTHAUMO2_v1_380029 | IBTHAUMv1_11240003 | 100 | 309 | 309 | 309 | 0 | 606 | 100.00 | 99.68 |
| | IBTHAUMO2_v1_590042 | IBTHAUMv1_4080008 | 99.286 | 420 | 423 | 423 | 0 | 845 | 99.29 | 99.05 |
| | IBTHAUMO2_v1_990017 | IBTHAUMv1_16760013 | 100 | 68 | 68 | 68 | 7.21E-43 | 130 | 100.00 | 98.53 |
| | IBTHAUMO2_v1_480009 | IBTHAUMv1_90003 | 100 | 197 | 197 | 237 | 5.88E-143 | 395 | 100.00 | 82.70 |
| | IBTHAUMO2_v1_700003 | IBTHAUMv1_20360001 | 100 | 204 | 204 | 252 | 2.91E-149 | 412 | 100.00 | 80.56 |
| Transposases | IBTHAUMO2_v1_1050022 | IBTHAUMv1_8580062 | 99.457 | 368 | 368 | 368 | 0 | 733 | 100.00 | 99.73 |
| | IBTHAUMO2_v1_560020 | IBTHAUMv1_23780003 | 100 | 120 | 121 | 144 | 2.04E-87 | 248 | 99.17 | 82.64 |
| | IBTHAUMO2_v1_1100079 | IBTHAUMv1_4230002 | 100 | 172 | 172 | 354 | 6.57E-127 | 358 | 100.00 | 48.31 |
| | IBTHAUMO2_v1_1110001 | IBTHAUMv1_14670003 | 100 | 147 | 147 | 339 | 1.13E-106 | 305 | 100.00 | 43.07 |
| | IBTHAUMO2_v1_20011 | IBTHAUMv1_18170003 | 100 | 119 | 119 | 454 | 1.75E-82 | 246 | 100.00 | 25.99 |
| Integrases | IBTHAUMO2_v1_240171 | IBTHAUMv1_2410001 | 99.74 | 385 | 557 | 387 | 0 | 801 | 69.12 | 99.22 |
| | IBTHAUMO2_v1_450024 | IBTHAUMv1_16200001 | 100 | 454 | 454 | 472 | 0 | 939 | 100.00 | 95.97 |
| | IBTHAUMO2_v1_730023 | IBTHAUMv1_1500005 | 98.063 | 413 | 448 | 424 | 0 | 840 | 92.19 | 96.93 |
| | IBTHAUMO2_v1_880010 | IBTHAUMv1_16080009 | 99.787 | 469 | 469 | 469 | 0 | 967 | 100.00 | 99.79 |
| | IBTHAUMO2_v1_780006 | IBTHAUMv1_11050003 | 99.674 | 307 | 307 | 343 | 0 | 628 | 100.00 | 89.21 |
| Restriction-modification | IBTHAUMO2_v1_1050026 | IBTHAUMv1_1120001 | 25.738 | 237 | 927 | 640 | 2.47E-06 | 48.5 | 25.57 | 31.41 |
| systems | IBTHAUMO2_v1_1130086 | IBTHAUMv1_13350012 | 25.895 | 475 | 612 | 1038 | 8.26E-32 | 129 | 77.61 | 45.47 |
| | IBTHAUMO2_v1_260066 | IBTHAUMv1_14890001 | 99.05 | 421 | 1333 | 586 | 0 | 863 | 31.58 | 71.67 |
| | IBTHAUMO2_v1_700006 | IBTHAUMv1_4230001 | 100 | 887 | 953 | 889 | 0 | 1845 | 93.07 | 99.66 |
| | IBTHAUMO2_v1_880011 | IBTHAUMv1_13350012 | 20.659 | 334 | 908 | 1038 | 2.49E-06 | 48.9 | 36.78 | 31.21 |
| | IBTHAUMO2_v1_890002 | IBTHAUMv1_13350012 | 99.711 | 1038 | 1037 | 1038 | 0 | 2118 | 100.10 | 99.90 |
| | IBTHAUMO2_v1_240074 | IBTHAUMv1_15140004 | 99.184 | 490 | 490 | 490 | 0 | 975 | 100.00 | 99.80 |
| | IBTHAUMO2_v1_240151 | IBTHAUMv1_16180005 | 100 | 574 | 574 | 582 | 0 | 1178 | 100.00 | 98.45 |
| | IBTHAUMO2_v1_280001 | IBTHAUMv1_11110026 | 98.119 | 638 | 675 | 729 | 0 | 1280 | 94.52 | 87.38 |
| | IBTHAUMO2_v1_380002 | IBTHAUMv1_11240027 | 99.42 | 862 | 862 | 862 | 0 | 1708 | 100.00 | 99.88 |
| | IBTHAUMO2_v1_730017 | IBTHAUMv1_3930004 | 99.719 | 711 | 711 | 711 | 0 | 1407 | 100.00 | 99.86 |
| | IBTHAUMO2_v1_800038 | IBTHAUMv1_3940004 | 100 | 648 | 648 | 648 | 0 | 1320 | 100.00 | 99.85 |
| | IBTHAUMO2_v1_260067 | IBTHAUMv1_980001 | 97.107 | 242 | 271 | 273 | 1.02E-174 | 480 | 89.30 | 88.28 |
| | IBTHAUMO2_v1_590086 | IBTHAUMv1_1620001 | 35 | 60 | 854 | 856 | 3.10E-01 | 32 | 7.03 | 6.89 |
| | IBTHAUMO2_v1_790008 | IBTHAUMv1_19180002 | 90.868 | 438 | 439 | 442 | 0 | 823 | 99.77 | 98.87 |

| | | | | | | | | | | |
|---|---|---|---|---|---|---|---|---|---|---|
| | IBTHAUMO2_v1_390013 | IBTHAUMv1_14530025 | 99.27 | 274 | 290 | 283 | 0 | 559 | 94.48 | 96.47 |
| | IBTHAUMO2_v1_590085 | IBTHAUMv1_9460007 | 29.957 | 464 | 633 | 608 | 3.18E-52 | 187 | 73.30 | 63.65 |
| | IBTHAUMO2_v1_590087 | IBTHAUMv1_9460007 | 35.641 | 390 | 594 | 608 | 7.75E-62 | 213 | 65.66 | 59.38 |
| | IBTHAUMO2_v1_740003 | IBTHAUMv1_70001 | 66 | 450 | 446 | 477 | 0 | 639 | 100.90 | 94.13 |
| | IBTHAUMO2_v1_320013 | IBTHAUMv1_3540002 | 100 | 281 | 281 | 281 | 0 | 561 | 100.00 | 99.64 |
| | IBTHAUMO2_v1_390007 | IBTHAUMv1_14530029 | 98.233 | 283 | 299 | 287 | 0 | 558 | 94.65 | 98.26 |
| | IBTHAUMO2_v1_770023 | IBTHAUMv1_19010011 | 100 | 269 | 269 | 269 | 0 | 558 | 100.00 | 99.63 |
| | IBTHAUMO2_v1_240101 | IBTHAUMv1_10080009 | 99.569 | 464 | 510 | 467 | 0 | 925 | 90.98 | 99.14 |
| | IBTHAUMO2_v1_590091 | IBTHAUMv1_7470001 | 93.216 | 398 | 462 | 399 | 0 | 763 | 86.15 | 99.50 |
| | IBTHAUMO2_v1_690028 | IBTHAUMv1_3820007 | 30.699 | 329 | 377 | 379 | 8.32E-33 | 124 | 87.27 | 79.68 |
| | IBTHAUMO2_v1_810008 | IBTHAUMv1_7470001 | 45.013 | 391 | 474 | 399 | 1.25E-98 | 300 | 82.49 | 94.99 |
| Toxin: antitoxin systems | IBTHAUMO2_v1_1130060 | IBTHAUMv1_13940004 | 100 | 83 | 83 | 83 | 2.73E-58 | 170 | 100.00 | 98.80 |
| | IBTHAUMO2_v1_990027 | IBTHAUMv1_13940004 | 34.667 | 75 | 86 | 83 | 2.83E-11 | 52 | 87.21 | 87.95 |
| | IBTHAUMO2_v1_570004 | IBTHAUMv1_840006 | 98.214 | 112 | 112 | 112 | 5.96E-77 | 220 | 100.00 | 99.11 |
| | IBTHAUMO2_v1_590016 | IBTHAUMv1_23220003 | 98.942 | 378 | 378 | 378 | 0 | 740 | 100.00 | 99.74 |
| | IBTHAUMO2_v1_330015 | IBTHAUMv1_1010002 | 99.091 | 330 | 377 | 330 | 0 | 655 | 87.53 | 99.70 |
| | IBTHAUMO2_v1_520002 | IBTHAUMv1_4560007 | 100 | 376 | 377 | 376 | 0 | 768 | 99.73 | 99.73 |
| | IBTHAUMO2_v1_210021 | IBTHAUMv1_6980001 | 100 | 294 | 402 | 294 | 0 | 598 | 73.13 | 99.66 |
| | IBTHAUMO2_v1_440006 | IBTHAUMv1_15080002 | 99.785 | 465 | 465 | 465 | 0 | 900 | 100.00 | 99.78 |
| | IBTHAUMO2_v1_870005 | IBTHAUMv1_5820001 | 100 | 214 | 447 | 216 | 1.21E-150 | 424 | 47.87 | 98.61 |
| | IBTHAUMO2_v1_940001 | IBTHAUMv1_4550011 | 100 | 418 | 419 | 465 | 0 | 835 | 99.76 | 89.68 |

**Table S3**. Expressed proteins assigned to the *I. basta* thaumarchaeote MAG ordered by NSAF value. Proteins that are encoded exclusively by *Ca*. N. bastadiensis among the AOA are labeled in orange. Proteins encoded by all AOA are labeled in blue.

| Accession | Gene | Description | AA length | NSAF | OG Family | Presence in 454 Thaumarchaeal bin | | | | Distribution of OGs** | Notes |
|---|---|---|---|---|---|---|---|---|---|---|---|
| | | | | | | Accession | maxLrap * | minLrap * | % BLASTp hit | | |
| IBTHAUMO2_v1_1100065 | | 4Fe-4S ferredoxin | 100 | **9.23%** | OG0000047 | IBTHAUMv1_12290026 | 1 | 1 | 100 | Core | |
| IBTHAUMO2_v1_240128 | *tuf* | Elongation factor 1-alpha | 432 | **5.90%** | OG0000743 | IBTHAUMv1_310002 | 1 | 1 | 100 | Core | |
| IBTHAUMO2_v1_720032 | *nirK* | putative nitrite reductase, copper-dependent | 468 | **3.95%** | OG0000247 | IBTHAUMv1_10010001 | 0.46795 | 0.99095 | 96.35 | not in *Ca*. N. islandicus and *Ca*. C. symbisoum | |
| IBTHAUMO2_v1_320009 | *amoB* | putative archaeal ammonia monooxygenase subunit B | 189 | **3.49%** | OG0000306 | IBTHAUMv1_240002 | 1 | 1 | 100 | Core | |
| IBTHAUMO2_v1_950017 | | conserved protein of unknown function | 355 | **3.34%** | OG0000097 | IBTHAUMv1_4200003 | 0.58028 | 0.78626 | 89.32 | | |
| IBTHAUMO2_v1_890024 | *ths* | Thermosome subunit | 546 | **3.22%** | OG0000022 | IBTHAUMv1_7040001 | 1 | 1 | 99.82 | Core | |
| IBTHAUMO2_v1_240148 | | Zn-dependent oxidoreductase | 356 | **3.09%** | OG0000035 | IBTHAUMv1_16180002 | 1 | 1 | 100 | Core | |
| IBTHAUMO2_v1_1130066 | | conserved protein of unknown function | 86 | **3.07%** | OG0001007 | IBTHAUMv1_620009 | 1 | 1 | 100 | | |
| IBTHAUMO2_v1_1100022 | *ths* | Thermosome subunit | 566 | **3.03%** | OG0000022 | IBTHAUMv1_4580008 | 1 | 1 | 99.82 | Core | |
| IBTHAUMO2_v1_510038 | *rrp41* | Exosome complex component Rrp41 | 243 | **2.53%** | OG0000833 | IBTHAUMv1_1960004 | 1 | 1 | 100 | Core | |
| IBTHAUMO2_v1_250013 | *trxA* | Thioredoxin 1 | 108 | **2.44%** | OG0000373 | IBTHAUMv1_11050023 | 1 | 1 | 100 | Core | |
| IBTHAUMO2_v1_170030 | *atpA* | V-type ATP synthase alpha chain | 589 | **2.39%** | OG0000512 | IBTHAUMv1_3990024 | 1 | 1 | 100 | Core | |
| IBTHAUMO2_v1_320010 | *amoC* | Ammonia monooxygenase/methane monooxygenase, subunit C | 187 | **2.35%** | OG0000065 | IBTHAUMv1_4820001 | 0.92118 | 1 | 100 | Core | |
| IBTHAUMO2_v1_1070007 | | Band 7 protein | 285 | **2.31%** | OG0001373 | IBTHAUMv1_16780016 | 0.8807 | 1 | 100 | | |
| IBTHAUMO2_v1_1110035 | *psmA* | Proteasome subunit alpha 2 | 240 | **1.83%** | OG0000072 | IBTHAUMv1_1190006 | 1 | 1 | 100 | Core | |
| IBTHAUMO2_v1_270002 | | protein of unknown function | 986 | **1.74%** | OG0001421 | IBTHAUMv1_16090001 | 0.91481 | 1 | 100 | All 3 sponge symbionts, 2 *Ca*. Nitrosotaleales, 1 *Ca*. N. brevis | s-layer protein family |
| IBTHAUMO2_v1_780001 | | exported protein of unknown function | 511 | **1.63%** | OG0001421 | IBTHAUMv1_2030001 | 0.48711 | 0.99804 | 100 | All 3 sponge symbionts, 2 *Ca*. Nitrosotaleales, 1 *Ca*. N. brevis | s-layer protein family |
| IBTHAUMO2_v1_450045 | *sufC* | FeS assembly ATPase SufC | 256 | **1.55%** | OG0000878 | IBTHAUMv1_9550001 | 1 | 1 | 100 | Core | |
| IBTHAUMO2_v1_1070037 | | 4Fe-4S ferredoxin | 181 | **1.46%** | OG0000029 | IBTHAUMv1_3930011 | 1 | 1 | 100 | Core | |
| IBTHAUMO2_v1_1130095 | *livK* | ABC-type branched-chain amino acid transport system, periplasmic component (modular protein) | 526 | **1.42%** | OG0001679 | IBTHAUMv1_16960002 | 1 | 1.03137 | 93.73 | All 3 sponge symbionts, *N. maritimus* and both Nitrosocosmicus spp. | |
| IBTHAUMO2_v1_660006 | | exported protein of unknown function | 510 | **1.38%** | OG0001421 | IBTHAUMv1_2030001 | 0.48902 | 1.00392 | 82.62 | All 3 sponge symbionts, 2 *Ca*. Nitrosotaleales, 1 *Ca*. N. brevis | s-layer protein family |
| IBTHAUMO2_v1_1030029 | | putative archaeal aspartate aminotransferase | 383 | **1.38%** | OG0000543 | IBTHAUMv1_3950021 | 1 | 1 | 100 | Core | |
| IBTHAUMO2_v1_240080 | | exported protein of unknown function | 2072 | **1.32%** | OG0001466 | IBTHAUMv1_11360001 | 1 | 1 | 98.65 | *Ca*. N. bastadiensis unique | DUF5011 domain-containing |
| IBTHAUMO2_v1_990028 | *psmA* | Proteasome subunit alpha | 244 | **1.26%** | OG0000072 | IBTHAUMv1_15240015 | 1 | 1 | 100 | Core | |
| IBTHAUMO2_v1_60001 | | exported protein of unknown function | 1110 | **1.23%** | OG0001421 | IBTHAUMv1_15240010 | 0.71754 | 1.05045 | 62.78 | All 3 sponge symbionts, 2 *Ca*. Nitrosotaleales, 1 *Ca*. N. brevis | s-layer protein family |

| | | | | | | | | | | | |
|---|---|---|---|---|---|---|---|---|---|---|---|
| IBTHAUMO2_v1_320005 | *rps15* | 30S ribosomal protein S15 | 149 | **1.18%** | OG0000795 | IBTHAUMv1_240006 | 1 | 1 | 99.33 | Core | |
| IBTHAUMO2_v1_170031 | *atpB* | V-type ATP synthase beta chain | 456 | **1.16%** | OG0000511 | IBTHAUMv1_3990023 | 1 | 1 | 100 | Core | |
| IBTHAUMO2_v1_980016 | *rpoA* | DNA-directed RNA polymerase subunit A'' | 1263 | **1.11%** | OG0000600 | IBTHAUMv1_12910016 | 1 | 1 | 100 | Core | |
| IBTHAUMO2_v1_450036 | | protein of unknown function | 82 | **1.07%** | OG0002396 | IBTHAUMv1_15190002 | 1 | 1 | 90.24 | Only shared b/n *Ca*. N. bastadiensis, *N. maritimus*, and *Ca*. N. gargensis | |
| IBTHAUMO2_v1_950026 | *ftnB* | putative ferritin-2 | 168 | **1.05%** | OG0001406 | IBTHAUMv1_3240002 | 1 | 1 | 99.4 | Sporadic distribution - seems to be concentrated in the Nitrosopumiliacae | |
| IBTHAUMO2_v1_530013 | *albA* | DNA/RNA-binding protein Alba (modular protein) | 170 | **1.03%** | OG0000027 | IBTHAUMv1_3270004 | 1 | 1 | 100 | Core | |
| IBTHAUMO2_v1_980006 | *mdh* | Malate dehydrogenase | 302 | **1.02%** | OG0000569 | IBTHAUMv1_4300002 | 0.91515 | 1 | 98.34 | Core | |
| IBTHAUMO2_v1_240105 | | conserved exported protein of unknown function | 451 | **0.97%** | OG0000059 | IBTHAUMv1_1870005 | 0.55654 | 1 | 100 | | |
| IBTHAUMO2_v1_770040 | | Cyclase/dehydrase | 198 | **0.89%** | OG0000265 | IBTHAUMv1_20630002 | 1 | 1 | 100 | Core | |
| IBTHAUMO2_v1_980019 | | Ribosomal protein L7Ae/L30e/S12e/Gadd45 | 104 | **0.85%** | OG0000601 | IBTHAUMv1_12970010 | 1 | 1 | 100 | Core | |
| IBTHAUMO2_v1_1110009 | | conserved exported protein of unknown function | 313 | **0.84%** | OG0000016 | IBTHAUMv1_920001 | 0.85942 | 1 | 99.26 | Core | |
| IBTHAUMO2_v1_1110029 | *erpA* | Iron-sulfur cluster insertion protein ErpA 1 | 116 | **0.76%** | OG0000603 | IBTHAUMv1_18460002 | 1 | 1 | 100 | Core | |
| IBTHAUMO2_v1_1110033 | *gdhA* | Glutamate dehydrogenase | 424 | **0.73%** | OG0000604 | IBTHAUMv1_18460006 | 0.80189 | 1 | 100 | Core | |
| IBTHAUMO2_v1_990049 | *rpl6* | 50S ribosomal protein L6 | 182 | **0.72%** | OG0000737 | IBTHAUMv1_4350015 | 1 | 1 | 99.45 | Core | |
| IBTHAUMO2_v1_720011 | | exported protein of unknown function | 1424 | **0.71%** | OG0001466 | IBTHAUMv1_4130010 | 1 | 1 | 99.16 | *Ca.* N. bastadiensis unique | |
| IBTHAUMO2_v1_470003 | *tbp* | TATA-box-binding protein | 187 | **0.71%** | OG0000031 | IBTHAUMv1_12200005 | 1 | 1 | 100 | Core | |
| IBTHAUMO2_v1_240046 | | conserved protein of unknown function | 130 | **0.68%** | OG0000007 | IBTHAUMv1_2100004 | 1 | 1 | 100 | only missing in *Ca*.N. islandicus | |
| IBTHAUMO2_v1_980022 | *rps7* | 30S ribosomal protein S7 | 199 | **0.66%** | OG0000837 | IBTHAUMv1_13050003 | 1 | 1 | 100 | Core | |
| IBTHAUMO2_v1_220008 | *sodA* | Superoxide dismutase [Mn] | 205 | **0.64%** | OG0000084 | IBTHAUMv1_13830025 | 1 | 1 | 100 | Core | |
| IBTHAUMO2_v1_260024 | | Universal stress protein (UspA domain-containing protein) | 141 | **0.62%** | OG0000011 | IBTHAUMv1_1640004 | 1 | 1 | 100 | only missing in *Ca*.N. islandicus | important for oxidative and acid stress |
| IBTHAUMO2_v1_260038 | | Cupin 2 conserved barrel domain protein | 142 | **0.62%** | OG0001023 | IBTHAUMv1_1140005 | 1 | 1 | 99.3 | only missing in *Ca*.N. islandicus | |
| IBTHAUMO2_v1_340008 | *dnaK* | Chaperone protein DnaK | 503 | **0.61%** | OG0000335 | IBTHAUMv1_11380008 | 0.75262 | 0.99801 | 100 | Core | |
| IBTHAUMO2_v1_210018 | *cofD* | LPPG:FO 2-phospho-L-lactate transferase | 306 | **0.57%** | OG0000676 | IBTHAUMv1_4620004 | 1 | 1 | 100 | Core | F420 biosynthesis |
| IBTHAUMO2_v1_1110028 | *dnaG* | DNA primase DnaG | 386 | **0.57%** | OG0000693 | IBTHAUMv1_2640008 | 0.80829 | 1 | 99.68 | Core | |
| IBTHAUMO2_v1_1030019 | *ppi* | putative peptidyl-prolyl cis-trans isomerase | 158 | **0.56%** | OG0000324 | IBTHAUMv1_12830004 | 1 | 1 | 100 | only missing in *Ca*.N. islandicus | |
| IBTHAUMO2_v1_210017 | | conserved exported protein of unknown function | 448 | **0.49%** | OG0000096 | IBTHAUMv1_1340002 | 1 | 1 | 100 | Core | |
| IBTHAUMO2_v1_320015 | | Alkyl hydroperoxide reductase | 182 | **0.48%** | OG0000941 | IBTHAUMv1_2320004 | 1 | 1 | 99.45 | only missing in *Ca*.N. salalaria | |
| IBTHAUMO2_v1_240129 | *fbp* | bifunctional fructose-1,6-bisphosphatase | 376 | **0.47%** | OG0000417 | IBTHAUMv1_310001 | 0.97606 | 1 | 99.46 | Core | |
| IBTHAUMO2_v1_1130058 | *pepA* | putative leucyl aminopeptidase | 477 | **0.46%** | OG0001191 | IBTHAUMv1_13940006 | 1 | 1 | 99.79 | missing in *Ca*. N. islandicus and all *Ca*. Nitrosotaleales | |
| IBTHAUMO2_v1_720005 | *rpoE* | DNA-directed RNA polymerase subunit E' | 193 | **0.46%** | OG0000408 | IBTHAUMv1_4130016 | 1 | 1 | 100 | Core | |

| | | | | | | | | | | | |
|---|---|---|---|---|---|---|---|---|---|---|---|
| IBTHAUMO2_v1_1130033 | *accC/ pccC* | acetyl-CoA/propionyl-CoA carboxylase, biotin carboxylase subunit | 485 | **0.45%** | OG0000362 | IBTHAUMv1_13940030 | 1 | 1 | 100 | Core | |
| IBTHAUMO2_v1_720031 | | NAD-binding D-isomer specific 2-hydroxyacid dehydrogenase | 310 | **0.43%** | OG0000385 | IBTHAUMv1_4660002 | 1 | 1 | 97.42 | Core | |
| IBTHAUMO2_v1_990024 | *fusA* | Elongation factor 2 | 730 | **0.42%** | OG0000645 | IBTHAUMv1_16760020 | 1 | 1 | 99.73 | Core | |
| IBTHAUMO2_v1_720043 | | exported protein of unknown function | 1462 | **0.42%** | OG0001466 | IBTHAUMv1_5230001 | 0.70588 | 0.99807 | 95.35 | *Ca.* N. bastadiensis unique | |
| IBTHAUMO2_v1_1100034 | | Peptidyl-prolyl cis-trans isomerase | 533 | **0.41%** | OG0000003 | IBTHAUMv1_17580009 | 1 | 1 | 99.25 | only missing in both Nitrosocosmicus spp. | |
| IBTHAUMO2_v1_40002 | | conserved protein of unknown function | 976 | **0.41%** | OG0000006 | IBTHAUMv1_3870008 | 0.61226 | 1.03381 | 75.22 | only missing in both Nitrosocosmicus spp. | s-layer protein family |
| IBTHAUMO2_v1_240132 | | conserved protein of unknown function | 217 | **0.41%** | OG0000436 | IBTHAUMv1_5220002 | 1 | 1 | 100 | Core | contains PF01865 domain - putative phosphate transport regulator |
| IBTHAUMO2_v1_1080008 | | exported protein of unknown function | 557 | **0.39%** | OG0001421 | IBTHAUMv1_15240001 | 0.33504 | 0.93896 | 62.33 | All 3 sponge symbionts, 2 *Ca*. Nitrosotaleales, 1 *Ca*. N. brevis | s-layer protein family |
| IBTHAUMO2_v1_240136 | | Short-chain dehydrogenase/reductase SDR | 574 | **0.38%** | OG0000810 | IBTHAUMv1_1110007 | 1 | 1 | 99.48 | Core | |
| IBTHAUMO2_v1_980023 | | membrane protein of unknown function | 586 | **0.38%** | OG0011727 | IBTHAUMv1_12910009 | 1 | 1 | 99.83 | singleton | |
| IBTHAUMO2_v1_1110030 | | 3-hydroxypropionyl-CoA dehydratase/Crotonyl-CoA hydratase [(S)-3-hydroxybutyryl-CoA forming] | 252 | **0.35%** | OG0000227 | IBTHAUMv1_18460003 | 1 | 1 | 100 | Core | |
| IBTHAUMO2_v1_510039 | *rrp42* | Exosome complex component Rrp42 | 270 | **0.33%** | OG0000834 | IBTHAUMv1_1960005 | 1 | 1 | 100 | Core | |
| IBTHAUMO2_v1_1100023 | *glyA* | Serine hydroxymethyltransferase | 448 | **0.29%** | OG0000541 | IBTHAUMv1_4580009 | 0.97545 | 1 | 100 | Core | |
| IBTHAUMO2_v1_940015 | *sucD* | succinyl-CoA synthetase, NAD(P)-binding, alpha subunit | 302 | **0.29%** | OG0000763 | IBTHAUMv1_330006 | 1 | 1 | 99.34 | Core | |
| IBTHAUMO2_v1_590109 | *sufD* | FeS assembly protein | 461 | **0.29%** | OG0000995 | IBTHAUMv1_4640007 | 1 | 1 | 100 | only missing in *Ca*.N. salalaria | |
| IBTHAUMO2_v1_800011 | | putative aldolase | 314 | **0.28%** | OG0000682 | IBTHAUMv1_14600002 | 1 | 1.0129 | 98.73 | Core | putatively functional in E.C. 4.1.2.13 |
| IBTHAUMO2_v1_590122 | *hemB* | Delta-aminolevulinic acid dehydratase | 321 | **0.27%** | OG0000183 | IBTHAUMv1_830005 | 0.85981 | 1 | 100 | Core | |
| IBTHAUMO2_v1_260012 | *trxB* | Thioredoxin reductase | 327 | **0.27%** | OG0000820 | IBTHAUMv1_4840006 | 0.93162 | 1 | 99.69 | Core | |
| IBTHAUMO2_v1_1080004 | | conserved exported protein of unknown function | 508 | **0.26%** | OG0000016 | IBTHAUMv1_16190003 | 1 | 1 | 99.41 | Core | |
| IBTHAUMO2_v1_170024 | | H(+)-transporting two-sector ATPase | 340 | **0.26%** | OG0000137 | IBTHAUMv1_2710003 | 1 | 1 | 100 | Core | |
| IBTHAUMO2_v1_950011 | *aroB* | 3-dehydroquinate synthase | 341 | **0.26%** | OG0000823 | IBTHAUMv1_560007 | 1 | 1 | 96.19 | Core | tyrosine, tryptophan, phe synthesis |
| IBTHAUMO2_v1_170028 | *atpI* | V-type ATP synthase subunit I | 694 | **0.25%** | OG0001210 | IBTHAUMv1_15570002 | 0.62104 | 1 | 100 | Core | |
| IBTHAUMO2_v1_1110031 | | 3-Hydroxypropionyl-CoA synthetase | 704 | **0.25%** | OG0000680 | IBTHAUMv1_18460004 | 1 | 1 | 100 | Core | |
| IBTHAUMO2_v1_770013 | | AAA family ATPase, CDC48 subfamily | 728 | **0.24%** | OG0000098 | IBTHAUMv1_11430014 | 1 | 1 | 100 | Core | |
| IBTHAUMO2_v1_300003 | | Redoxin domain-containing protein | 371 | **0.24%** | OG0000167 | IBTHAUMv1_13230016 | 1 | 1 | 99.73 | only missing in *Ca*.N. islandicus | |
| IBTHAUMO2_v1_470029 | | Beta-lactamase domain-containing protein | 421 | **0.21%** | OG0000769 | IBTHAUMv1_1130003 | 1 | 1 | 99.76 | Core | |
| IBTHAUMO2_v1_260042 | *hemL* | Glutamate-1-semialdehyde 2,1-aminomutase 2 | 441 | **0.20%** | OG0000389 | IBTHAUMv1_13740001 | 0.92517 | 0.99756 | 99.75 | Core | F430/tetrapyrrole/B12 biosynthesis |
| IBTHAUMO2_v1_980015 | *rpoB* | DNA-directed RNA polymerase subunit B | 1115 | **0.20%** | OG0000599 | IBTHAUMv1_1480002 | 0.71031 | 0.99874 | 99.87 | Core | |
| IBTHAUMO2_v1_370006 | *cysM* | Cysteine synthase | 492 | **0.18%** | OG0001268 | IBTHAUMv1_12200010 | 1 | 1 | 98.78 | missing in *Ca.* N. islandicus, both Nitrosocosmicus spp., all *Ca*. Nitrosotenuaceae | |

| | | | | | | | | | | | |
|---|---|---|---|---|---|---|---|---|---|---|---|
| IBTHAUMO2_v1_690011 | | conserved exported protein of unknown function | 492 | **0.18%** | OG0000003 | IBTHAUMv1_11050007 | 1 | 1 | 99.59 | not in both Nitrosocosmicus and many AOA have multiple copies | SSF51004: Cytochrome cd1-nitrite reductase-like, haem d1 domain superfamily |
| IBTHAUMO2_v1_320044 | *aco* | Aconitate hydratase | 745 | **0.18%** | OG0000268 | IBTHAUMv1_4090010 | 1 | 1 | 99.46 | Core | |
| IBTHAUMO2_v1_370005 | | von Willebrand factor type A | 498 | **0.18%** | OG0000623 | IBTHAUMv1_12200011 | 0.58635 | 1 | 99.66 | Core | COG4548: P, Nitric oxide reductase activation protein |
| IBTHAUMO2_v1_330024 | *pstS* | ABC-type phosphate transport system periplasmic component (PstS) | 508 | **0.17%** | OG0000113 | IBTHAUMv1_17840008 | 1 | 1 | 99.41 | missing in *Ca.* N. salaria, *Ca.*N. koreensis AR2, *Ca.* N. sediminis, and *Ca.* N. catalina | |
| IBTHAUMO2_v1_170026 | *amt2* | Ammonium transporter | 520 | **0.17%** | OG0000221 | IBTHAUMv1_15380013 | 0.96923 | 1 | 100 | missing in both Nitrosocosmicus spp. | |
| IBTHAUMO2_v1_620022 | | protein of unknown function | 1844 | **0.17%** | OG0001466 | IBTHAUMv1_12370012 | 0.85016 | 1 | 98.86 | *Ca.* N. bastadiensis unique | DUF5011 domain-containing |
| IBTHAUMO2_v1_960004 | *ths* | Thermosome subunit | 560 | **0.16%** | OG0002000 | IBTHAUMv1_4580008 | 0.89223 | 0.90179 | 42.57 | | |
| IBTHAUMO2_v1_530015 | *sdhA* | Succinate dehydrogenase or fumarate reductase, flavoprotein subunit | 570 | **0.15%** | OG0000437 | IBTHAUMv1_4000002 | 0.92632 | 1 | 98.48 | Core | |
| IBTHAUMO2_v1_590087 | | Type III restriction-modification system methyltransferase | 593 | **0.15%** | OG0001596 | IBTHAUMv1_9460007 | 0.95058 | 0.97302 | 31.37 | | |
| IBTHAUMO2_v1_240089 | *ppdK* | Pyruvate, phosphate dikinase | 884 | **0.10%** | OG0001061 | IBTHAUMv1_4310008 | 1 | 1 | 100 | | |
| IBTHAUMO2_v1_700001 | | protein of unknown function | 3094 | **0.10%** | OG0001466 | IBTHAUMv1_20360004 | 0.50517 | 0.96125 | 99.42 | *Ca.* N. bastadiensis unique | DUF5011 domain-containing |
| IBTHAUMO2_v1_880008 | | exported protein of unknown function | 2027 | **0.04%** | OG0000017 | IBTHAUMv1_19190004 | 0.32215 | 0.88243 | 46.86 | | putative M72 metalloendopeptidase |

*These values are ratios of alignment lengths computed for each comparison using the BLAST software :

minLrap = Lmatch/min(Lprot1, Lprot2)

maxLrap = Lmatch/max(Lprot1, Lprot2)

where Lmatch = length of the match, Lprot1 = length of protein 1, Lprot2 = length of protein 2

**if minLrap=1 and maxLrap=1** => the 2 proteins both align on their whole length

**if minLrap=1 and maxLrap<1** => one of the proteins is longer than the other, or the alignment is partial.

singletons refer to genes which did not have a hit above an expected threshold in OrthoFinder amongst all queried sequences

**If certain AOA do not have members in certain orthologous groups it does not mean that a more distant homologous gene is not present

**Table S4.** Comparison of net nitrite formation and nitrification rates from our study with previously published sponge studies.

| Sponge species | Net nitrification rates [µmol N (cm$^{-3}$ or g$^{-1}$ wet wt.) day$^{-1}$] | Net nitrite formation rates [µmol N (cm$^{-3}$ or g$^{-1}$ wet wt.) day$^{-1}$] | Marine area, depth, and season | Experimental Setup | AOB/AOA Diversity | Reference |
|---|---|---|---|---|---|---|
| ***Anthosigmella varians***<br>***Chondrilla nucula*** | 0.003 – 0.105 (g$^{-1}$*)<br>1.03 – 1.49 ◊ (g$^{-1}$*) | N.D. | Caribbean coral reef (6m) | 4h - 2.25 L batch incubations +/- light and 5 µM $NH_4^+$ | N.D. | Corredor *et al*., 1988 |
| ***Chondrilla nucula***<br>***Pseudaxinella zeai***<br>***Oligoceras violacea***<br>***Plakortis halichondroides*** | 0.864 – 6.36 (g$^{-1}$*)<br>0 – 2.47 (g$^{-1}$*)<br>0 – 1.37 (g$^{-1}$*)<br>0 – 0.768 (g$^{-1}$*) | 0.014 – 0.24 (g$^{-1}$*)<br>0 – 0.048 (g$^{-1}$*)<br>0.408 – 1.39 (g$^{-1}$*)<br>0 – 0.192 (g$^{-1}$*) | Caribbean coral reef (20-40m) and mangroves (1-3m); June-Sept. | 6 to 12h – 3 and 20L batch inc., +/- light | N.D. | Diaz & Ward, 1997 |
| ***Alpysina aerophoba***<br>***Dysidea avara***<br>***Chondrosia reniformis*** | 3.6 – 9.2 ◊ (g$^{-1}$)<br>0<br>~0.3 (g$^{-1}$) | N.D. | Med. Sea, 2-20m; | 9 to 28h - 3L batch, +/- 100 or 200 µM $NH_4^+$ | β-AOB 16S – 9 OTUs<br>– 1 OTU | Bayer *et al*., 2007 |
| ***Alpysina aerophoba*** | 0.214 – 0.826 (g$^{-1}$*) | N.D. | Med. Sea, 2-15m April - Sept. | 21 to 28h – 3L batch inc., +/- 100 or 200 µM $NH_4^+$ +/- nitrapyrin | A- and β-*amoA*/16S – 5, 7, 9 OTUs | Bayer *et al*., 2008 |
| ***Aplysina aerophoba***<br>***Agelas oroides***<br>***Dysidea avara***<br>***Chondroisa reniformis***<br>***Axinella polypoides***<br>***Ircinia oros*** | 1.13 (g$^{-1}$*)<br>0.875 (g$^{-1}$*)<br>N.S.<br>1.68 (g$^{-1}$*)<br>0.452 (g$^{-1}$*)<br>0.544 (g$^{-1}$*) | N.D. | Med. Sea, 10-20m; Sept. for all except for *A. oroides* (July) | 6h – 7L batch inc. | 6 A-*amoA* OTUs in 3 clust.<br>12 β-*amoA* OTUs in 3 clust.<br>83 γ-*amoA* clones | Jiménez & Ribes, 2007; Ribes *et al*., 2012 |
| ***Aplysina cauliformis***<br>***Smenospongia aurea***<br>**7 other species: *A. archeri, A. lacunose, I. felix, I. strobilina, P. crassa, V. rigidia*, and *X. muta*** | 4.08 (g$^{-1}$*)<br>4.32 (g$^{-1}$*)<br>+ + + + + + + | N.D. | Florida Keys, ~4m | 6 to 8h – 2-4L batches; *in situ* sampling | N.D. | Southwell *et al*., 2008 |
| ***Chondrosia reniformis*,**<br>***Dysidea avara*** | 0.176 (g$^{-1}$)<br>0.294 (g$^{-1}$) | 0.016 (g$^{-1}$)<br>0.016 (g$^{-1}$) | Med. Sea, 10-15m | 24h – 1L batches with 10 µM $NH_4^+$ | 1 A-*amoA* OTU | Schläppy *et al*., 2010 |
| ***Phakellia ventilabrum***<br>***Antho dichotoma***<br>***Geodia barretti*** | 0.14 – 2.26 (g$^{-1}$○)<br>0.<br>0.679 (g$^{-1}$○) | 0 – 0.192 (g$^{-1}$○)<br>0.360 (g$^{-1}$○)<br>0.180 (g$^{-1}$○) | Norwegian coast (fjords), 200-300m; Mar.-Nov. | 24 to 48 h in 500 to 900 mL batches with 10-12 µM $NH_4^+$ | 3 A-*amoA* OTUs<br>3 A-*amoA* OTUs<br>1 A-*amoA* OTU | Radax *et al*., 2012; Hoffmann *et al*., 2009 |
| ***Ianthella basta*** | 1.67 – 11.3 (g$^{-1}$)<br>1.54 – 13.3 ◊ (g$^{-1}$) | 0.64 – 6.07 (g$^{-1}$)<br>0.35 – 10.36 ◊ (g$^{-1}$) | Coral Sea, Australia, 10 m; Sept. – Oct. | 7- day long 24 h batch incubations in 1.74 L +/- 25, 100 µM $NH_4^+$ +/- PTIO | 1 A-*amoA* OTU | This study |

**(*)** signifies that sponge dry wt. g$^{-1}$ was converted by assuming dry weight = 10% of wet weight while a (○) signifies that sponge cm$^{-3}$ was converted by assuming 1 cm$^3$ sponge = 1.2 g wet weight sponge (Schläppy *et al*., 2010). A (◊) denotes that the rate is a potential rate (i.e. with added ammonium). A (+) indicates positive $NO_x^-$ production where quantity was not assessed. "A-, β- , and γ-*amoA*" refer to archaeal, betaproteobacterial and gammaproteobacterial *amoA* OTUs, respectively.

**Table S5**. AOA used for comparative genome analyses.

| Organism/Name | Assembly Accession | doi | Size (Mb) | GC% | Status | BioProject Accession | BioSample Accession | Strain | Completeness | Contamination |
|---|---|---|---|---|---|---|---|---|---|---|
| *Candidatus* Nitrosocaldus icelandicus | GCF_002906225.1 | doi: https://doi.org/10.1101/235028 | 1.62 | 41.5 | Complete Genon | PRJNA413650 | SAMN07759886 | 3F | 100 | 0 |
| *Nitrosopumilus maritimus* SCM1 | GCA_000018465.1 | doi: 10.1073/pnas.0913533107 | 1.65 | 34.2 | Complete Genon | PRJNA19265 | SAMN00000032 | SCM1 | 100 | 0.97 |
| *Candidatus* Cenarchaeum symbiosum A | GCA_000200715.1 | doi: 10.1073/pnas.0608549103 | 2.05 | 57.4 | Chromosome | PRJNA202 | SAMN02744041 | - | 99.03 | 0 |
| *Candidatus* Nitrosoarchaeum limnia SFB1 | GCA_000204585.1 | doi: 10.1371/journal.pone.0016626 | 1.77 | 32.6 | Chromosome | PRJNA52465 | SAMN02471010 | SFB1 | 98.06 | 0 |
| *Candidatus* Nitrosoarchaeum koreensis MY1 | GCA_000220175.2 | doi: 10.1128/JB.05717-11 | 1.61 | 32.7 | Contig | PRJNA67913 | SAMN02470178 | MY1 | 100 | 0 |
| *Candidatus* Nitrosopumilus salaria BD31 | GCA_000242875.3 | doi: 10.1128/JB.00013-12 | 1.57 | 33.8 | Contig | PRJNA50075 | SAMN00016669 | BD31 | 92.39 | 1.94 |
| *Candidatus* Nitrosopumilus koreensis AR1 | GCA_000299365.1 | doi: 10.1128/JB.01857-12 | 1.64 | 34.2 | Complete Genon | PRJNA174387 | SAMN02603137 | AR1 | 94.66 | 0 |
| *Candidatus* Nitrosopumilus sediminis | GCA_000299395.1 | doi: 10.1128/JB.01869-12 | 1.69 | 33.6 | Complete Genon | PRJNA174388 | SAMN02603138 | AR2 | 97.09 | 0 |
| *Candidatus* Nitrososphaera gargensis Ga9.2 | GCA_000303155.1 | doi: 10.1111/j.1462-2920.2012.02893.x | 2.83 | 48.3 | Complete Genon | PRJNA60505 | SAMN02603264 | - | 100 | 2.91 |
| *Candidatus* Nitrosotenuis chungbukensis | GCA_000685395.1 | doi: 10.1128/AEM.03730-13 | 1.76 | 41.8 | Contig | PRJNA210247 | SAMN02767256 | MY2 | 99.03 | 0.97 |
| *Nitrososphaera viennensis* EN76 | GCA_000698785.1 | doi: 10.1099/ijs.0.063172-0 | 2.53 | 52.7 | Complete Genon | PRJEA60103 | SAMN02721150 | EN76 | 100 | 0.97 |
| *Candidatus* Nitrosotenuis uzonensis N4 | GCA_000723185.1 | doi: 10.1371/journal.pone.0080835 | 1.64 | 42.2 | Contig | PRJEB4650 | SAMEA3139018 | N4 | 100 | 0.97 |
| *Candidatus* Nitrososphaera evergladensis SR1 | GCA_000730285.1 | doi: 10.1371/journal.pone.0101648 | 2.95 | 50.1 | Complete Genon | PRJNA235208 | SAMN03081530 | - | 100 | 2.91 |
| *Candidatus* Nitrosocosmicus oleophilus | GCA_000802205.2 | doi: 10.1111/1758-2229.12477 | 3.43 | 34.1 | Complete Genon | PRJNA210256 | SAMN03074222 | MY3 | 98.06 | 0.97 |
| *Candidatus* Nitrosopelagicus brevis | GCA_000812185.1 | doi: 10.1073/pnas.1416223112 | 1.23 | 33.2 | Complete Genon | PRJNA223412 | SAMN03273964 | CN25 | 99.51 | 0 |
| *Candidatus* Nitrosopumilus piranensis | GCA_000875775.1 | doi: 10.1038/ismej.2015.200 | 1.71 | 33.8 | Complete Genon | PRJNA269924 | SAMN03257648 | D3C | 100 | 0.97 |
| *Candidatus* Nitrosotenuis cloacae | GCA_000955905.3 | doi: 10.1038/srep23747 | 1.62 | 41.0 | Complete Genon | PRJNA272771 | SAMN03286947 | SAT1 | 100 | 1.94 |
| *Candidatus* Nitrosopumilus adriaticus | GCA_000956175.1 | doi: 10.1038/ismej.2015.200 | 1.80 | 33.4 | Complete Genon | PRJNA269341 | SAMN03253153 | NF5 | 100 | 0 |
| *Candidatus* Nitrosopumilus sp. Nsub (glass sponge MAG) | GCA_001541925.1 | doi: 10.1128/mSystems.00184-16 | 1.38 | 31.4 | Contig | PRJNA308059 | SAMN04386402 | Nsub | 100 | 0 |
| *Candidatus* Nitrosocosmicus exaquare | GCA_001870125.1 | doi: 10.1038/ismej.2016.192 | 2.99 | 33.9 | Complete Genon | PRJNA317395 | SAMN04606696 | G61 | 99.03 | 2.91 |
| *Candidatus* Nitrosomarinus catalina | GCA_002156965.1 | doi: 10.1111/1462-2920.13768 | 1.36 | 31.4 | Complete Genon | PRJNA341864 | SAMN05730076 | SPOT01 | 100 | 0 |
| *Candidatus* Nitrosotenuis aquarius AQ6f | GCA_002787055.1 | doi: 10.1128/AEM.01430-18 | 1.70 | 42.2 | Complete Genon | PRJNA406986 | SAMN07637255 | AQ6F | 99.68 | 1.94 |
| *Candidatus* Nitrosotalea devanaterra | GCA_900065925.1 | doi: 10.1128/AEM.04031-15 | 1.81 | 37.1 | Complete Genon | PRJEB10948 | SAMEA3577360 | - | 98.54 | 0 |
| *Candidatus* Nitrosotalea sinensis Nd2 | GCA_900143675.1 | doi: 10.1111/1462-2920.13971 | 1.60 | 37.4 | Contig | PRJEB15449 | SAMEA20449918 | - | 99.51 | 0.97 |
| *Candidatus* Nitrosotalea bavarica SbT1 | GCA_900167955.1 | doi: 10.1111/1462-2920.13971 | 1.55 | 36.0 | Scaffold | PRJEB15449 | SAMEA101071168 | - | 97.57 | 1.94 |
| *Candidatus* Nitrosotalea okcheonensis CS | GCA_900177045.1 | doi: 10.1111/1462-2920.13971 | 1.97 | 37.5 | Chromosome | PRJEB15449 | SAMEA20449168 | - | 99.51 | 0 |
| CcThau Ga0078905 (Sponge MAG) | IMG_2626541593_protein | doi:10.1038/ismej.2017.25 | 2.16 | 38.4 | Contig | NA | NA | - | 97.12 | 4.01 |
| *Candidatus* Nitrosospongia bastadiensis | NA | This publication | 1.99 | 64.8 | Scaffold | PRJEB29556 | SAMEA5126984 | O2 | 99.03 | 0.97 |
